# Implementation and acceleration of optimal control for systems biology

**DOI:** 10.1101/2021.03.17.435721

**Authors:** Jesse A Sharp, Kevin Burrage, Matthew J Simpson

## Abstract

Optimal control theory provides insight into complex resource allocation decisions. The forward-backward sweep method (FBSM) is an iterative technique commonly implemented to solve two-point boundary value problems (TPBVPs) arising from the application of Pontryagin’s Maximum Principle (PMP) in optimal control. In this review we discuss the PMP approach to optimal control and the implementation of the FBSM. By conceptualising the FBSM as a fixed point iteration process, we leverage and adapt existing acceleration techniques to improve its rate of convergence. We show that convergence improvement is attainable without prohibitively costly tuning of the acceleration techniques. Further, we demonstrate that these methods can induce convergence where the underlying FBSM fails to converge. All code used in this work to implement the FBSM and acceleration techniques is available on GitHub at https://github.com/Jesse-Sharp/Sharp2021.

## 1 Introduction

Across the life sciences, we encounter systems over which we wish to exert control. Whether we consider outbreak control in epidemiology [1,54], chemotherapy in oncology [8,19,73], muscle contraction and gait regulation in biomechanics [31,45,61], engineering cellular processes in synthetic biology [27,39], cell population growth in tissue engineering [24,75], or biodiversity and invasive species management in ecology [9,20,22], we face decisions around how a particular intervention should be applied to best achieve desired outcomes. Using mathematical models of such systems, optimal control theory provides insight into these resource allocation decisions.

Optimal control is a science of trade-offs; between competing objectives, or in weighing up the benefits of control measures against their costs. We illustrate some key concepts of optimal control in Figure 1. Suppose that without intervention, a crop yield will double, from *x*_0_ to 2*x*_0_, between now and harvest time. We might consider applying a control, such as fertiliser, to increase the growth rate of the crop; thereby increasing the yield at harvest to 3*x*_0_. Of course, applying fertiliser comes at a cost, and this must be considered against the increase in crop yield. As such, it is not immediately apparent how much fertiliser should be applied, and over what time period. This depends entirely on our characterisation of optimality; the *pay-off*. Depending on the pay-off, the optimal control may be continuous; whereby the strength can be readily and continuously adjusted throughout time, or bang-bang (discontinuous); whereby the control is applied at either a lower or upper bound with finitely many discrete switches between the two. The pay-off determines the objective(s) of control; which in our stylised example may be to maximise profits after cost of fertilising is considered, or achieve a specific yield, for example 3*x*_0_, using the minimum amount of fertiliser.

**Fig. 1.**
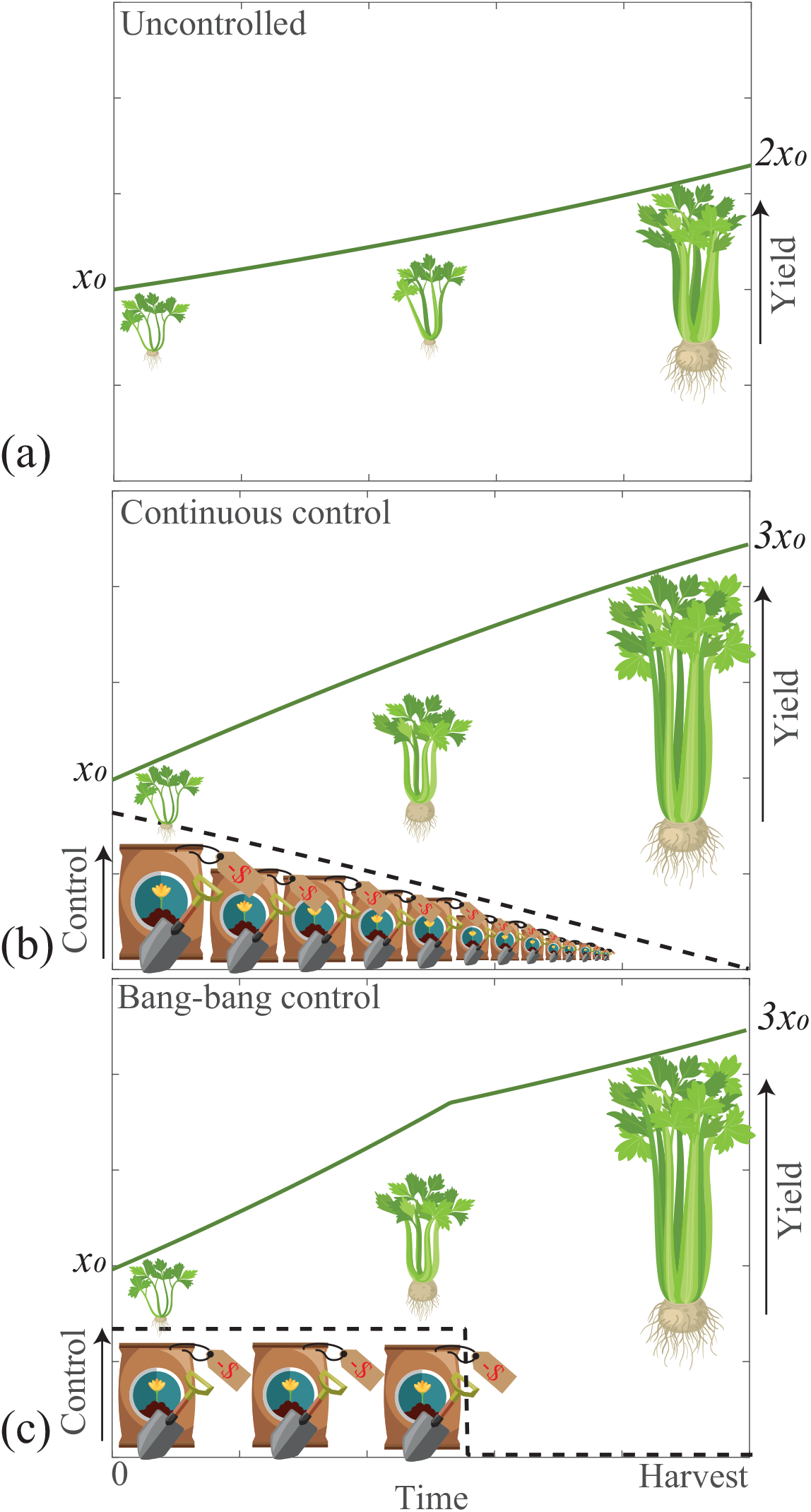
A pictorial example of optimal control for a growing crop. Suppose that initially, the crop yield is *x*_0_. We want to grow this crop to increase the yield, represented by the green line, come harvest time. Actions taken to increase the growth rate of the crop; such as applying fertiliser, are the controls, represented in black dash. Scenarios are presented for (a) no control, (b) continuous control, and (c) bang-bang control. Optimal control theory helps us determine how best to apply these controls. Illustrations adapted from ilyakalinin/iStock/Getty Images, johavel/iStock/Getty Images.

Much of modern day optimal control theory stems from the seminal works of Pontryagin; through the Pontryagin Maximum Principle (PMP) [64], and Bellman; through the advent of dynamic programming and the Hamilton-Jacobi-Bellman equation [11], in the 1950s and 1960s. These foundations of optimal control are built upon centuries of development in the calculus of variations [33]. For brief but broad expositions of the theoretical roots of optimal control and developments following these seminal works, we direct the reader to articles such as [17,71].

Often we are unable to solve optimal control problems analytically, so we pursue computational approaches. Broadly, the numerical methods for optimal control can be classed as either indirect or direct methods; for indirect methods optimality conditions are derived in the calculus of variations fashion via the PMP, leading to a two-point boundary value problem (TPBVP), while for direct methods the control problem is discretised and reformulated as a non-linear programming problem [67]. For an early history of numerical methods in optimal control, including gradient and conjugate gradient methods, Newton-Raphson methods, quasilinearisation, feasible direction algorithms and feed-back solutions we suggest [63]. Surveys [67,71] give an excellent overview of more recent developments in relevant numerical methods, including the forward-backward sweep method (FBSM), multiple-shooting methods, control parameterisation, collocation and pseudospectral methods and complete discretisation into finite-dimensional nonlinear programming problems.

The FBSM is an iterative method for solving the TPBVPs that arise from the indirect PMP approach to optimal control. FBSM has achieved popularity in optimal control, owing particularly to its straightforward scalability to large systems, and to its moderate computational cost and mathematical complexity [46]. In this work we review the implementation of the FBSM to solve optimal control problems, and investigate means of accelerating the convergence. To contextualise our discussion of the FBSM, we first consider the more familiar technique of successive over-relaxation (SOR). SOR is a generalisation of the Gauss–Seidel method, and is widely applied in numerical linear algebra to accelerate convergence when solving linear systems iteratively [80]. Essentially, the process of SOR involves specifying an acceleration or relaxation parameter, *β* ∈ (0, 2); a weighting factor that serves to reduce the spectral radius of the iterative matrix operator [69]. The error and rate of convergence of SOR is sensitive to this (problem dependent) choice of *β*, prompting investigation into theoretical convergence results and methods of determining *β* [23,44,69]. Despite challenges in identifying the optimal *β*, the SOR has historically been widely applied and studied in the literature due to the ease with which it can be implemented, and the rapid convergence it can deliver; even without identifying the optimal *β* [37,83].

This narrative closely parallels that of the FBSM in optimal control, where a weighting factor *ω* can be applied when updating the control between iterations to aid convergence. The optimal choice of *ω* is problem dependent, and significantly impacts the rate of convergence, or whether the FBSM converges at all. Nonetheless, the FBSM is frequently used in applied optimal control work as it is relatively straight-forward to implement, and can still converge in absence of the optimal *ω*. Theoretical convergence results of the FBSM are available in the literature [36,52], although the focus is on the FBSM without weighted updating, with no consideration for choosing *ω*. Using regularisation techniques, the FBSM is modified in [47] to improve convergence properties for large systems in a continuous setting, with a view to training deep neural networks in machine learning. These convergence results have recently been extended to the numerically discretised setting through symplectic Runge-Kutta discretisation; taking advantage of the variational structure of optimal control problems [48]. The authors also demonstrate that the rate of convergence of the regularised FBSM with symplectic discretisation can be improved with Anderson acceleration, an iterative acceleration technique. Although promising, this regularisation introduces a regularisation parameter, *ρ*. Similar to *ω*, the choice of *ρ* impacts convergence, and its choice is problem dependent. Understanding and implementing the regularisation and symplectic techniques is not trivial, and introduces conceptual complexity beyond what is necessary for many applied optimal control problems. As such, the standard FBSM remains an attractive choice for practitioners.

To this end, we aim to review acceleration techniques that can be paired with the standard FBSM. We implement such techniques alongside the FBSM with the goals of: (1) increasing the rate and frequency of convergence, and (2) reducing the importance of, and challenges associated with, selecting *ω*. A graphical overview of the optimal control process we employ in this work, including the incorporation of acceleration methods, is presented in Figure 2. We note that all code used to implement the algorithms presented in this review; the FBSM and the Wegstein, Aitken-Steffensen and Anderson acceleration methods, is available on GitHub.

**Fig. 2.**
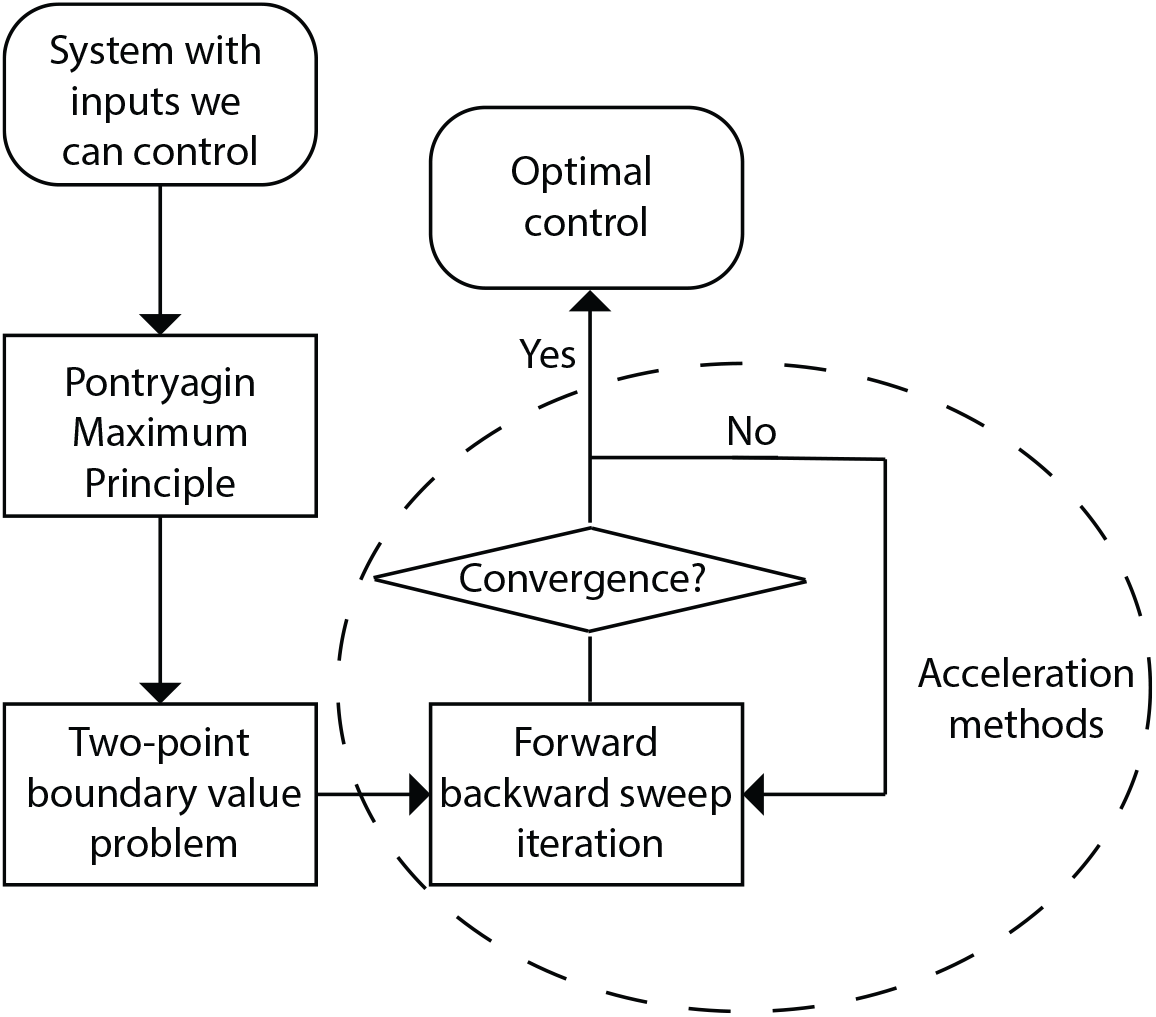
The process of optimal control via the Pontryagin Maximum Principle approach, with the incorporation of acceleration methods.

Throughout this work we consider optimal control in the systems biology context. However, we note that optimal control is relevant to a wide variety of fields including chemical engineering [49], aeronautics and astronautics [67], management science and economics [72]. The FBSM, and by extension, the acceleration techniques we consider in this work, can be readily applied in any of these areas.

In §2 we review the PMP approach to optimal control, and the implementation of the FBSM. We provide a single-variable linear model, and a multi-variable nonlinear model in §3; and pose and solve example continuous, bang-bang (discontinuous), and fixed endpoint control problems. We review potential iterative acceleration methods in §4, and present the results of selected techniques in §5. We discuss the performance of these techniques in §6, and identify opportunities for application and further investigation.

## 2 Forward-backward sweep method

In an optimal control problem with one state variable, *x*(*t*), one control, *u*(*t*), over a fixed time interval, *t* ∈ [*t*_0_*, t_N_*]; such as the crop growth example presented in Figure 1, we seek the optimal control *u*\*(*t*) that minimises or maximises a specified pay-off function, *J*, subject to the dynamics of the state. In this section we briefly review the Pontryagin Maximum Principle approach to such an optimal control problem, and the standard implementation of the FBSM for solving the resulting two-point boundary value problem. The FBSM is readily extended to problems with multiple state variables, multiple controls, state constraints and free end-times [46,52,73,74], however for this overview we restrict ourselves to the single variable, single control, fixed end-time case for clarity.

The pay-off typically comprises a cost function 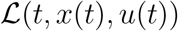 integrated over the time interval, and/or a function of the final state, *x*(*t_N_*). As such, we seek to minimise or maximise *J*, subject to

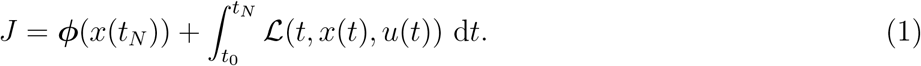

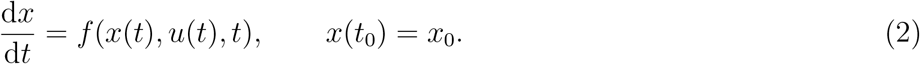

Applying the PMP, we construct the Hamiltonian; 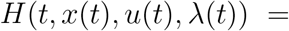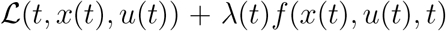, where *λ*(*t*) is the co-state variable linking our state to our pay-off. The necessary conditions for optimal control are obtained from the Hamiltonian:

1. The optimal control, *u*\* (*t*), is obtained by minimising the Hamiltonian,

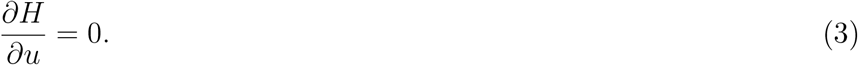
2. The co-state is found by setting,

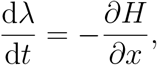
3. satisfying the transversality condition,

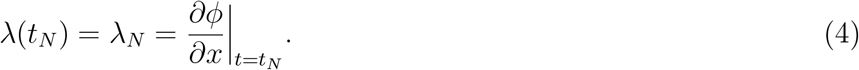

Following these steps yields a TPBVP to solve for *x*(*t*), *λ*(*t*), subject to *x*(*t*_0_) = *x*_0_, and *λ*(*t_N_*) = *λ_N_*. To solve this numerically, we discretise *t* into *N* + 1 time points separated by a step-size d*t* = (*t_N_* − *t*_0_)*/N*; **t**= [*t*_0_*, t*_0_ + d*t,…, t*_0_ + *N*d*t*] = [*t*_0_*, t*_1_*,…, t_N_*]. Here, we consider a uniform discretisation in time; although this is not strictly necessary, as discussed in §3. Using superscripts to denote the iteration number, provide an initial guess of the control at each 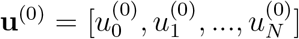. From **u**(0), solve Equation (2) numerically from *t*_0_ to *t_N_* to obtain 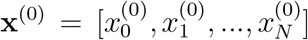. Now, using **x**(0), solve for 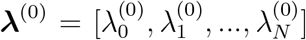 backwards in time from *t_N_* to *t*_0_, starting from *λ_N_*. With the optimality condition from Equation (3), generate a temporary update for the control, 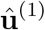. The next iteration begins with an updated guess for the control, **u**(1). These steps are repeated until a convergence condition is satisfied. The algorithm for the FBSM is summarised in §1 of the supplementary material.

In some instances, directly updating the control, such that

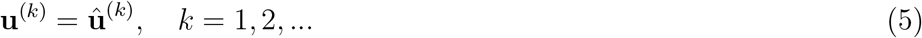

is sufficient, however more commonly a weighted update is performed [46,73], such that in the (*k* + 1)th iteration,

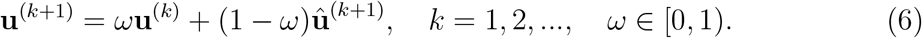

This weighted updating is also referred to as applying a relaxation factor, similar to SOR as discussed in §1. An appropriate choice of *ω* in Equation (6) can accelerate convergence relative to Equation (5), or in some cases induce convergence where Equation (5) leads to divergence. The weighting parameter, *ω*, can be held constant between iterations, although faster convergence may by achieved by updating *ω*. For example, by reducing *ω* as the system approaches convergence, a greater portion of the updated control is maintained relative to the control from the previous iteration [46], possibly accelerating convergence. A challenge commonly faced in implementing this control updating scheme is that the best choice for *ω* is problem dependent, and often is determined heuristically in practice. We address the extent to which the proposed acceleration algorithms address this issue in §4.

To facilitate the following discussion regarding acceleration, we note that the FBSM can be thought of as a generalised fixed point iteration [52], where each iteration comprises a forward and backward sweep and a control update. As such, for a control problem with one control, discretised into *N* + 1 time points, each iteration of the FBSM can be thought of as the application of a nonlinear operator, 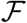, of dimension *N* + 1, such that 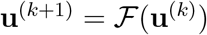, or:

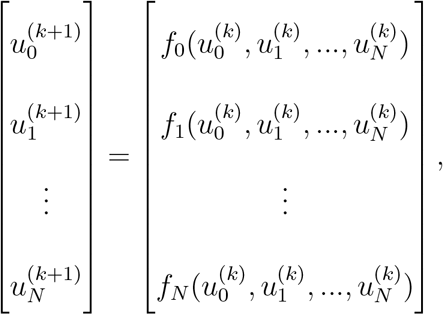

where 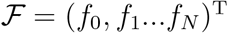. However, in general we are not able to write down an explicit expression for 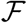. Conceptualising the FBSM as a fixed point iteration process informs the choice of acceleration methods discussed in §4.

Importantly, we use the term *function evaluation* in this work to refer to the process of solving the system of ODEs for the state forward in time and the system of ODEs for the co-state backwards in time, once. This aligns with a single iteration of the standard FBSM. The function evaluation nomenclature becomes convenient when discussing the FBSM in the context of acceleration algorithms, that typically focus on reducing the number of times expensive functions are evaluated. Producing numerical solutions to the ODE systems is by far the most computationally expensive component of the FBSM. This computational expense increases with the size and complexity of the systems; reducing the number of times these systems must be solved becomes more advantageous as the size and complexity of the systems increases. The function evaluation description also facilitates comparison between acceleration methods that require solving the ODE systems a different number of times per iteration. Throughout this work, we use 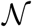 to denote the total number of function evaluations a given method takes to achieve convergence.

### 2.1 Adapted forward-backward sweep method

The FBSM can be extended to handle problems where we aim not only to minimise or maximise a given quantity over time, but also ensure that a specific state is reached at final time. This aligns with the crop growth example from Figure 1 if the objective is to achieve a specific yield of 3*x*_0_ at harvest, rather than to maximise yield. In this case we may have an integral term in the pay-off as described in Equation (1), however the function of the final state, ****ϕ****(*x*(*t_N_*)), is redundant in a control problem with a prescribed final state. Equation (2) is also modified to incorporate the additional constraint:

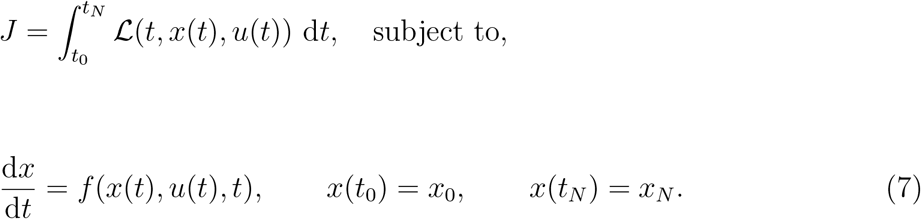

Here, *x_N_* is the specified state that must be reached at final time. Since we have introduced an additional boundary value to the system, we no longer obtain the transversality condition from Equation (4). Instead, we seek the final time condition on the co-state, *λ_N_*, and associated optimal control that satisfies Equation (7). We proceed by considering an adapted FBSM that takes as an input a guess for this final time condition, 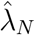, and solves the corresponding control problem. If we denote this application of the FBSM as the function 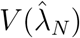, and the corresponding final value of the state, 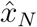, then the adapted FBSM is an iterative process that solves for the root of 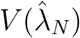; the value of 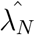 for which 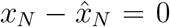. This outer iterative process can be solved using standard techniques such as the bisection method or secant method; the former converging more reliably provided that the initial guesses for 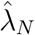 form an interval that brackets the root, the latter converging in fewer iterations [46]. Each of these outer iterations necessitates solving a boundary value problem to convergence, often involving numerous iterations of the FBSM. In this work we apply the secant algorithm as presented in [46] without modification, for the adapted FBSM. The acceleration techniques described in §4 are applied only to the inner FBSM processes, reducing 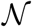 for each internal FBSM problem, leaving the outer secant iterations unchanged. Using 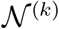 to denote the number of function evaluations in the *k*th internal FBSM problem, we can express the cumulative function evaluations required for convergence of the adapted FBSM as Σ, such that 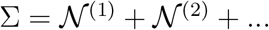.

## 3 Control problems

To investigate the robustness and effectiveness of the iterative acceleration techniques that we will discuss in §4, we consider two distinct systems, and for each system we study three example control problems. The first system is a single species linear differential equation subject to a control. We later demonstrate that under certain conditions we are able to obtain exact solutions for control problems applied to this model. The second system is a three species model for acute myeloid leukaemia (AML) governed by a coupled nonlinear system of differential equations, subject to a control. We construct the linear model to examine the behaviour of the acceleration techniques as applied to a simple idealised set of control problems. We include the AML model, variations upon which have been considered in the literature [26,73,74], to examine how the acceleration techniques perform when applied to problems more reflective of those considered in applied optimal control. For each model, we consider three distinct control problems; continuous control, bang-bang control and continuous control with fixed endpoint.

For all control results presented in this work, convergence is deemed to be achieved when the error, *ε*, measured as the Euclidean norm of the difference between subsequent controls, falls below a tolerance of 1 × 10^−10^. Numerical solutions to ODEs are obtained using a fourth-order Runge-Kutta method [65] with constant time-stepping. A uniform time discretisation is sufficient for all control problems considered in this work. However, the FBSM and acceleration methods readily generalise to a non-uniform discretisation. If the desired discretisation for the state equations differs from that of the co-state equations, it is necessary to perform interpolation within each iteration of the FBSM to obtain values at corresponding time points. This can be computationally expensive and introduce an additional source of error. Where the desired discretisations for the state and co-state differ, numerical schemes with internal interpolation such as Continuous Runge-Kutta methods may be appropriate [10,84].

### 3.1 Single-variable linear model

The linear model is a single species model for the growth of *x*(*t*), subject to control *u*(*t*) that increases the growth rate. This model could represent our stylised crop growth example presented in §1. We suppress the explicit time dependence of the state and co-state variables and the control in the following equations for notational convenience. For numerical results, we solve the linear problems on the domain 0 ≤ *t* ≤ 1, with time-step d*t* = 3.91 × 10^−3^, giving *N* = 257 time points.

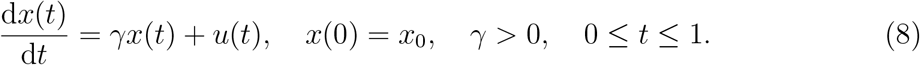

In absence of control, *u*(*t*) ≡ 0, this model admits the solution *x*(*t*) = *x*_0_e*^γt^*, describing exponential growth.

#### 3.1.1 Continuous control

We seek to maximise a quadratic cost function *J*, subject to

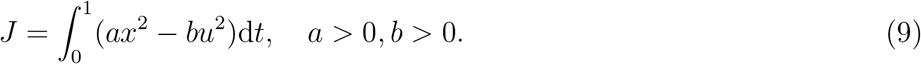

Following the standard Pontryagin Maximum Principle approach for solving optimal control problems, we form the Hamiltonian and derive the co-state equation, transversality condition and optimality condition. The Hamiltonian is given by

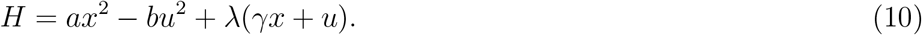

The co-state equation is

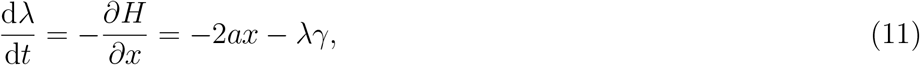

with transversality condition *λ*(1) = 0. In this case the optimality condition is

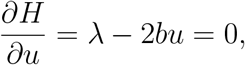

such that the optimal control is given by

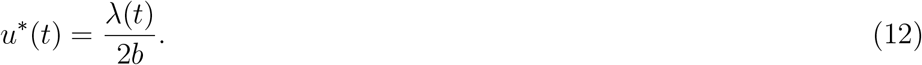

For model parameter *γ* = 0.5 and pay-off weightings *a* = *b* = 1, with initial condition *x*_0_ = 1, we are able to solve the control problem analytically using standard techniques for linear systems with complex eigenvalues [42]. The process is laborious so we present the approach and analytical solution in §2 of the supplementary material. In the supplementary material we also plot the analytical results against the numerical results to demonstrate the excellent agreement. The numerical solution to the linear continuous control problem is presented in Figure 3. Convergence via the FBSM requires 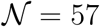 iterations.

**Fig. 3.**
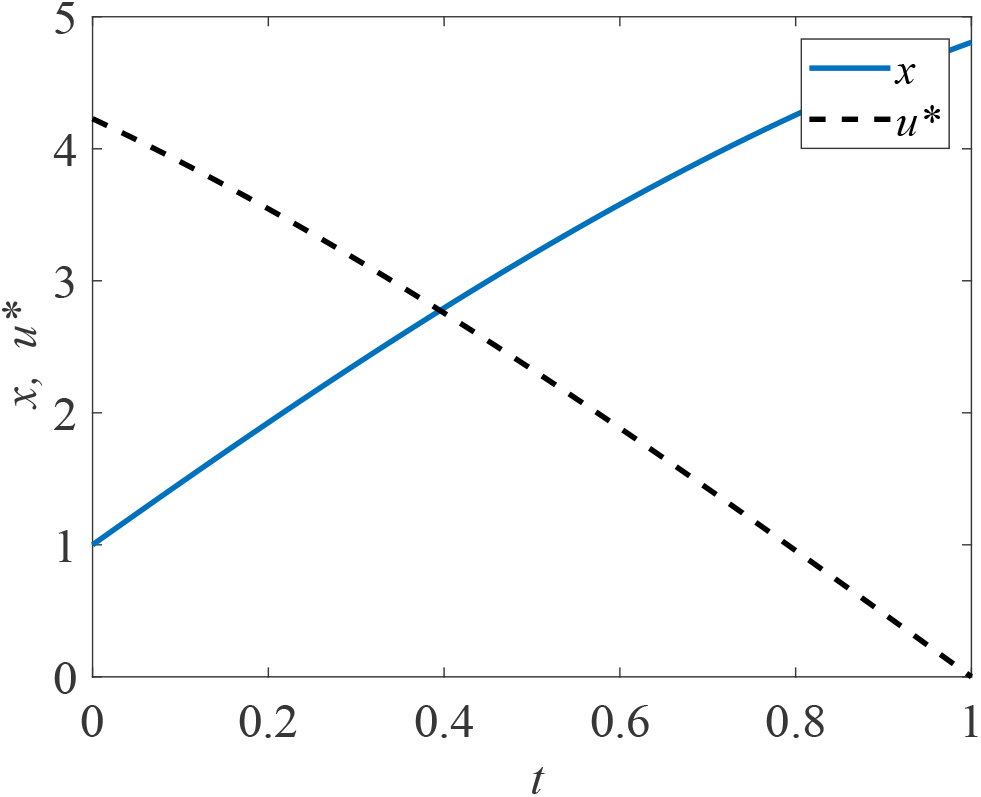
Solution to the linear continuous control problem. The optimal control, *u*\*(*t*), is shown in black dash and the corresponding state, *x*(*t*), in blue. This solution is produced with model parameter *γ* = 0.5, time-step d*t* = 3.91 × 10^−3^, over the interval 0 ≤ *t* ≤ 1. The contributions of the state and the control to the pay-off are equally weighted, with *a* = *b* = 1.

#### 3.1.2 Bang-bang control

For the bang-bang control we consider the same state equation as in Equation (8), and incorporate bounds on the control.

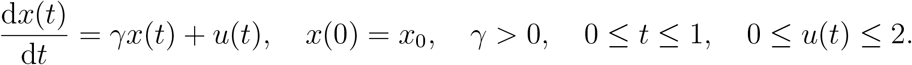

We seek to maximise a cost function *J* that is linear in *u*,

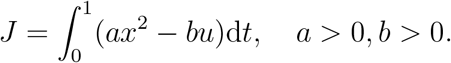

We form the Hamiltonian and derive the co-state equation and transversality condition:

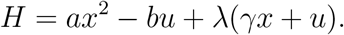

The co-state equation is

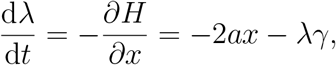

with transversality condition *λ*(1) = 0.

In seeking the optimality condition we find

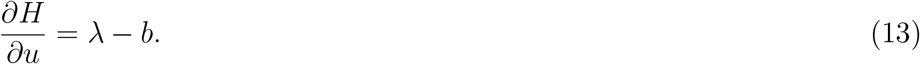

As Equation (13) does not depend on *u*, we define a switching function

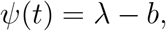

and produce an expression for the control, based on the bounds on *u* and the sign of the switching function:

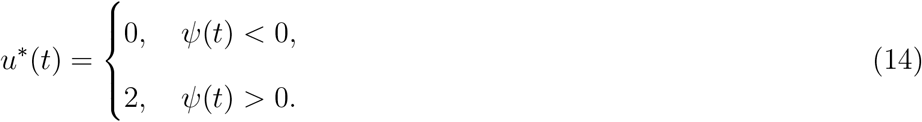

If *ψ*(*t*) is zero over any finite interval excluding isolated points, the optimal control is singular rather than bang-bang. Over such intervals, minimisation of the Hamiltonian does not provide sufficient information to determine the optimal control, and further conditions must be considered. [16,46]. We restrict our focus in this work to non-singular bang-bang optimal control problems. The numerical solution to the linear bang-bang control problem is presented in Figure 4. Convergence to this solution via the FBSM required 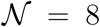 iterations.

**Fig. 4.**
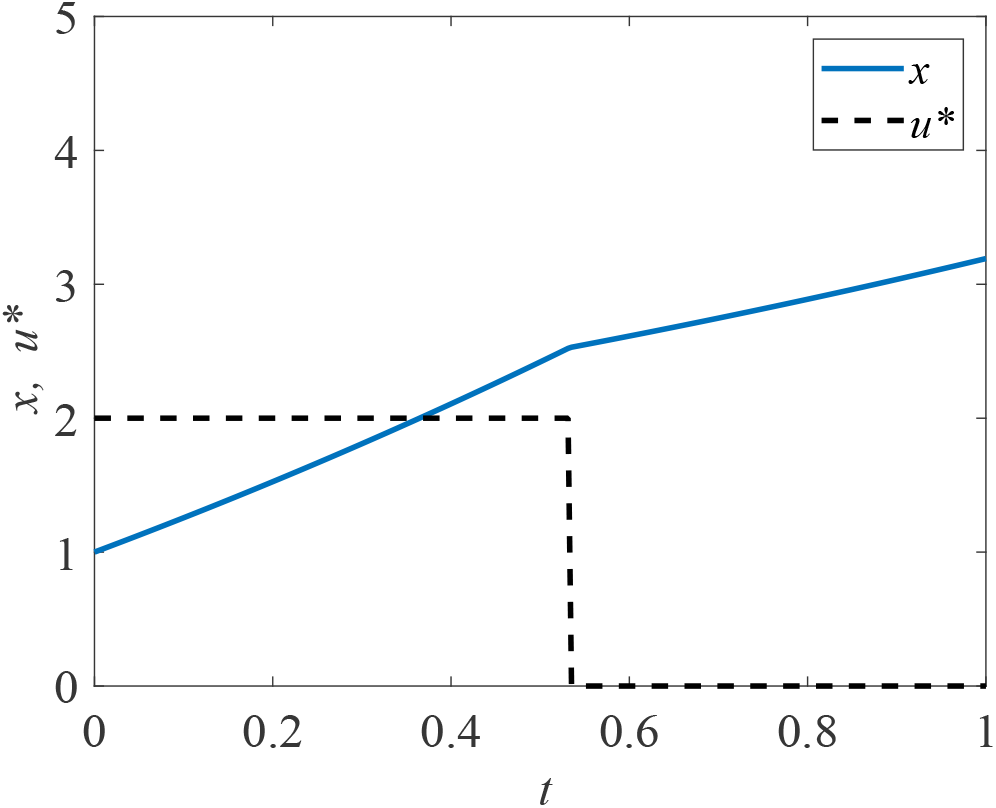
Solution to the linear bang-bang control problem. The optimal control, *u*\*(*t*), is shown in black dash and the corresponding state, *x*(*t*), in blue. This solution is produced with model parameter *γ* = 0.5, time-step d*t* = 3.91 × 10^−3^, over the interval 0 ≤ *t* ≤ 1, with pay-off weightings of *a* = 1 for the state, and *b* = 3 for the control. The bang-bang control has prescribed bounds of 0 ≤ *u*\*(*t*) ≤ 2.

#### 3.1.3 Continuous control with fixed endpoint

For the fixed endpoint problem, we proceed with the same state equation, however we now impose a terminal condition on *x*.

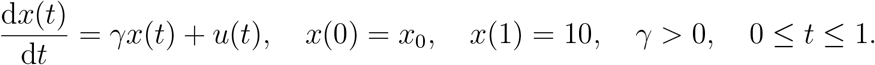

We seek to maximise the same quadratic cost function *J*, as considered in Equation (9). As such, we form the same Hamiltonian given in Equation (10) and derive the same co-state, Equation (11), and expression for the control, Equation (12). Note however that we do not prescribe a final time condition on the co-state equation via the transversality condition; as the system already has two boundary conditions, doing so would cause it to be overdetermined. Instead, we make two guesses for *λ*(1); for example, *λ*^(0)^(1) = −10 and *λ*^(1)^(1) = 10. We proceed by applying the adapted FBSM outlined in §2, using these guesses to initialise the secant method. Numerical results for the linear fixed endpoint control problem are presented in Figure 5. Convergence of the adapted FBSM is achieved after Σ = 177 iterations.

**Fig. 5.**
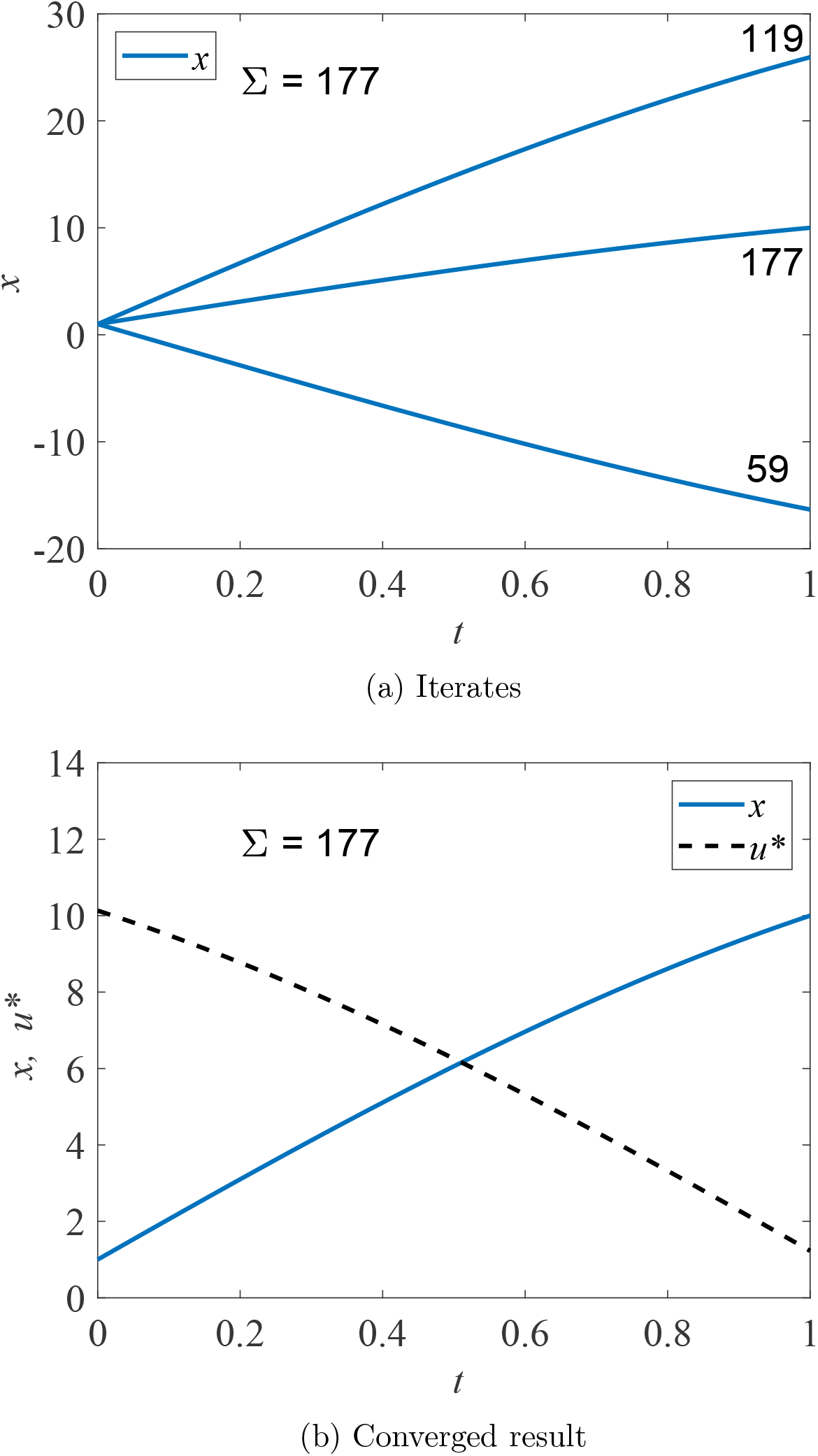
Results are presented for the linear problem with specified terminal state value, *x*(*t_N_*) = 10, solved using the adapted FBSM. Underlying FBSM problems are solved with time-step d*t* = 3.91 × 10^−3^, over the interval 0 ≤ *t* ≤ 1, with pay-off weightings of *a* = *b* = 1. In (a) the *x*(*t*) iterates of the adapted FBSM are presented. We annotate the cumulative function evaluations after the first 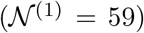 and second 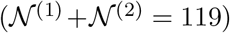 iterations of the adapted FBSM, based on initial guesses for *λ*(*t_N_*) of *λ*(*t_N_*) = −10 and *λ*(*t_N_*) = 10. The total cumulative function evaluations required for convergence of the adapted FBSM, 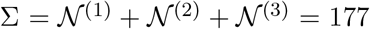, is indicated. The converged result for *x*(*t*), satisfying |*x*(*t_N_*) − 10| ≤ 1 × 10^−10^ is presented in (b); this figure also includes the optimal control, *u*\*(*t*).

### 3.2 Multiple-variable nonlinear model

The AML model is a nonlinear coupled multi-species model describing the interactions between progenitor blood cells, *A*(*t*), and leukaemic stem cells, *L*(*t*), that occupy the same niche in the bone marrow; thereby competing for space and resources. Haematopoietic stem cells, *S*(*t*), act as upstream production of *A*(*t*). These dynamics have been explored in the literature both experimentally [41,76], and through mathematical modelling [26,50]. We subject the model to a chemotherapy-like control, *u*(*t*), that acts as an additional death term for *L*(*t*). The state can be expressed as ****x****(*t*) = [*S*(*t*)*, A*(*t*)*, L*(*t*)]*^T^*. As there are now three state equations, we require three co-state equations; ****λ****(*t*) = [*λ*_1_(*t*)*, λ*_2_(*t*)*, λ*_3_(*t*)]*^T^*. We suppress the explicit time dependence of the state and co-state variables and the control in the following equations for notational convenience:

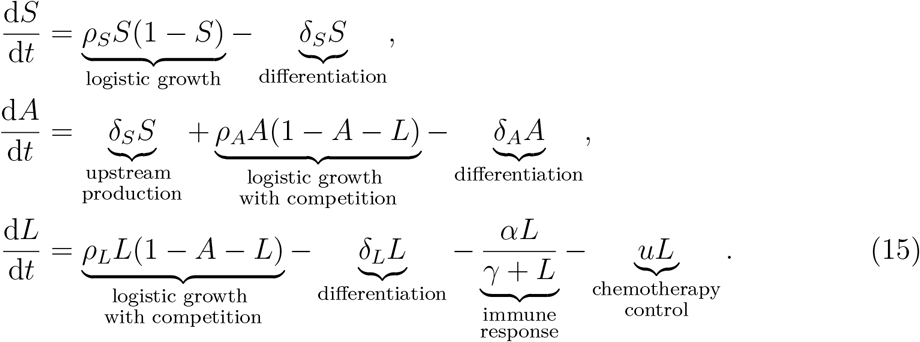

For each control problem associated with the AML model, we use initial conditions that yield a coexisting steady state in absence of control (all three species non-zero); *S*(0) = 1 − *δ_S_/ρ_S_*, *A*(0) = 0.3255, and *L*(0) = 0.3715. We solve the AML problems numerically on the domain 0 ≤ *t* ≤ 10, with time-step d*t* = 4.88 × 10^−4^, giving *N* = 20481 time points. Model parameters are specified in Table 1.

**Table 1:**
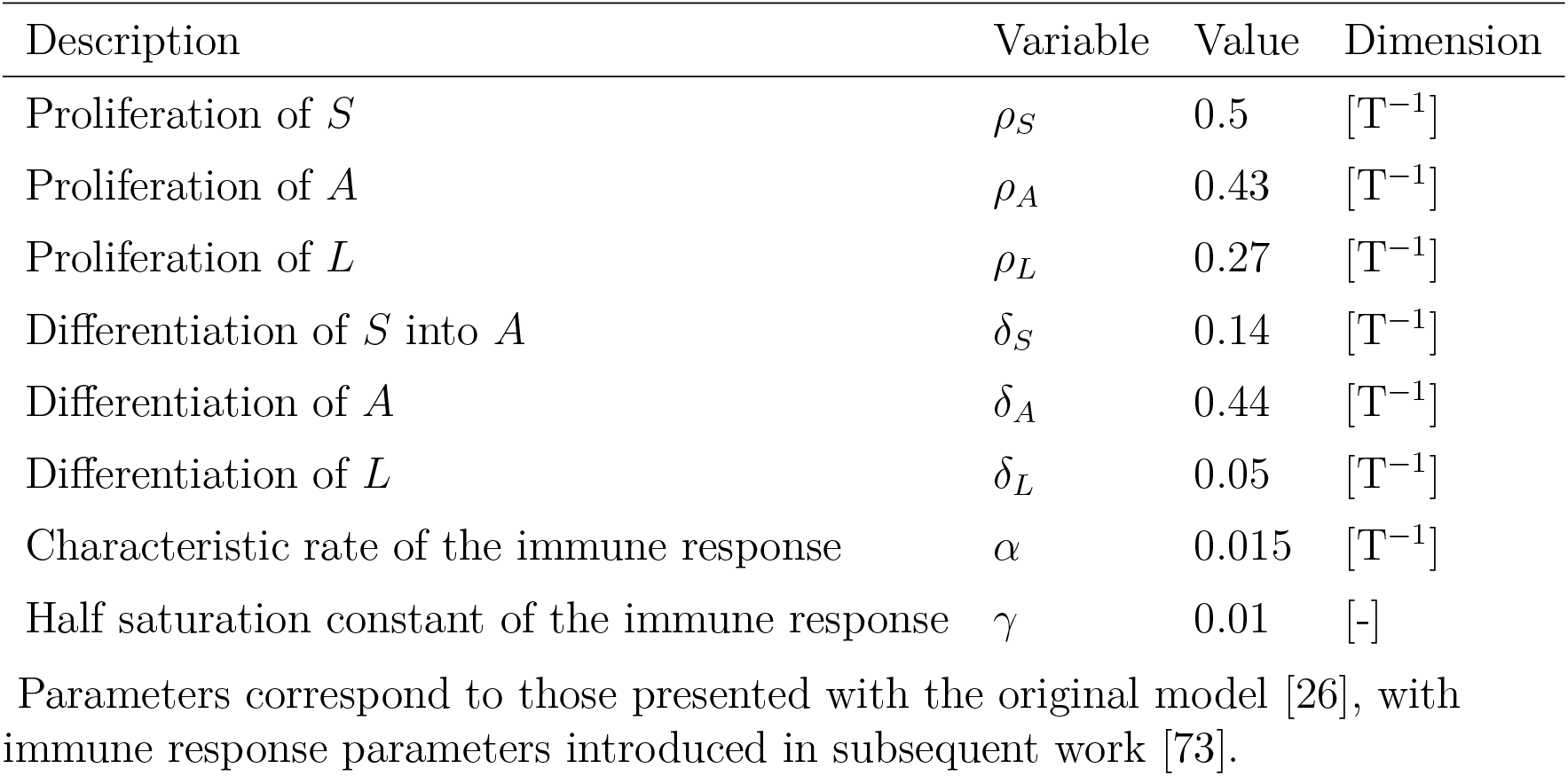
AML model parameters

#### 3.2.1 Continuous control

For the AML continuous control problem we seek to minimise a quadratic cost function *J*, that accounts for both the cost of applying the control and the cost of the leukaemic burden, subject to

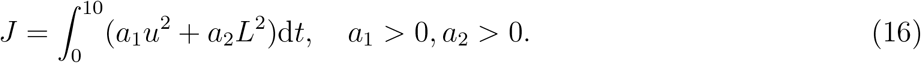

We form the Hamiltonian and derive the co-state equation, transversality condition and optimality condition. The Hamiltonian is given by

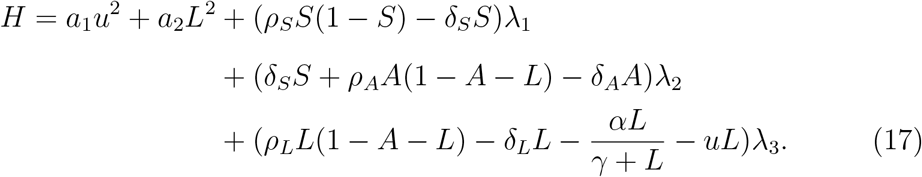

The co-state equations are

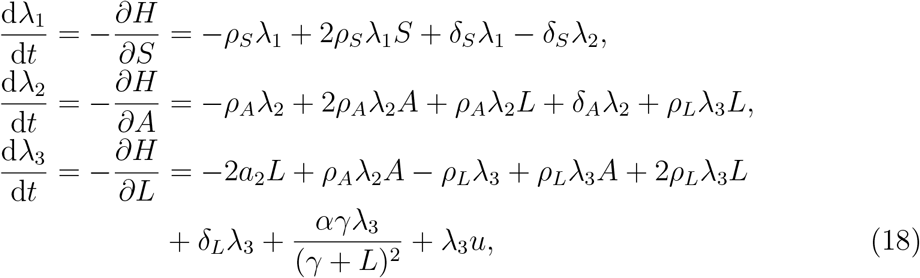

with transversality conditions *λ*_1_(10) = *λ*_2_(10) = *λ*_3_(10) = 0, obtained in the usual way. In this case the optimality condition is

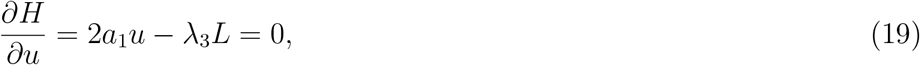

such that the optimal control is given by

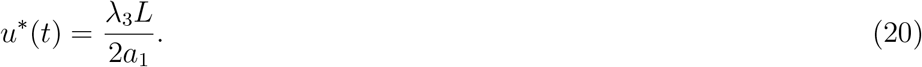

Numerical solutions for the AML continuous control problem are presented in Figure 6. These solutions are obtained via the FBSM, requiring 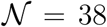 iterations with *ω* = 0.55. This choice of *ω* minimises 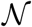 for the AML continuous control problem solved with the FBSM without acceleration techniques. We discuss the choice of *ω* further in §5.

**Fig. 6.**
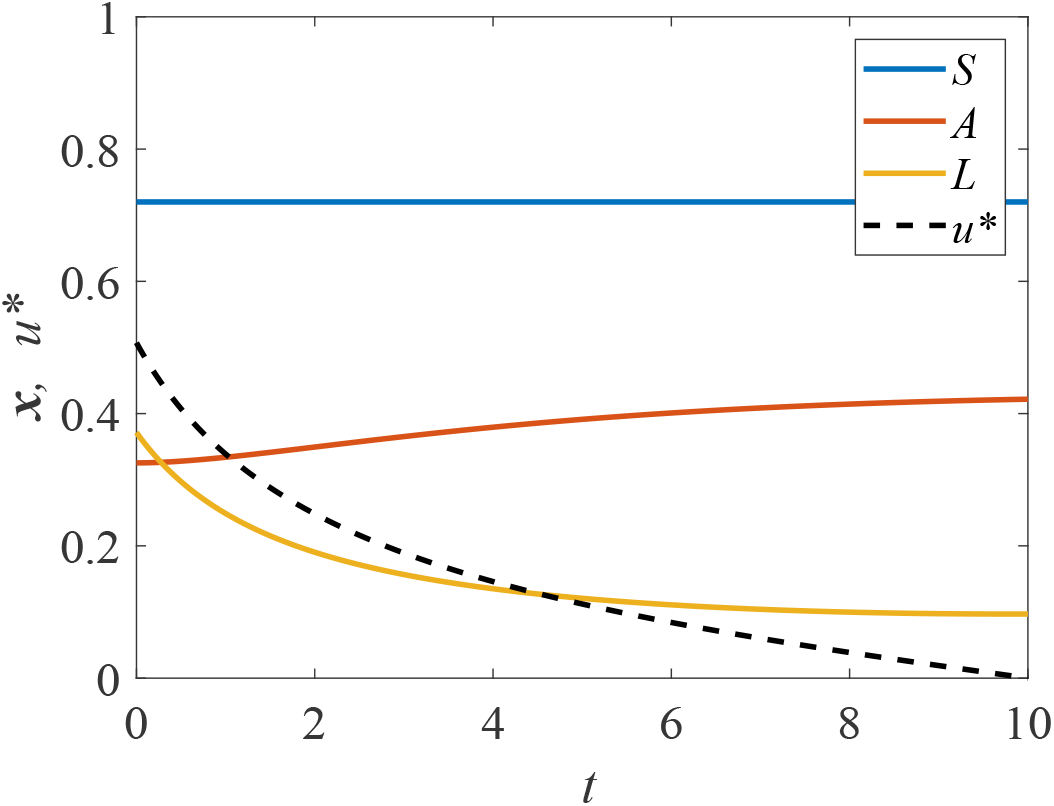
Solution to the AML continuous control problem. The optimal control, *u*\*(*t*), is shown in black dash and the corresponding state equations for *S*(*t*), *A*(*t*) and *L*(*t*) are shown in blue, red and yellow, respectively. This solution is produced with model parameters given in Table 1, time-step d*t* = 4.88 × 10^−4^, over the interval 0 ≤ *t* ≤ 10, with pay-off weightings of *a*_1_ = 1 for the control, and *a*_2_ = 2 for the state variable *L*(*t*).

#### 3.2.2 Bang-bang control

For the bang-bang AML problem we consider the same states as in Equation (15), and incorporate bounds, 0 ≤ *u* ≤ 0.3, on the control. We seek to minimise a cost function *J* that is linear in the control and the state variable *L*:

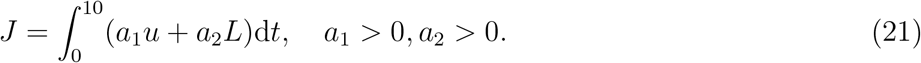

We form the Hamiltonian and derive the co-state equations, transversality conditions and optimality condition. The Hamiltonian is given by

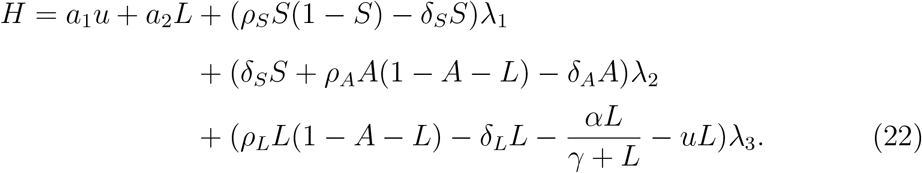

The co-state equations are

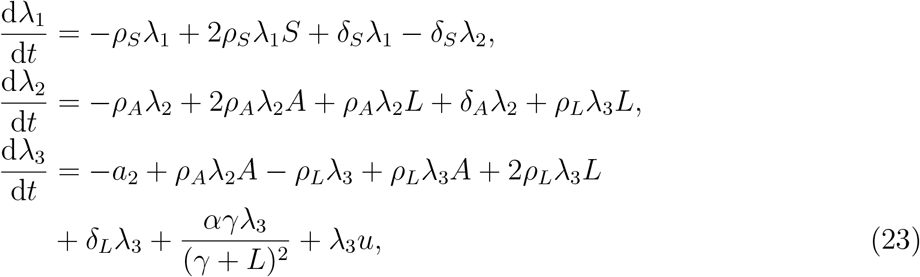

with transversality conditions *λ*_1_(10) = *λ*_2_(10) = *λ*_3_(10) = 0. In this case the switching function is

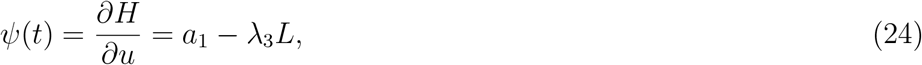

such that the optimal control is given by

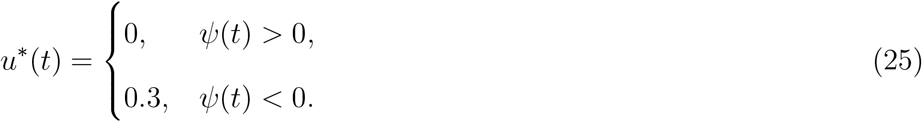

Note that the correspondence between the sign of *ψ*(*t*) and the chosen bound is reversed in Equation (25) relative to Equation (14) as we are now performing minimisation rather than maximisation. Numerical solutions for the AML bang-bang control problem are presented in Figure 7. These solutions are obtained via the FBSM, requiring 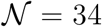 iterations with *ω* = 0.4. This choice of *ω* minimises 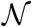 for the AML bang-bang control problem solved with the FBSM without acceleration techniques. We discuss the choice of *ω* further in §5.

**Fig. 7.**
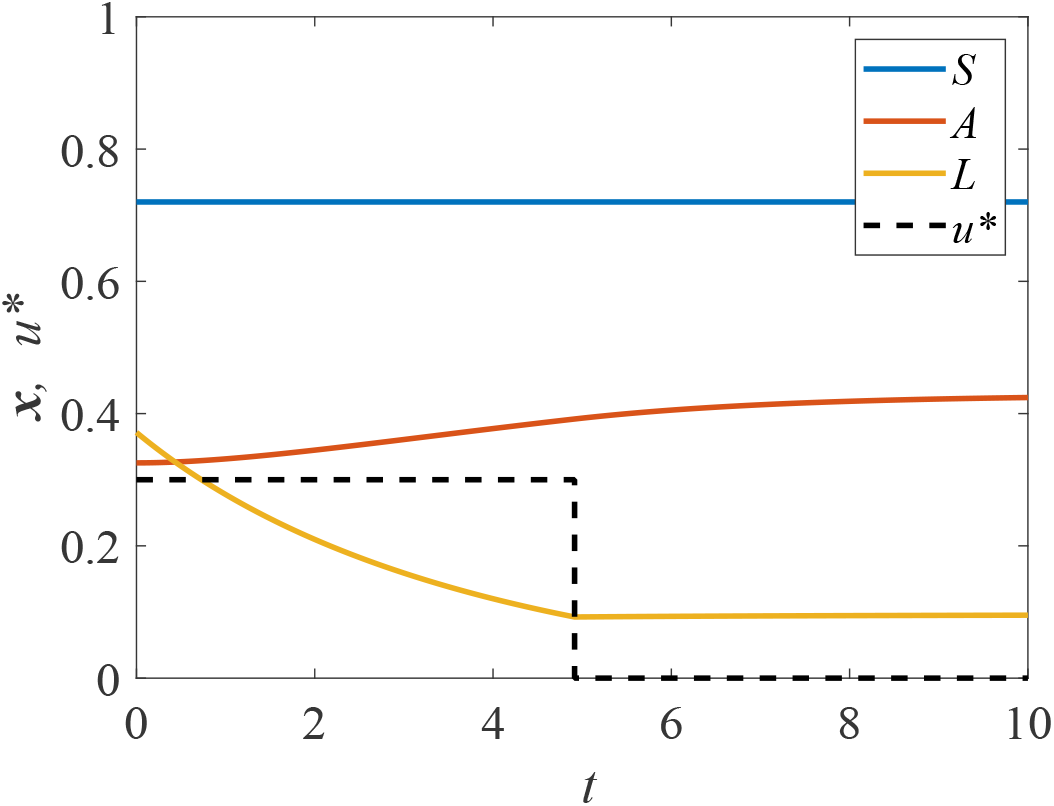
Solution to the AML bang-bang control problem. The optimal control, *u*\*(*t*), is shown in black dash and the corresponding state equations for *S*(*t*), *A*(*t*) and *L*(*t*) are shown in blue, red and yellow, respectively. This solution is produced with model parameters given in Table 1, time-step d*t* = 4.88 × 10^−4^, over the interval 0 ≤ *t* ≤10, with pay-off weightings of *a*_1_ = 1 for control, and *a*_2_ = 2 for the state variable *L*(*t*).

#### 3.2.3 Continuous control with fixed endpoint

For the fixed endpoint problem, we proceed with the same state equations as for the AML continuous control problem given in Equation (15), however we now impose a terminal condition on the leukaemic population; *L*(10) = 0.05. We seek to minimise the same quadratic cost function *J*, as considered in Equation (16). We form the same Hamiltonian given in Equation (17) and derive the same co-state, Equation (18), and expression for the control, Equation (20).

We obtain final time conditions, *λ*_1_(10) = *λ*_2_(10) = 0, via the transversality conditions as usual, however we do not prescribe *λ*_3_(10). Instead, we make two guesses for *λ*_3_(10); for instance, 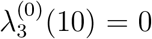 and 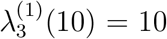. We then proceed by applying the adapted FBSM outlined in §2, using these guesses to initialise the secant method. Numerical results for the AML fixed endpoint control problem are presented in Figure 8. These results are produced using the adapted FBSM with *ω* = 0.55 in Σ = 434 iterations. This choice of *ω* minimises Σ for the AML fixed endpoint control problem solved with the FBSM without acceleration techniques. We discuss this further in §5.

**Fig. 8.**
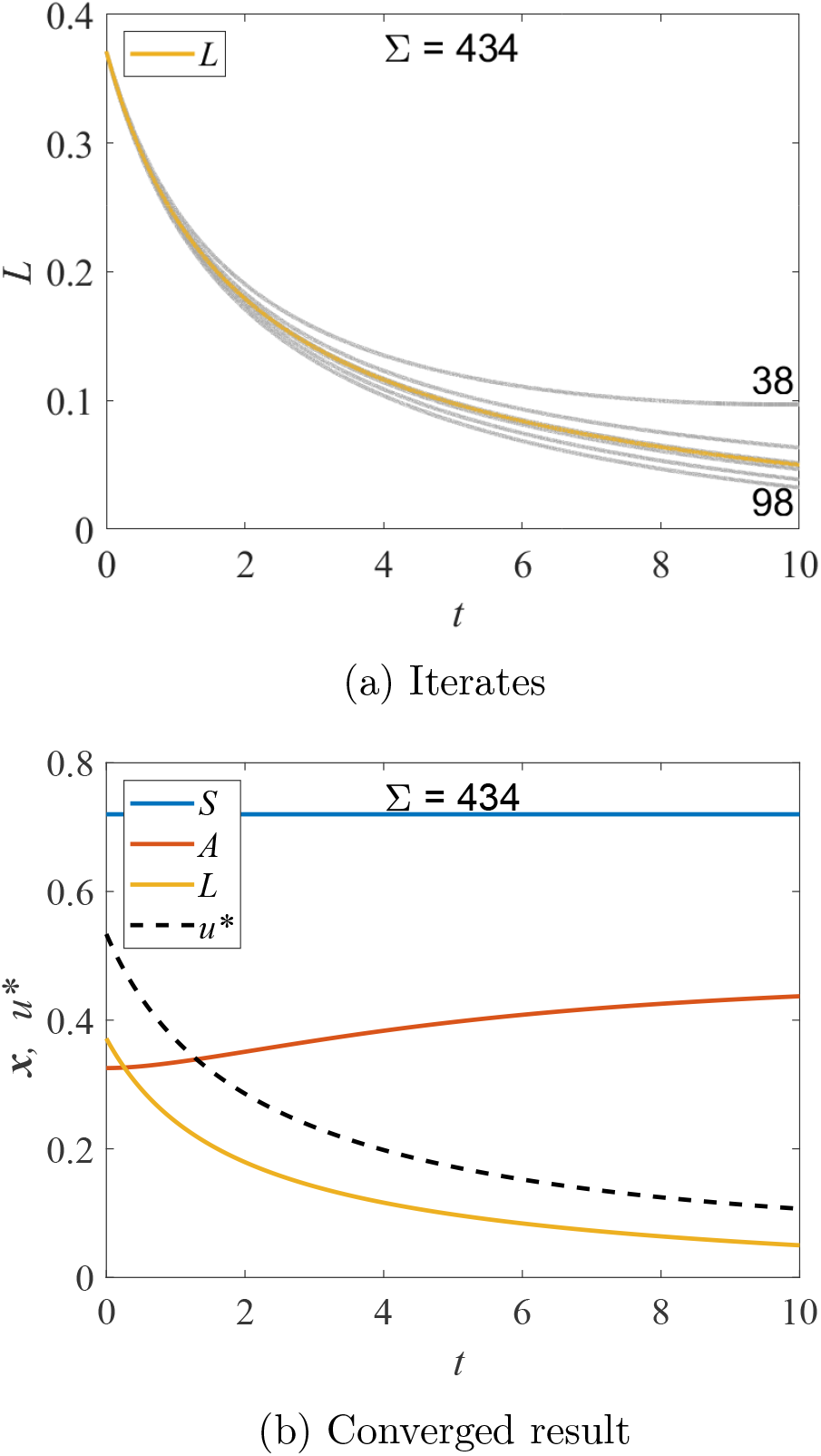
Results are presented for the AML problem with specified terminal state, *L*(*t_N_*) = 0.05, solved using the adapted FBSM. Each underlying FBSM problem is solved with model parameters given in Table 1, time-step d*t* = 4.88 × 10^−4^, over the interval 0 *t* 10, with pay-off weightings of *a*_1_ = 1 for the control, and *a*_2_ = 2 for the state variable *L*(*t*). In (a) the *L*(*t*) iterates of the adapted FBSM are presented in grey; the converged solution satisfying *L*(*t_N_*) = 0.05 is plotted in yellow. We annotate 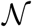 for the first 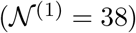 and second 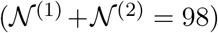 iterations of the adapted FBSM, based on initial guesses for *λ*_3_(*t_N_*) of *λ*_3_(*t_N_*) = 0 and *λ*_3_(*t_N_*) = 10. Due to the close proximity, subsequent iterations are not annotated. The cumulative function evaluations required for convergence of the adapted FBSM (Σ = 434) is indicated. The converged result for *L*(*t*), satisfying |*L*(*t_N_*) − 0.05| ≤ 1 × 10^−10^, is presented in (b); this figure also includes the optimal control, *u*\*(*t*) and trajectories for *S*(*t*) and *A*(*t*).

## 4 Iterative accelerators

In this section we outline several techniques for acceleration of iterative schemes. Where appropriate, we first present the univariate/scalar version of the method for familiarity, then provide the multivariate/vector analogue of the method for use with accelerating the FBSM. We attempt to use notation that aligns most closely with commonly used notation in the literature, while maintaining internal consistency in this work. In the scalar case, we consider the iterative process *x*^(*k*+1)^ = *f* (*x*^(*k*)^), where *x*^(*k*)^ is the *k*th iterate and *f* is the iterating function. In the vector case we consider *X*^(*k*+1)^ = *F* (*X*^(*k*)^), where 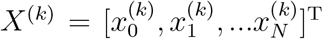 is the *k*th iterate, consisting of *N* + 1 values, and *F* = [*f*_0_*, f*_1_*,…f_N_*]^T^ is the *N* + 1 dimensional operator of the iterative process. For clarity, we stress that in the context of the acceleration algorithms applied to the FBSM, *X*^(*k*)^ is the discretised control in the *k*th iteration.

The acceleration methods considered in this work apply either to problems stated as fixed point iterations (as above), or as root-finding problems. For acceleration via root-finding algorithms, we can consider the complementary problems in the scalar and vector setting, respectively; *g*(*x*):= *x* − *f* (*x*) = 0 and *G*(*X*):= *X* − *F* (*X*) = **0**, where **0** is the zero column vector of length *N* + 1.

We note that many of the methods presented here can be written in several different forms. While some forms better facilitate analysis of aspects such as convergence speed and numerical stability, others emphasise ease of understanding and implementation. In this work we prioritise usability and present methods and algorithms in forms reflective of their implementation where possible. For the purpose of this work, we feel it is sufficient to present the methods and discuss their implementation without delving into their derivation or rigorous theoretical convergence results. For readers interested in these aspects, we suggest these articles [14,66], and numerical analysis texts [18,40].

### 4.1 Newton and Quasi-Newton methods

Newton’s method is one of the most prevalent root-finding algorithms, due to its relatively straightforward implementation and potential for quadratic convergence [40]. For a univariate function Newton’s method is given by

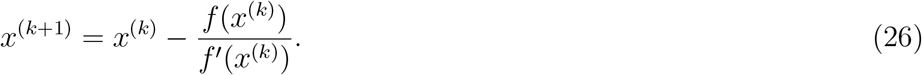

We arrive at the scalar secant method by replacing the derivative term, *f*′(*x*^(*k*)^), in Equation (26) with a finite difference approximation:

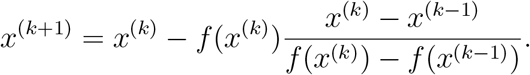

Newton’s method for multivariate systems is

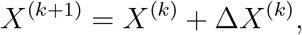

where Δ*X*^(*k*)^ is obtained by solving

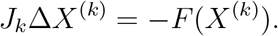

Here, *J_k_* is the Jacobian matrix of *F* evaluated at *X*^(*k*)^ [40]. Setting aside the interpretation of the Jacobian in the context of the FBSM; numerically approximating an *N* × *N* Jacobian matrix using finite differences requires 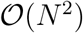 FBSM iterations at each Newton step. A range of Quasi-Newton methods have been developed to minimise the computational expense associated with computing the Jacobian at each Newton step. It is not immediately apparent how the secant method should be extended to multivariate systems, but one such interpretation is the Quasi-Newton Broyden’s method. Broyden’s method reduces the number of function evaluations required at each Newton step by forming the full Jacobian only initially, then updating the Jacobian matrix via a rank-one update based on the secant method [15,18]. We later discuss the Wegstein method [82], which is another interpretation of the secant method in multivariate settings.

In the context of accelerating the FBSM, techniques that require forming or approximating a full Jacobian, even once, are not appropriate. We have an *N* + 1 dimensional system, where *N* + 1 is the number of time points in the discretisation of the ODEs, so we expect *N* to be large, relative to the number of iterations required for the FBSM to converge without acceleration techniques, via Equation (6). As such, we restrict our focus to Jacobian-free methods in the remainder of this section; in particular, we discuss and implement the Wegstein and Aitken-Steffensen methods and Anderson acceleration. We provide a broad overview alongside the key equations here, and provide complete algorithms alongside notes for implementation in §4 of the supplementary material.

### 4.2 Wegstein method

Wegstein’s method can be thought of as an element-wise extension of the secant method to multivariate systems [35]. Although Wegstein’s method appears less popular than other methods considered in this work, it has found practical utility, particularly in chemical and process engineering software [51,77]. We include it here due to the striking similarity it bares to the control update with relaxation presented in Equation (6). It is also one of the more straightforward techniques, both in conception and implementation:

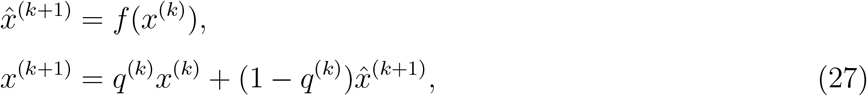

where 

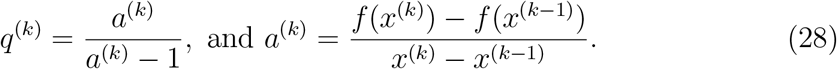

In implementation, from an initial value *x*_0_ it is necessary to perform two function evaluations; i.e. *x*_1_ = *f* (*x*_0_), and *f* (*x*_1_), before it is possible to compute Equation (28) for the Wegstein method [82]. In subsequent iterations only one new function evaluation is required.

The extension of Wegstein’s method to multivariate systems follows exactly the process outlined in Equations (27) and (28), as it is extended element-wise. While convergence is guaranteed when using Wegstein’s method for a single nonlinear equation, the uncoupling implied by the element-wise extension can lead to divergence [60].

In Equations (27) and (28) *q*^(*k*)^ denotes *q* in the *k*th iteration, however, we note that it is not necessarily most effective to update *q* every iteration. As such, in this work we explore various updating regimes. There is also the option of applying bounds on *q*. Bounds of −5 *< q_i_ <* 0, ∀*i*, where *i* denotes the *i*th element of the system, are frequently applied when implementing Wegstein’s method [7,70]. This bounding appears to work reasonably well for the small nonlinear test systems we consider in §5 of the supplementary material, although we were not able to identify a theoretical result supporting this specific choice. For the control problems we consider, this bounding is not effective, so we apply different bounds, discussed further in §5. The univariate Wegstein method can be thought of as a modification of the Aitken method, which at the time the Wegstein method was developed, was only understood for the univariate case [25].

### 4.3 Aitken-Steffensen method

Aitken’s Δ^2^ method, also referred to as Aitken’s delta-squared process or Aitken extrapolation, was originally posed by Aitken in 1927 as a means of extending Bernoulli’s method of approximating the largest root of an algebraic equation. This extension facilitates numerically approximating not only the largest root, but all roots of the equation [3]. Aitken’s method generates a 481 new sequence, 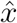, in parallel to the fixed point iteration.

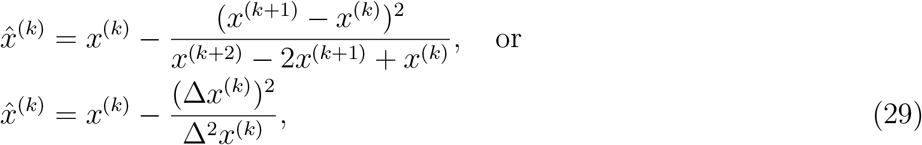

where Δ is the difference operator; Δ*x*^(*k*)^ = *x*^(*k*+1)^ − *x*^(*k*)^, and the higher order operator is applied recursively; Δ^2^*x*^(*k*)^ = Δ(Δ*x*^(*k*)^) = Δ*x*^(*k*+1)^ − Δ*x*^(*k*)^ [40]. From an initial value, *x*^(0)^, two function evaluations; iterations of the underlying fixed point process, must be performed to obtain *x*^(1)^ and *x*^(2)^, before Equation (29) can be computed.

The derivation of Aitken’s method assumes an underlying linearly converging series of iterates. The order of convergence of the resulting Aitken accelerated series is still linear, however this series converges faster than the original series [18]. We discuss Aitken’s Δ^2^ method and Steffensen iteration together, as Steffensen iteration is a straightforward extension of Aitken’s method, whereby the Aitken value, 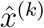, is used to continue the fixed point iteration, i.e. 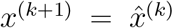. Despite the striking similarity, Steffensen’s method was seemingly developed shortly after (1933) and without knowledge of Aitken’s method [79]. Steffensen iteration can achieve quadratic convergence [2,40,55]. Further theoretical convergence results for the Steffensen method are established by Nievergelt [56] and in a series of papers by Noda [57,58,59].

Aitken and Steffensen iteration can be extended to multivariate systems [40]. In the following statements we outline the method for an *N* + 1 dimensional system; 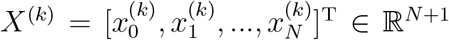, as appropriate for use with the FBSM:

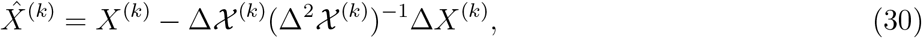

where Δ*X*^(*k*)^ = *X*^(*k*+1)^ − *X*^(*k*)^, 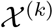 is a matrix constructed with columns (*X*^(*k*)^*, X*^(*k*+1)^*,…, X*^(*k*+*N*)^), such that 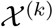 is a square matrix of dimension *N* +1, with 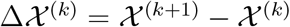, and 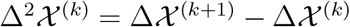.

In the form given by Equation (30) there are glaring issues with using the Steffensen method to accelerate convergence of the FBSM. Setting aside the question of whether 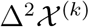 is invertible, forming 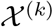 would require 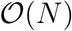 iterations of the FBSM to be performed, and since *N* relates to the number of time points in the discretisation of the ODEs in the FBSM, we expect *N* to be large, relative to the number of iterations required for the FBSM to converge without acceleration.

We instead consider a modification of the Steffensen method, requiring fewer function evaluations per iteration. Introduce *m < N*, and define Δ*X*^(*k*)^ = *X*^(*k*+1)^ − *X*^(*k*)^ as before, 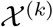 is now a rectangular matrix constructed with columns (*X*^(*k*)^*, X*^(*k*+1)^*,…, X*^(*k*+*m*+1)^), such that 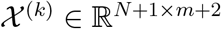, with 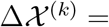 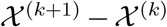, and 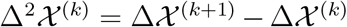, both of dimension *N* + 1 × *m*. We now interpret the matrix inverse in Equation (30) as the Moore-Penrose pseudoinverse [62], a generalisation of the matrix inverse for singular and rectangular matrices; we discuss this further in §3 of the supplementary material. This partial implementation requires only *m* + 1 function evaluations per iteration. For the remainder of this document when referring to the Steffensen method we are specifically referring to this partial Steffensen implementation. We present the derivation of the multivariate Aitken-Steffensen method and outline where the partial implementation differs, in § 3 of the supplementary material.

### 4.4 Anderson Acceleration

Anderson Acceleration or Anderson Mixing, originally denoted as the extrapolation algorithm by Anderson in the 1960s [5], is a technique developed for accelerating convergence of fixed point iteration problems with slowly converging Picard iterations [6]. Anderson Acceleration is of particular interest in this work, as it has recently been implemented to accelerate the convergence of a regularised version of the FBSM [48]. In contrast to a standard fixed point iteration; whereby the next iterate depends only on the immediately preceding iterate, Anderson Acceleration has ‘memory’ through the inclusion of previous iterates [30]. Unlike other methods considered in this work, Anderson Acceleration explicitly utilises the differences between residuals of subsequent iterates alongside iterates and their differences in computing future iterates.

Anderson Acceleration involves solving a least-squares problem at each iteration. The problem can be expressed in both constrained and unconstrained forms, with the updating step dependent on the form [32,81]. We solve the following unconstrained least-squares problem in each iteration of Anderson Acceleration:

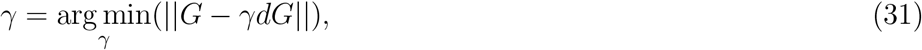

where arg min(·) returns the argument, *γ*, that minimises the expression in Equation (31). The corresponding updating step is

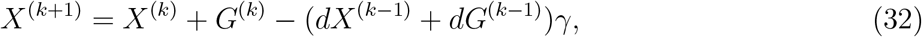

where *G*^(*k*)^ = *F* (*X*^(*k*)^) − *X*^(*k*)^ is the residual, *dX*^(*k*)^ is a matrix with columns (Δ*X*^(*k*−*m*)^, Δ*X*^(*k*−*m*+1)^*,…,* Δ*X*^(*k*)^), and *dG*^(*k*)^ is a matrix with columns (Δ*G*^(*k*−*m*)^, Δ*G*^(*k*−*m*+1)^*,…,* Δ*G*^(*k*)^), and *m* indicates the number of previous iterates that are incorporated.

### 4.5 Acceleration methods applied to typical fixed point problems

As a precursor to implementing these acceleration methods for control problems, we apply them to solve example nonlinear systems of dimension 2 × 2, 3 × 3, and 4 × 4. We provide these systems and the results of the acceleration methods compared to standard fixed point iteration in §5 of the supplementary material. We do not discuss these results in detail, although broad comparisons regarding the application of the acceleration methods to these systems and to control problems are made in §6. We provide code on GitHub for implementing the acceleration algorithms to solve systems of arbitrary size.

## 5 Acceleration results

In this section we discuss the results of applying the acceleration algorithms. When discussing results we are solely focused on reducing 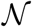, the number of function evaluations required for the control problems to reach convergence; as in all convergent cases we arrive at the same optimal control results. We first discuss the aspects of each method that can be tuned, then outline the results of the standard FBSM with the best choice of *ω* but without acceleration methods applied, to establish a baseline against which to compare the acceleration methods. A detailed suite of results for each control problem and each acceleration method, for various combinations of tuning parameters, is provided in §6 of the supplementary material.

### 5.1 Tuning

Each method we consider has parameters that can be tuned to improve performance for a given problem. For the FBSM without acceleration, we can select *ω* ∈ [0, 1); the parameter that weights the contribution of the control from the previous iteration, and the newly calculated control, to the control used in the next iteration, as stated in Equation (6). Control problems based on the linear model are able to converge via direct updating, as given in Equation (5), equivalent to *ω* = 0. Increasing *ω* in this case only serves to increase 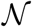, so we do not attempt to tune *ω* when considering the linear model. Using the standard FBSM without acceleration the continuous linear problem requires 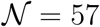 while the bang-bang linear problem requires only 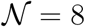.

In Figure 9 we plot 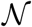 against *ω* ∈ [0, 1), for the continuous and bang-bang AML problems. As expected, for small *ω* we find that the problem does not converge, and for large *ω*, 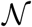 increases. For the continuous AML problem we identify *ω* = 0.55 as the best choice, with 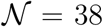. For the bang-bang AML problem we find that *ω* = 0.4 is best, with 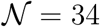.

**Fig. 9.**
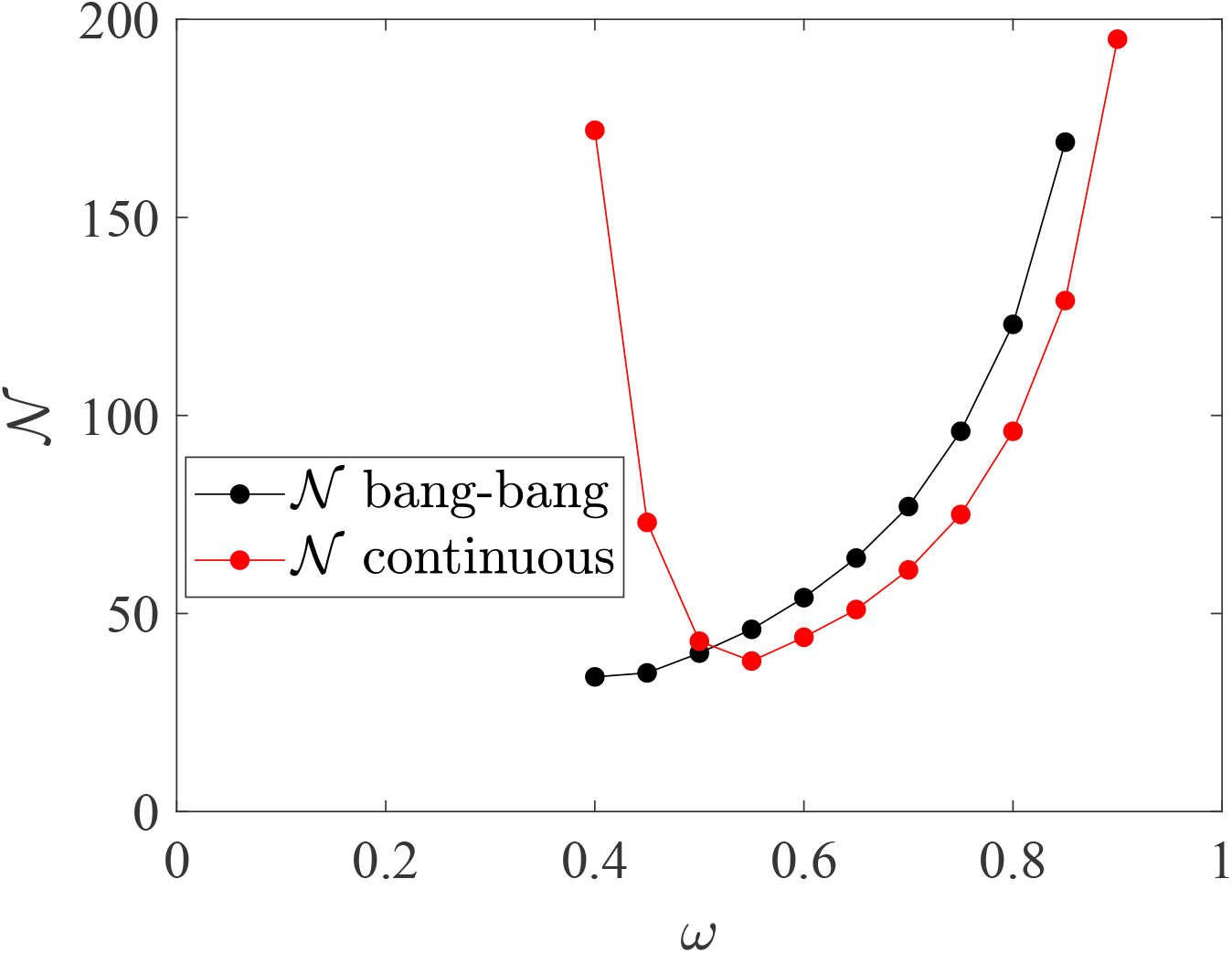
Here we plot 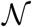 against *ω* ∈ [0, 1) in increments of 0.05, for the AML continuous (in red) and bang-bang (in black) control problems, using FBSM with no acceleration. Results correspond to model parameters given in Table 1, time-step d*t* = 4.88 × 10^−4^, over the interval 0 ≤ *t* ≤ 10. The continuous problem is solved with pay-off weightings of *a*_1_ = 1 for control, and *a*_2_ = 2 for the state variable *L*(*t*), while the bang-bang problem is solved with *a*_1_ = 1 and *a*_2_ = 3. Where an *ω* value does not have a corresponding marker, this indicates that the procedure fails to converge within 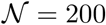.

Recall that fixed endpoint problems are solved using the adapted FBSM; this entails solving several control problems to convergence with the FBSM. Each of these control problems can have a different optimal *ω*. In this instance, *ω* = 0.55 also happens to be best for the AML fixed endpoint problem if holding *ω* constant, when considering *ω* ∈ [0, 1) at increments of 0.05. These *ω* values will not coincide in general. When applying the acceleration methods to the fixed endpoint problems, we employ the tuning parameters that perform best for the continuous problem. This does not imply that we are using the best tuning parameters for the acceleration methods in the context of the fixed endpoint problem. Importantly, this demonstrates whether or not the techniques can effectively reduce Σ, the cumulative function evaluations required for convergence of the adapted FBSM for fixed endpoint problems, without requiring prohibitive tuning.

With the Wegstein method, we only select *ω* for the two FBSM iterations required for initialisation, and specify *n*, such that we update *q* every *n*th iteration. We generate results for *n* ∈ {1, 2*,…,* 10}. We also bound *q*, however identifying suitable bounds is challenging. In this work we select bounds that perform reasonably, but acknowledge that we do not search for optimal bounds, nor do we think that attempting to do so is realistic. This drawback of Wegstein’s method contributes to its inconsistent performance relative to other methods. For the partial Aitken-Steffensen methods, we choose *ω*, and the parameter *m* that specifies the dimension of the *N* + 1 × *m* matrices in the updating step, requiring *m* + 1 function evaluations per iteration. We generate results for *m* ∈ {1, 2*,…,* 10}. Similarly, for Anderson Acceleration we select *ω* and *M*; where *M* determines the maximum number of previous iterations to retain when solving the least squares problem and performing the updating step. We produce results for *M* ∈ {1, 2*,…,* 10}.

### 5.2 Wegstein method

For the continuous linear problem, we apply bounds of −2 ≤ *q* ≤ 0. For the bang-bang linear problem, we leave *q* unbounded. For both AML problems we apply bounds −1 ≤ *q* ≤ 1. We explore the effect of updating *q* every *n*th iteration, *n* ∈ {1, 2*,…,* 10}. For the continuous linear problem, *n* = 4 minimises 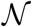, although *n* ∈ {1, 2*,…,* 5} all perform well. For the linear bang-bang problem, the Wegstein method converges without bounding on *q*, and varying *n* does not affect convergence. The Wegstein method outperforms other acceleration methods for the linear bang-bang problem with 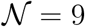, but does not improve upon 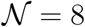 for the FBSM without acceleration.

For the continuous AML problem, the performance of Wegstein’s method is inconsistent. With *n* = 6 and *ω* = 0.55, the Wegstein method achieves convergence with 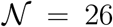, outperforming the FBSM; however, almost every other combination of tuning parameters considered with *ω* ∈ [0, 1) and *n* ∈ {1, 2*,…,* 10} require larger 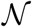 than the FBSM without acceleration. Generally, increasing *n* produces worse outcomes. We do, however, observe that the Wegstein method can induce convergence for *ω <* 0.4, where the standard FBSM does not converge. For the bang-bang AML problem, the Wegstein method appears more robust; consistently outperforming the standard FBSM across most of the tuning parameter space. The best result requires only 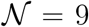, with *ω* = 0 and *n* = 7, although several other combinations of tuning parameters are similarly successful. For *ω* ≥ 0.4, corresponding to values that the underlying FBSM converges, we find that moderate *n* ∈ {3, 4*,…,* 7} produces the best results; while for smaller *ω*, larger *n* ∈ {6, 7*,…,* 10} consistently performs best. Once again we observe that convergence is achieved for *ω* values where the underlying FBSM would not converge.

For the linear fixed endpoint problem the adapted FBSM with the Wegstein method consistently generates a moderate reduction in Σ, compared to the adapted FBSM without acceleration, for all *n* ∈ {1, 2*,…,* 10}, −2 ≤ *q* ≤ 0. For the AML fixed endpoint problem we do not observe improvement. Using the tuning parameters that perform best for the continuous AML problem, we find that Σ for the adapted FBSM with Wegstein’s method is more than double that of the adapted FBSM without acceleration. This results from the inconsistency of Wegstein’s method with poor tuning. In §6 of the supplementary material it can be seen that some control problems within the adapted FBSM that require 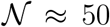 without Wegstein’s method, require 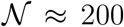 with the specified Wegstein tuning parameters.

### 5.3 Partial Aitken-Steffensen method

Both Aitken and Steffensen methods significantly and consistently outperform the FBSM without acceleration for the continuous linear problem. The Aitken method performs best for *m* ∈ {1, 2, 3}, requiring 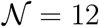. Steffensen’s method performs best when *m* = 6, requiring only 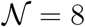. In the linear bang-bang case, both Aitken and Steffensen methods perform marginally worse than the FBSM without acceleration, which requires only 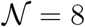. In the best cases, with *m* = 1 the Aitken method requires 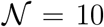, and with *m* = 7 the Steffensen method requires 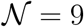.

For the continuous AML problem, we observe a stark difference between the Aitken and Steffensen methods; while the Steffensen method is able to achieve convergence for values of *ω* where the underlying FBSM fails to converge; *ω* ≤ 0.35, particularly for *m* ∈ {1, 2, 3, 4}, the Aitken method only converges to the optimal control for *ω* values where the underlying FBSM converges. For *ω* ≤ 0.35 the Aitken method achieves apparent convergence; the iterative procedure terminates as the convergence criteria is met. However, explicitly calculating the pay-off associated with these controls via Equation (16), and comparing this result to the pay-off associated with the control obtained via the standard FBSM, indicates that the controls obtained via the Aitken method for *ω* ≤ 0.35 are not optimal, as they fail to minimise *J*. The best result for the Aitken method, with *ω* = 0.5 and *m* = 5, requires 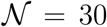, marginally improving on the FBSM without acceleration, requiring 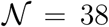. Steffensen’s method produces more significant improvements, requiring only 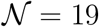 with *ω* = 0.5 and *m* = 5. In each case neighbouring combinations of tuning parameters also yield equivalent or comparable improvement over the standard FBSM. In the bang-bang AML problem, we observe similar behaviour; for *ω* values that the underlying FBSM fails to converge, the Steffensen method consistently converges. The Aitken method achieves apparent convergence for these values of *ω*; the iterative procedure terminates as the convergence criteria is met, however the resulting controls contain intermediate values between the lower and upper bounds. As such the resulting controls are not bang-bang, so we treat these results as failing to converge. At best, Aitken’s method requires 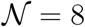, with *ω* = 0.5 and *m* = 1, while Steffensen’s requires only 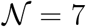, with *ω* = 0.5 and *m* = 5. The vast majority of tuning parameter combinations yield improvements over the 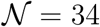 of the standard FBSM.

Aitken and Steffensen methods consistently offer significant improvement over the standard adapted FBSM for the linear fixed endpoint problem for *m* ∈ {1, 2*,…,* 10}, with the exception of *m* = 1 for the Steffensen method, which yields only marginal improvement. Using the best performing tuning parameters for the continuous AML problem, we find that both Aitken and Steffensen methods improve upon the standard adapted FBSM for the AML fixed endpoint problem. Relative to Σ = 434 required without acceleration, the Σ = 360 required with Aitken’s method reflects a modest improvement, while the Σ = 238 required with the Steffensen method is a significant improvement.

### 5.4 Anderson Acceleration

Anderson Acceleration performs exceptionally well on the continuous linear problem, requiring only 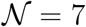 for *M* ∈ {4, 5*,…,* 10} compared to 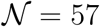 for the standard FBSM. For the linear bang-bang problem however, it is the worst performing acceleration method; achieving at best 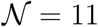, with *M* = 1.

Similarly to the Wegstein and Steffensen methods, Anderson Acceleration achieves convergence in both the continuous and bang-bang AML problems for *ω* values where the underlying FBSM fails to converge. Anderson Acceleration achieves the best individual result for the continuous AML problem, requiring only 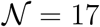, with *ω* = 0.85 and *M* = 6. Again, we observe comparable improvement over a wide range of tuning parameters. For the bang-bang AML problem Anderson Acceleration consistently outperforms FBSM without acceleration, particularly for *ω <* 0.7, at best requiring 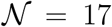, with *ω* = 0.35 and *M* ∈ {7, 8, 9, 10}, with other non-neighbouring tuning parameter combinations also yielding 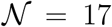.

For both the linear and AML fixed endpoint problems Anderson Acceleration produces the most significant reduction in Σ, and improves upon the adapted FBSM over a wide range of tuning parameters. In the linear case, Anderson Acceleration requires only Σ = 24 for *M* ∈ {4, 5*,…,* 10}. In the AML fixed endpoint problem, using the tuning parameters that perform best for the continuous AML problem, Anderson Acceleration converges in only Σ = 204; less than half as many as the standard adapted FBSM.

### 5.5 Method comparison with best tuning

Results presented in Figure 10 and Figure 11 provide comparison of the error, *ε*, as each method approaches convergence, for the linear and AML problems, respectively. Error is measured as the Euclidean norm of the difference between subsequent controls; *ε* = ||*F* (*X*^(*k*)^) − *X*^(*k*)^||, with the exception of Aitken’s method, where error is measured as the difference between subsequent values in the Aitken series; 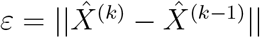. Convergence is achieved when *ε* ≤ 1 × 10^−10^, marked in black dash. In each case, we are plotting the result that minimises 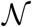 for each method, over the space of tuning parameters considered, including the best tuning of *ω* for the FBSM without acceleration. Error is plotted on a logarithmic scale. For the linear bang-bang problem with the Wegstein and Anderson methods, and the AML bang-bang problem with the Wegstein method, the error after the final iteration is *ε* = 0, as two subsequent iterates for the control are identical. This is represented on the logarithmic scale as a line that intersects the horizontal axis.

**Fig. 10.**
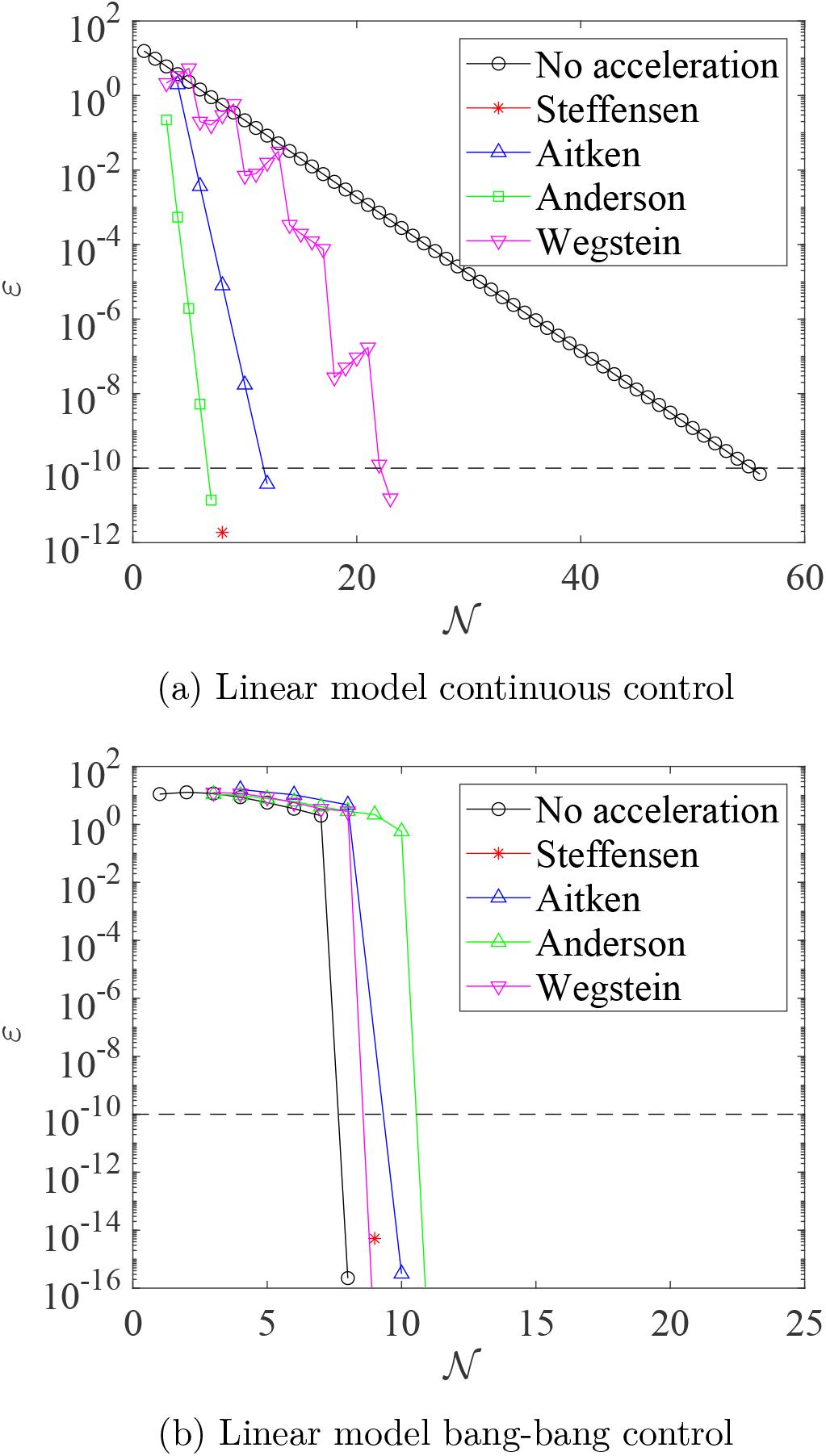
Convergence rates for the result that minimises 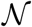, for each acceleration method, when applied to the linear control problems. Results in this plot are produced with model parameter *γ* = 0.5, time-step d*t* = 3.91 × 10^−3^, over the interval *t* 1, with pay-off weighting *a* = *b* = 1 for the continuous control (a), and *a* = 1, *b* = 3 for the bang-bang control (b). The tolerance of 1 × 10^−10^ required for convergence is marked in black dash. As the methods do not necessarily use the same number of function evaluations per iteration, markers indicate each time *ε* is computed. Continuous control results correspond to the FBSM with no acceleration, the partial Steffensen method with *m* = 6, partial Aitken method with *m* = 1, Anderson Acceleration with *M* = 4 and Wegstein with bounds 2 ≤ *q* ≤ 0, updating *q* every 4th iteration. Bang-bang results correspond to the FBSM with no acceleration, the partial Steffensen method with *m* = 7, partial Aitken method with *m* = 1, Anderson Acceleration with *M* = 1 and Wegstein without bounds on *q*, updating *q* every iteration. The standard FBSM outperformed all acceleration methods in solving the linear bang-bang control problem. We attribute this to how few iterations were required 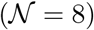 for convergence without acceleration.

**Fig. 11.**
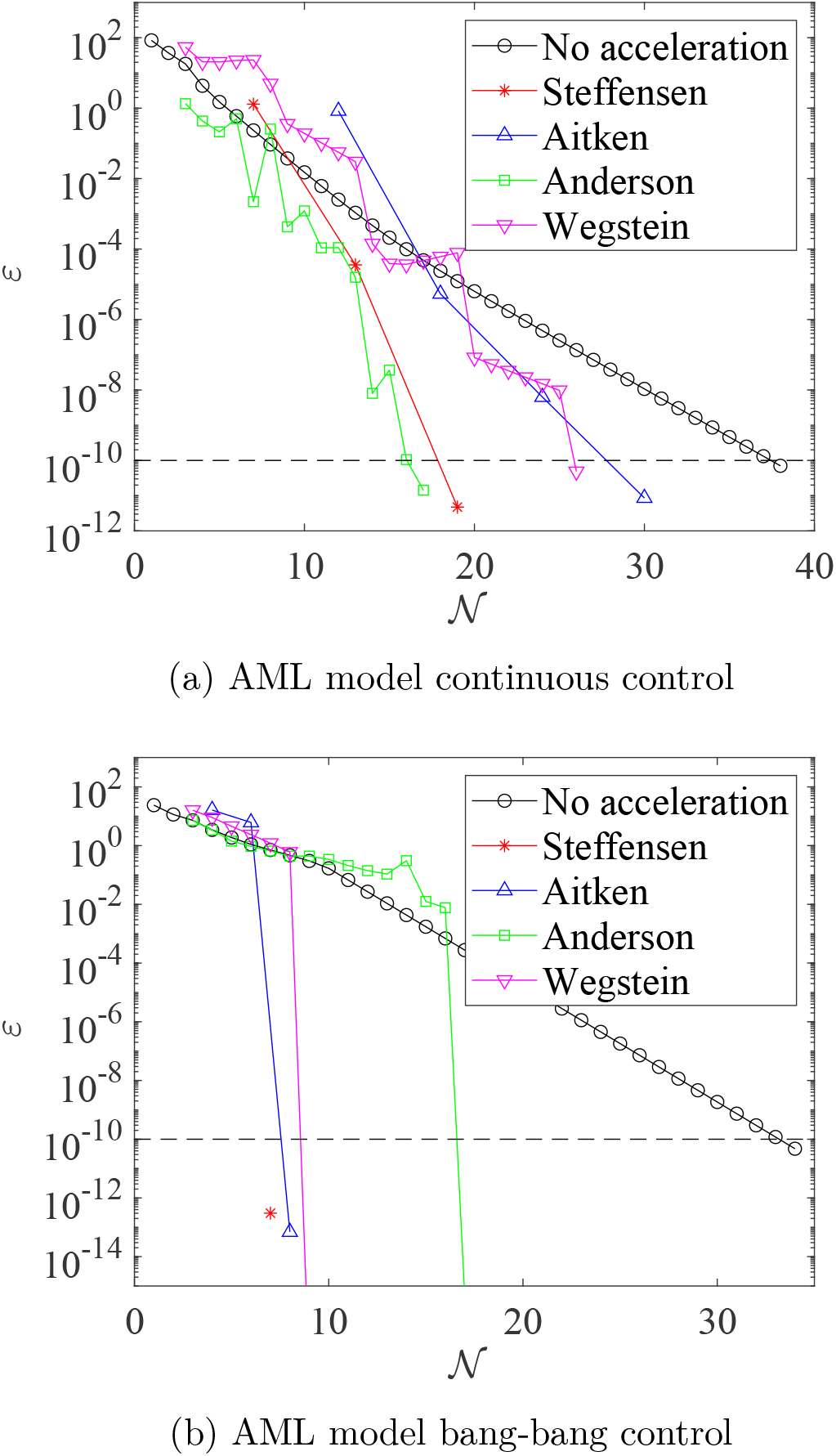
Convergence rates for the converged result that minimises 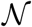, for each acceleration method, when applied to the AML control problems. Results in this plot are produced with model parameter *γ* = 0.5, time-step d*t* = 4.88 × 10^−4^, over the interval 0 ≤ *t* ≤ 1, with pay-off weighting *a* = *b* = 1 for the continuous control (a), and *a* = 1, *b* = 3 for the bang-bang control (b). The tolerance of 10^−10^ required for convergence is marked in black dash. As the methods do not necessarily use the same number of function evaluations per iteration, markers indicate each time *ε* is computed. Continuous control results correspond to the FBSM with no acceleration, *ω* = 0.55, the partial Steffensen and partial Aitken methods with *m* = 5 and *ω* = 0.5, Anderson Acceleration with *M* = 6 and *ω* = 0.85, and Wegstein method with *ω* = 0.55, bounds −1 *< q <* 1, updating *q* every 6th iteration. Bang-bang results correspond to the FBSM with no acceleration, *ω* = 0.4, the partial Steffensen method with *m* = 5 and *ω* = 0.5, partial Aitken method with *m* = 1 and *ω* = 0.5, Anderson Acceleration with *M* = 7 and *ω* = 0.35, and Wegstein method with *ω* = 0, bounds −1 *< q <* 1, updating *q* on the 7th iteration.

## 6 Discussion

Modelling processes in the life sciences is complex; frequently involving large state systems consisting of several ODEs [12,28,78]. The acceleration methods we implement act only on the control term; the number and form of state equations has no bearing on the mathematical and computational complexity of the acceleration methods. As such, the methods scale excellently with system complexity. In this section we discuss the results presented in §5, and draw insights into the convergence behaviour of the FBSM when augmented with acceleration techniques. We highlight opportunities for application of these methods, and outline several avenues for further investigation.

### 6.1 Acceleration outcomes

In evaluating the performance of each acceleration method, we are interested in: (1) how significantly they are able to reduce 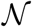, (2) method robustness, and (3) method accessibility. In this context we use robustness to refer to how consistently the method outperforms the best tuned FBSM over the range of tuning parameters considered. We judge the accessibility of each method based on implementation and conceptual complexity. Overall, we find that the acceleration methods, particularly Anderson and Steffensen, significantly and robustly reduce 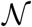. Anderson Acceleration appears most effective for continuous control, while the Steffensen method appears best for bang-bang control. The Aitken method occasionally outperforms Steffensen, but overwhelmingly the Steffensen method appears to be the better option of the two for the range of parameters we consider. Implementing the Anderson and Steffensen methods introduces challenge beyond that of the underlying FBSM, although it is not prohibitively difficult; particularly with reference to the code where we implement these methods, that we make available on GitHub. Both methods introduce conceptual complexity, perhaps marginally less-so for the Steffensen method due to the similarities it shares with the familiar Newton’s method.

We produce heatmaps to visualise the convergence behaviour of the acceleration methods across the range of tuning parameters considered. Figure 12 corresponds to the AML continuous control problem, while Figure 13 corresponds to the AML bang-bang control problem. Recall that with the tuning of *ω* that minimises 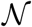, the FBSM with no acceleration requires 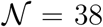 for the AML continuous control problem, and 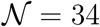 for the AML bang-bang control problem. Tuning parameter combinations that reflect a reduction in 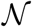 relative to the these FBSM results are shaded in the green spectrum, while worse performing combinations are shaded in the red spectrum. The midpoint of the colour spectra, yellow, corresponds to the FBSM result with the best tuning, without acceleration. Simulations are terminated when 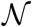 exceeds 100; reflecting a combination of tuning parameters that do not yield convergence within this specified maximum. Data supporting these heatmaps, and similar results for the linear control problems are provided in §6 of the supplementary material.

**Fig. 12.**
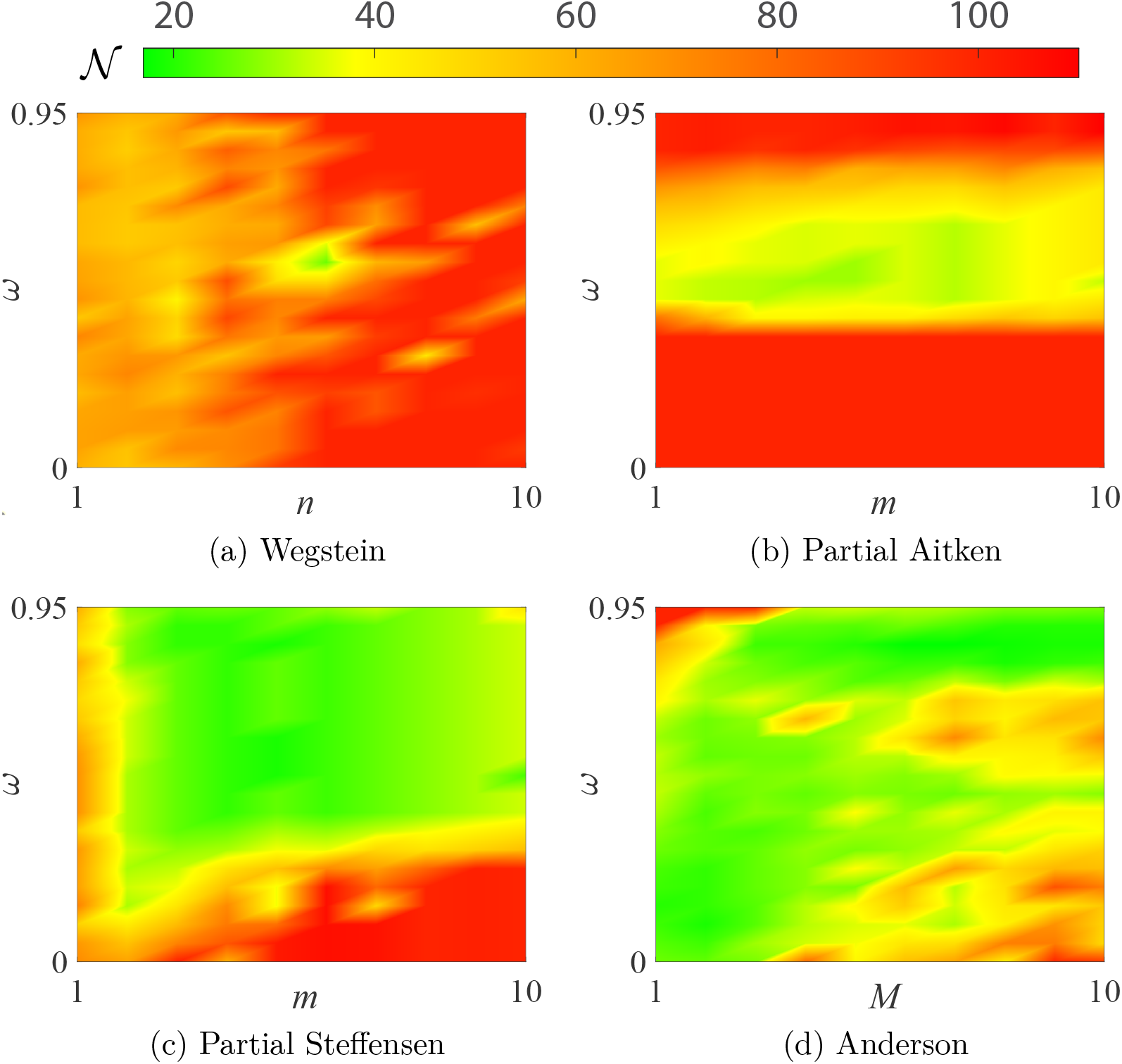
This heatmap provides insight into the convergence behaviour of the acceleration methods for the AML continuous control problem. Here we visualise 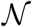 against *ω* and the method specific tuning parameter; *n* for Wegstein (a), *m* for partial Aitken (b) and partial Steffensen (c), and *M* for Anderson Acceleration (d). Tuning parameter combinations requiring 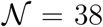, equivalent to the best tuned FBSM without acceleration, are shaded yellow. Colours in the green-yellow spectrum represent a reduction in 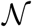 relative to FBSM without acceleration, while colours in the yellow-red spectrum represent an increase in 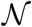.

**Fig. 13.**
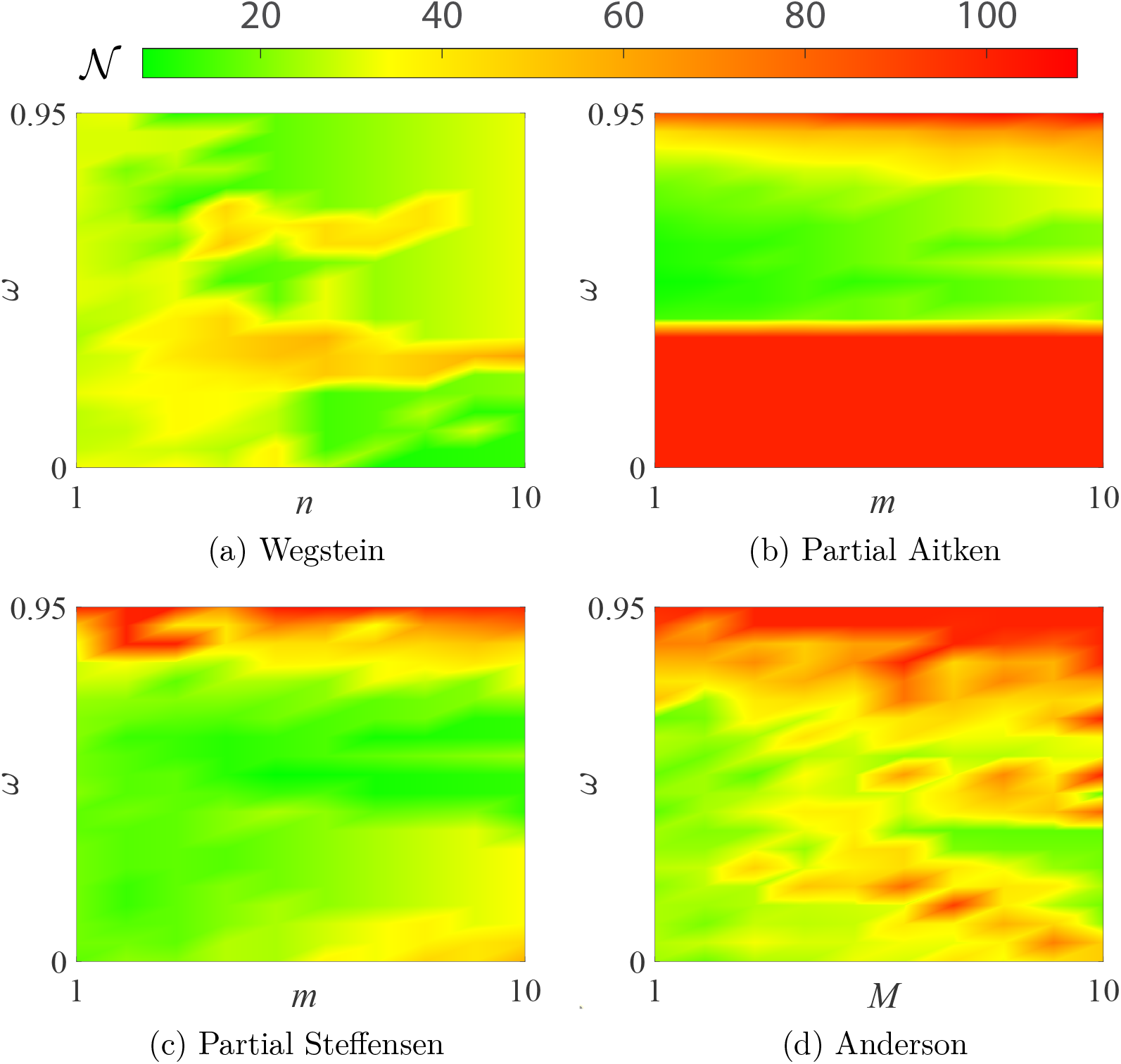
This heatmap provides insight into the convergence behaviour of the acceleration methods for the AML bang-bang control problem. Here we visualise 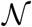 against *ω* and the method specific tuning parameter; *n* for Wegstein (a), *m* for partial Aitken (b) and partial Steffensen (c), *M* for Anderson Acceleration (d). Tuning parameter combinations requiring 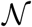, equivalent to the best tuned FBSM without acceleration, are shaded yellow. Colours in the green-yellow spectrum rep-resent a reduction in 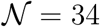 relative to FBSM without acceleration, while colours in the yellow-red spectrum represent an increase in 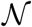.

In identifying tuning parameter combinations that yield significant reductions in 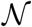, we are looking for bright green areas in the heatmaps. We assess the robustness of each method by considering whether we observe large contiguous areas in the green spectrum, such as in Figure 12(c), indicating robustness, or patchy areas with both green spectrum and red spectrum, such as Figure 13(d), suggesting a lack of robustness.

In Table 2 we provide our subjective but informed rating of the methods against the criteria of reduction in 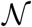, robustness and accessibility. We consider the continuous and bang-bang control cases separately in terms of reduction in 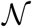 and robustness.

**Table 2:** Method comparison

| | Reduction in $\mathcal{N}$ | | Robustness | | Accessibility | |
| --- | --- | --- | --- | --- | --- | --- |
| Method | Cts | BB | Cts | BB | Imp | Complexity |
| FBSM | ∼ | ∼ | ✓ | ✓ | ✓ | ✓ |
| Wegstein | ∼ | ✓✓ | ✗ | ✓ | ✓ | ∼ |
| Aitken | ✓ | ✓ | ∼ | ✓ | ∼ | ∼ |
| Steffensen | ✓✓ | ✓✓ | ✓ | ✓✓ | ∼ | ∼ |
| Anderson | ✓✓ | ✓ | ✓ | ✓ | ∼ | ✗ |

We rate the methods considered in this work against key factors such as the reduction in $\mathcal{N}$ that they deliver and how robustly they perform over the range of tuning parameters considered, for both continuous (Cts) and bang-bang (BB) control problems. We also consider how accessible the methods are from the standpoints of ease of implementation (Imp) and conceptual complexity. Methods are rated as being either strongly positive (✓✓), positive (✓), neutral ( ∼ ), negative (✗) against each aspect.

Despite its conceptual simplicity and straightforward implementation, Wegstein’s method is significantly hampered by the difficulty in choosing bounds. If there were a more informed approach for identifying suitable bounds, Wegstein’s method could be particularly useful for bang-bang control problems. Due to the effect of *ω*, intermediate control iterates of the FBSM do not appear bang-bang; as such the bulk of 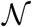 are incurred in refining the control about the switching points. Wegstein’s method can accelerate this refinement by adaptively setting *q_i_* = 0 where appropriate.

### 6.2 Convergence insights

As outlined in §5, the linear model control problems converge with *ω* = 0. It may at first seem counter-intuitive that Wegstein’s method can improve upon this, given that the computed *q* in Wegstein’s method acts as a stand-in for *ω*. There are two aspects of distinction that enable Wegstein’s method to generate improvement in this case; first, while *ω* is held constant both within the time discretisation and between iterations, the element-wise nature of Wegstein’s method enables each element of the discretisation to have a different value, *q_i_, i* ∈ {0, 1*,…, N*}, and *q* can be updated between iterations; secondly, observing the values of *q_i_* in Wegstein’s method indicates that *q <* 0 can be appropriate. This suggests that *ω <* 0 could also be used to accelerate the standard implementation of the FBSM. Preliminary investigation suggests that this is true for the linear model, however we do not pursue this further as we expect it to be of limited applicability beyond contrived problems.

We apply the acceleration methods to small nonlinear test systems in §5 of the supplementary material. We know these systems have multiple fixed points; all methods we consider aside from Aitken’s method, in some of our examples, reach different fixed points to fixed point iteration. In contrast, when applied to accelerate control problems, we observe only the Aitken method converging to a result other than the optimal control obtained via the FBSM, as discussed in §5. This apparent convergence of the Aitken method to controls that are not optimal is a significant deterrent to using the Aitken method in situations where the optimal control is not known a priori. Outside of this issue with the Aitken method, each acceleration method produces the same optimal control for a given problem. However, they each approach the converged control differently. In Figure 14 we plot the control as it converges for the FBSM and acceleration methods. In the code we provide on GitHub, users can view the control iterates of each method as they approach convergence. Visualising these methods as they converge gives insight into how they may be able to arrive at different fixed points; under certain circumstances the accelerated series of iterates may leave the basin of attraction for the fixed point found via fixed point iteration.

**Fig. 14.**
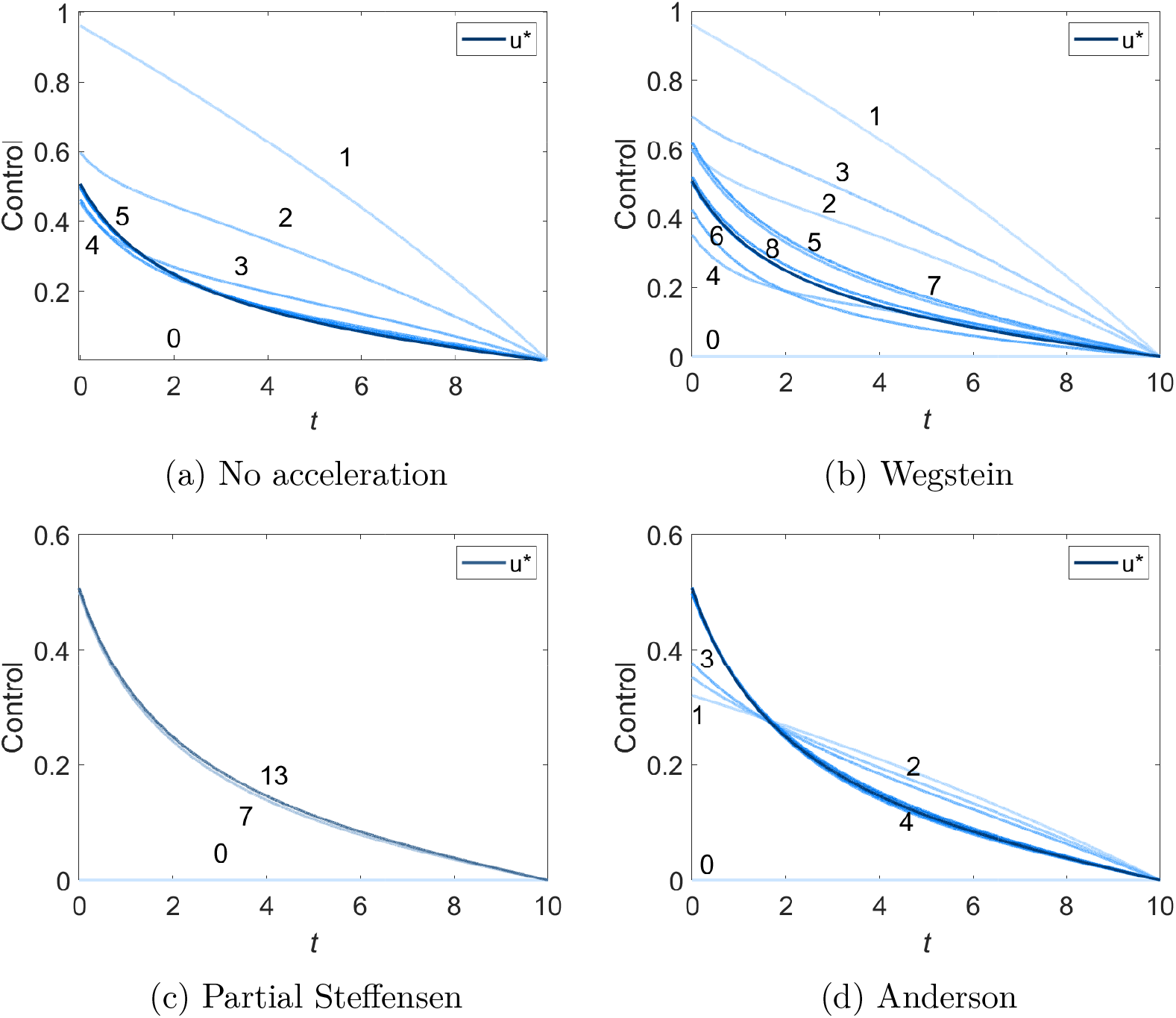
Here we observe the iterates of the control in the AML continuous control problem as it converges, for (a) the FBSM with no acceleration, (b) the Wegstein method, (c) the partial Steffensen method and (d) Anderson Acceleration. Initial iterates are shown in light blue, while darker blue denotes later iterates. Results for the Aitken method are not shown as they are visually similar to the Steffensen result. We present the results corresponding to the tuning parameters that minimise 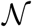, outlined in §5. Where it is visually distinguishable, we indicate the number of function evaluations corresponding to a particular iterate. While all methods produce the same eventual result for *u*\*, they follow considerably different series of iterates. Note that the vertical scale in (a) and (b) differs from that of (c) and (d).

### 6.3 Summary and outlook

In this work, we review the theory and implementation of the FBSM for solving TPBVPs that arise from application of PMP in solving optimal control problems. We study a single-variable linear model and a multiple-variable non-linear model and consider continuous, bang-bang and fixed endpoint control problems. Conceptualising the FBSM as a fixed point iteration, we leverage and adapt existing acceleration methods to significantly and robustly increase the convergence rate of the FBSM for a range of optimal control problems. The Anderson and partial Steffensen methods appear to perform best, without requiring prohibitive tuning.

Accelerating the convergence of the FBSM, and reducing the importance of appropriately selecting *ω* for a single control problem, is promising. That said, the real utility of the robust acceleration methods in this work is in application to families of control problems. We provide a glimpse of this benefit through considering fixed endpoint control problems, though other excellent opportunities for application arise due to the uncertainty prevalent in the life sciences. First, it is common for there to be uncertainty around model parameters and structure [29,34]. In this case solving optimal control problems over several model structures and sets of model parameters provides insight into the sensitivity of the control strategy [13,38,53,68]. Secondly, when performing multi-objective optimisation, a trade-off is made between objectives. For example, in Equation (21) we seek to minimise the cumulative negative impact of leukaemia and of the control; parameters *a*_1_ and *a*_2_ weight the relative important of each objective. In practical applications, it is not always clear how to determine these weightings. It can therefore be useful to generate a family of optimal controls that are each optimal for their specific combination of pay-off weighting parameters, akin to a Pareto frontier [4,43,49]. Producing these sets of control results benefits significantly from acceleration techniques such as the Anderson and Steffensen methods, where a consistent reduction in 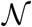 is obtained without optimal tuning.

Here, we have only considered systems subject to a single control. While this is reflective of the vast majority of applications featured in the control literature, there are instances where we are interested in applying multiple controls simultaneously [9,20,74]. The FBSM can be readily applied to solve problems with multiple controls [74]; a logical extension of this work is to adapt the acceleration methods or identify suitable alternative methods for accelerating convergence of the FBSM for problems with multiple controls.

Over a range of tuning parameters the Wegstein, Steffensen, and Anderson methods are able to induce convergence where the underlying FBSM fails to converge; such as in the AML control problems with *ω* = 0. This behaviour has been documented for Anderson acceleration [81] and Wegstein’s method [35] when applied to standard fixed point iteration problems. This presents an opportunity for future exploration, in identifying control problems that cannot be solved via the FBSM for any *ω*, and attempting to produce solutions using these acceleration techniques.

## Acknowledgements

JAS acknowledges support from the Australian Government Research Training Program and the AF Pillow Applied Mathematics Trust. JAS and KB acknowledge support from the Australian Centre of Excellence for Mathematical and Statistical Frontiers (CE140100049). MJS is supported by the Australian Research Council (DP200100177).

