## Supplementary material for "Implementation and acceleration of optimal control for systems biology"

**Contents**

| | | |
|---|---|---|
| 1 | Forward-backward sweep method algorithm | 3 |
| 2 | Single-variable linear continuous control analytical solution | 4 |
| 3 | Steffensen derivation | 7 |
| 4 | Acceleration algorithms | 13 |
| 4.1 | Wegstein method | 13 |
| 4.2 | Aitken-Steffensen method | 15 |
| 4.3 | Anderson Acceleration | 17 |
| 5 | Test nonlinear systems | 19 |
| 5.1 | Results | 20 |
| 6 | Control results | 27 |
| 6.1 | Linear continuous control problem | 29 |
| 6.2 | Linear bang-bang control problem | 29 |

*22 March 2021*

6.3 AML continuous control problem with the Wegstein method 30
6.4 AML continuous control problem with the partial Aitken method 31
6.5 AML continuous control problem with the partial Steffensen method 32
6.6 AML continuous control problem with Anderson acceleration 33
6.7 AML bang-bang control problem with the Wegstein method 34
6.8 AML bang-bang control problem with the partial Aitken method 35
6.9 AML bang-bang control problem with the partial Steffensen method 36
6.10 AML bang-bang control problem with Anderson acceleration 37
6.11 Linear fixed endpoint control problem 38
6.12 AML fixed endpoint control problem 38
References 40

#### Supporting code

Code for implementing the algorithms presented in this work is freely available on GitHub at https://github.com/Jesse-Sharp/Sharp2021.

* Corresponding author

### 1 Forward-backward sweep method algorithm

The algorithm for the forward-backward sweep method (FBSM) is adapted from [14,24]. Recall that in the standard control notation we represent the control as $u(t)$, the state variables as $\boldsymbol{x}(t)$, and the co-state variables as $\boldsymbol{\lambda}(t)$.

**Algorithm 1: Forward-backward sweep**

i. Make an initial guess, $u^{(0)}(t)$.
ii. Iterate for $k = 0, 1, ...,$ until converged or iteration limit met:
iii. Solve for $\boldsymbol{x}^{(k)}(t)$ forward in time using initial values $\boldsymbol{x}(0)$, and $u^{(k)}(t)$.
iv. Solve for $\boldsymbol{\lambda}^{(k)}(t)$ backwards in time from the transversality condition $\boldsymbol{\lambda}(t_N)$, using $u^{(k)}(t)$ and $\boldsymbol{x}^{(k)}(t)$.
v. Compute temporary update, $\hat{u}^{(k+1)}(t)$, using $\boldsymbol{x}^{(k)}(t)$, $\boldsymbol{\lambda}^{(k)}(t)$, and the optimality condition derived from minimising the Hamiltonian.
vi. Update $u^{(k+1)}(t) = \omega u^{(k)}(t) + (1 - \omega)\hat{u}^{(k+1)}(t)$.
vii. Check for convergence. If not converged, return to Step ii.

In Step i., an initial guess of $u^{(0)}(t) \equiv \mathbf{0}$ is often sufficient, though an initial guess closer to the optimal control can improve convergence. The choice of $\omega \in [0, 1)$ in Step vi. can significantly impact the rate of convergence, and whether or not the process converges at all [14,24]. We do not prescribe a specific convergence criterion in Step vii., as there are several valid choices. A general approach is to check if $\boldsymbol{x}^{(k)}(t)$, $\boldsymbol{\lambda}^{(k)}(t)$, and $u^{(k)}(t)$ are within a specified absolute or relative tolerance of their previous iteration values. For well-behaved systems, convergence of the control terms generally implies convergence of the associated state and co-state variables.

#### 2 Single-variable linear continuous control analytical solution

Consider the linear continuous control problem posed in the main document, repeated here as Equation (S1).

$$\frac{\mathrm{d}x(t)}{\mathrm{d}t} = \gamma x(t) + u(t), \quad x(0) = x_0, \quad \gamma > 0, \quad 0 \leq t \leq 1. \tag{S1}$$

The co-state equation is

$$\frac{\mathrm{d}\lambda(t)}{\mathrm{d}t} = -2ax(t) - \lambda(t)\gamma, \tag{S2}$$

with transversality condition $\lambda(1) = 0$, and optimal control characterised by

$$u^*(t) = \frac{\lambda(t)}{2b}. \tag{S3}$$

We set model parameter $\gamma = 0.5$ and pay-off weightings $a = b = 1$, with initial condition $x_0 = 1$ In this case, we are able to solve the control problem analytically. Substituting Equation (S3) into Equation (S1) we can combine with Equation (S2) to form the following system:

$$\begin{bmatrix} \frac{\mathrm{d}x(t)}{\mathrm{d}t} \\ \frac{\mathrm{d}\lambda(t)}{\mathrm{d}t} \end{bmatrix} = \begin{bmatrix} \frac{1}{2} & \frac{1}{2} \\ -2 & -\frac{1}{2} \end{bmatrix} \begin{bmatrix} x(t) \\ \lambda(t) \end{bmatrix}. \tag{S4}$$

This system has complex eigenvalues $e_1 = (\sqrt{3}/2)i$, $e_2 = (-\sqrt{3}/2)i$ and corresponding eigenvectors $v_1 = [(\sqrt{3}i-1)^{-1}, 1]^T$, $v_2 = [(-\sqrt{3}i-1)^{-1}, 1]^T$. Following the standard approach for systems with complex eigenvalues [13], and noting that $v_1 = [-1/4 - \sqrt{3}i/4, 1]^T$ we are able to produce a general solution of the form

$$\begin{bmatrix} x(t) \\ \lambda(t) \end{bmatrix} = C_1 \left( \begin{bmatrix} -\frac{1}{4} \\ 1 \end{bmatrix} \cos\left(\frac{\sqrt{3}t}{2}\right) + \begin{bmatrix} \frac{\sqrt{3}}{4} \\ 0 \end{bmatrix} \sin\left(\frac{\sqrt{3}t}{2}\right) \right) + C_2 \left( - \begin{bmatrix} \frac{\sqrt{3}}{4} \\ 0 \end{bmatrix} \cos\left(\frac{\sqrt{3}t}{2}\right) + \begin{bmatrix} -\frac{1}{4} \\ 1 \end{bmatrix} \sin\left(\frac{\sqrt{3}t}{2}\right) \right). \tag{S5}$$

Evaluating Equation (S5) with the initial condition $x(0) = 1$, and transversality condition $\lambda(1) = 0$, we can solve for the coefficients $C_1$ and $C_2$.

$$\begin{aligned} C_1 &= -4 - \sqrt{3}C_2, \qquad \text{where} \\ C_2 &= \frac{4\cos\left(\frac{\sqrt{3}}{2}\right)}{\sin\left(\frac{\sqrt{3}}{2}\right) - \sqrt{3}\cos\left(\frac{\sqrt{3}}{2}\right)}. \end{aligned}$$

We can then form the complete analytical solution for the optimal state and co-state:

$$\begin{bmatrix} x(t) \\ \lambda(t) \end{bmatrix} = \left( \frac{4\cos\left(\frac{\sqrt{3}}{2}\right)}{\sin\left(\frac{\sqrt{3}}{2}\right) - \sqrt{3}\cos\left(\frac{\sqrt{3}}{2}\right)} \right) \left( \begin{bmatrix} 0 \\ -\sqrt{3} \end{bmatrix} \cos\left(\frac{\sqrt{3}t}{2}\right) + \begin{bmatrix} -1 \\ 1 \end{bmatrix} \sin\left(\frac{\sqrt{3}t}{2}\right) \right) + \left( \begin{bmatrix} 1 \\ -4 \end{bmatrix} \cos\left(\frac{\sqrt{3}t}{2}\right) + \begin{bmatrix} -\sqrt{3} \\ 0 \end{bmatrix} \sin\left(\frac{\sqrt{3}t}{2}\right) \right). \tag{S6}$$

From Equation (S3), recalling that $b = 1$ we can also obtain an analytical expression for the optimal control that depends only on $t$.

$$u^*(t) = \left(\frac{2\cos\left(\frac{\sqrt{3}}{2}\right)}{\sin\left(\frac{\sqrt{3}}{2}\right) - \sqrt{3}\cos\left(\frac{\sqrt{3}}{2}\right)}\right)\left(-\sqrt{3}\cos\left(\frac{\sqrt{3}t}{2}\right) + \sin\left(\frac{\sqrt{3}t}{2}\right)\right) - 2\cos\left(\frac{\sqrt{3}t}{2}\right). \tag{S7}$$

In Figure S1 we plot the exact solutions against the optimal state and control obtained from the FBSM to demonstrate excellent agreement.

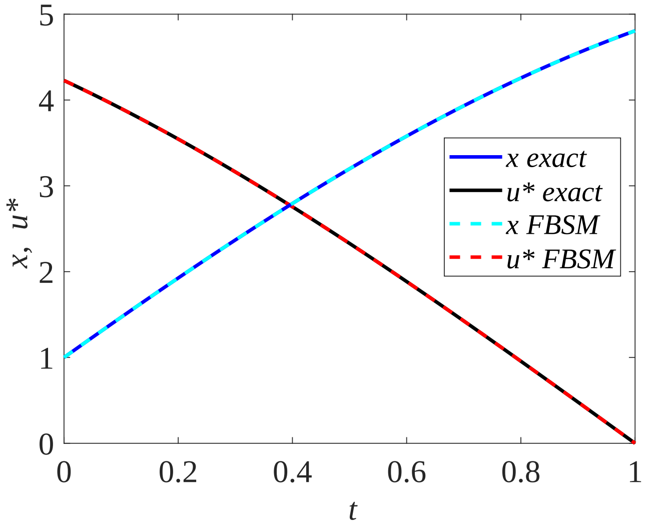


Fig. S1. We observe excellent agreement between the analytical and numerical solutions to the linear continuous control problem. The analytical optimal control is shown in black, with the optimal control obtained via the FBSM shown in red dash. The analytical solution for the state is plotted in blue, with the numerical solution overlaid in cyan dash. These solutions are produced with model parameter $\gamma = 0.5$, time-step $\mathrm{d}t = 3.91 \times 10^{-3}$, over the interval $0 \leq t \leq 1$. The contributions of the state and the control to the pay-off are equally weighted, with $a = b = 1$.

#### 3 Steffensen derivation

In this section, we present the derivation of the multivariate Steffensen method, adapted from the two-variable derivation presented in [9], and then outline the proposed modification for use with accelerating convergence of the FBSM for an iterative process, $X^{(k+1)} = F(X^{(k)})$, where $X^{(k)} = [x_0^{(k)}, x_1^{(k)}, ..., x_N^{(k)}]^{\mathrm{T}} \in \mathbb{R}^{N+1}$ and $F = [f_0, f_1, ..., f_N]^{\mathrm{T}}$. Suppose $S = [s_0, s_1, ..., s_N]^{\mathrm{T}}$ is a solution, such that $S = F(S)$. Denote the error in the $k$th iterations as $E^{(k)} = [e_0^{(k)}, e_1^{(k)}, ..., e_N^{(k)}]^{\mathrm{T}}$ where $E^{(k)} = X^{(K)} - S$, such that

$$e_i^{(k)} = x_i^{(k)} - s_i, \quad i = 0, ..., N.$$

Considering the first error term, and performing a Taylor expansion about $s_0$ we have

$$\begin{aligned} e_0^{(k+1)} &= x_0^{(k+1)} - s_0, \\ &= f_0(x_0^{(k)}, x_1^{(k)}, ..., x_N^{(k)}) - f_0(s_0.s_1, ..., s_N), \\ &= f_0(s_0 + e_0^{(k)}, s_1 + e_1^{(k)}, ..., s_N + e_N^{(k)}) - f_0(s_0.s_1, ..., s_N), \\ &= \frac{\partial f_0(s_0, s_1, ..., s_N)}{\partial x_0} e_0^{(k)} + \frac{\partial f_0(s_0, s_1, ..., s_N)}{\partial x_1} e_1^{(k)} + ... \\ &+ \frac{\partial f_0(s_0, s_1, ..., s_N)}{\partial x_N} e_N^{(k)} + \mathcal{O}(||E||^2). \end{aligned} \tag{S8}$$

Similarly for the $j$th element of $E$, with $j = 1, 2, ..., N$ we have

$$\begin{aligned} e_j^{(k+1)} &= \frac{\partial f_j(s_0, s_1, ..., s_N)}{\partial x_0} e_0^{(k)} + \frac{\partial f_j(s_0, s_1, ..., s_N)}{\partial x_1} e_1^{(k)} + ... \\ &+ \frac{\partial f_j(s_0, s_1, ..., s_N)}{\partial x_N} e_N^{(k)} + \mathcal{O}(||E||^2). \end{aligned} \tag{S9}$$

Recalling the definition of the Jacobian, evaluated at the solution $S$; $J_S$,

$$J_S = \begin{bmatrix} \frac{\partial f_0}{x_0} & \frac{\partial f_0}{x_1} & \cdots & \frac{\partial f_0}{x_N} \\ \frac{\partial f_1}{x_0} & \frac{\partial f_1}{x_1} & \cdots & \frac{\partial f_1}{x_N} \\ \vdots & \vdots & \ddots & \vdots \\ \frac{\partial f_N}{x_0} & \frac{\partial f_N}{x_1} & \cdots & \frac{\partial f_N}{x_N} \end{bmatrix}, \tag{S10}$$

we can express Equations (S8-S9) in matrix form:

$$E^{(k+1)} = J_S E^{(k)} + \mathcal{O}(||E^{(k)}||^2). \tag{S11}$$

In line with the approach for deriving Aitken's formula for a single variable [9], we proceed by assuming that Equation (S11) is true when neglecting the $\mathcal{O}(||E^{(k)}||^2)$ term. Then we have

$$X^{(k+1)} - S = J_S(X^{(k)} - S), \quad k = 0, 1, 2.... \tag{S12}$$

We want to determine $S$, however we do not know $J_S$. Subtracting two consecutive terms of Equation (S12), and denoting $\Delta X^{(k)} = X^{(k+1)} - X^{(k)}$ gives

$$\Delta X^{(k+1)} = J_S \Delta X^{(k)}, \quad k = 0, 1, 2... \tag{S13}$$

Defining $\mathcal{X}^{(k)}$ as a matrix constructed with columns $(X^{(k)}, X^{(k+1)}, ..., X^{(k+N)})$, such that $\mathcal{X}^{(k)} \in \mathbb{R}^{N+1 \times N+1}$;

$$\mathcal{X}^{(k)} = \begin{bmatrix} x_0^{(k)} & x_0^{(k+1)} & \dots & x_0^{(k+N)} \\ x_1^{(k)} & x_1^{(k+1)} & \dots & x_1^{(k+N)} \\ \vdots & \vdots & \ddots & \vdots \\ x_N^{(k)} & x_N^{(k+1)} & \dots & x_N^{(k+N)} \end{bmatrix}, \tag{S14}$$

and $\Delta\mathcal{X}^{(k)} = \mathcal{X}^{(k+1)} - \mathcal{X}^{(k)}$, then we can use Equation (S13) to obtain

$$J_S \Delta\mathcal{X}^{(k)} = \Delta\mathcal{X}^{(k+1)}, \quad k = 0, 1, 2..., \text{ and hence:} \tag{S15}$$

$$J_S = \Delta\mathcal{X}^{(k+1)}(\Delta\mathcal{X}^{(k)})^{-1}. \tag{S16}$$

Rearranging Equation (S12) to solve for $S$, and assuming $(I - J_S)$ is non-singular, where $I$ is the $N + 1 \times N + 1$ identity matrix, we have

$$\begin{aligned} (I - J_S)S &= X^{(k+1)} - J_S X^{(k)}, \\ (I - J_S)S &= (I - J_S)X^{(k)} + \Delta X^{(k)}, \\ S &= X^{(k)} + (I - J_S)^{-1}\Delta X^{(k)}. \end{aligned} \tag{S17}$$

Combining Equation (S17) with Equation (S16), and noting that $(AB)^{-1} = B^{-1}A^{-1}$ then

$$S = X^{(k)} - \Delta\mathcal{X}^{(k)}(\Delta^2\mathcal{X}^{(k)})^{-1}\Delta X^{(k)}. \tag{S18}$$

To arrive at this expression for the true solution, $S$, we have assumed that

Equation (S11) holds when neglecting the $\mathcal{O}(||E^{(k)}||^2)$ term. Though this may not be true, we assume that

$$\hat{X}^{(k)} = X^{(k)} - \Delta\mathcal{X}^{(k)}(\Delta^2\mathcal{X}^{(k)})^{-1}\Delta X^{(k)} \tag{S19}$$

is closer to $S$ than $X^{(k)}$, provided that $\Delta^2(\mathcal{X}^{(k)})^{-1}$ is non-singular. This is the form of Steffensen acceleration for multivariate systems. In the context of accelerating the FBSM, the number of iterations required to construct the $\mathcal{X}^{(k)}$ matrix in Equation (S14) is $\mathcal{O}(N)$. As outlined in the main document, we expect the number of iterations required for the FBSM to converge without acceleration to be fewer than $\mathcal{O}(N)$.

Suppose instead that we chose a number, $m < N+1$, such that when we would form $\mathcal{X}^{(k)}$ in Equation (S14), we instead form a rectangular matrix, $\mathcal{Y}^{(k)}$ of dimension $N+1 \times m+2$:

$$\mathcal{Y}^{(k)} = \begin{bmatrix} x_0^{(k)} & x_0^{(k+1)} & \dots & x_0^{(k+m+1)} \\ x_1^{(k)} & x_1^{(k+1)} & \dots & x_1^{(k+m+1)} \\ \vdots & \vdots & \ddots & \vdots \\ x_N^{(k)} & x_N^{(k+1)} & \dots & x_N^{(k+m+1)} \end{bmatrix}, \tag{S20}$$

We can no longer follow the standard derivation, as Equation (S13) now gives a relationship between rectangular matrices, from which we can no longer isolate $J_S$ via matrix inversion.

$$J_S \Delta \mathcal{Y}^{(k)} = \Delta \mathcal{Y}^{(k+1)}, \quad k = 0, 1, 2... \tag{S21}$$

Instead, we suppose that $\mathcal{X}^{(k)}(\Delta^2 \mathcal{X}^{(k)})^{-1} \Delta X^{(k)}$ in Equation (S19) can be approximated by $\Delta \mathcal{Y}^{(k)}(\Delta^2 \mathcal{Y}^{(k)})^{+} \Delta X^{(k)}$, such that we obtain

$$\hat{X}^{(k)} = X^{(k)} - \Delta \mathcal{Y}^{(k)}(\Delta^2 \mathcal{Y}^{(k)})^{+} \Delta X^{(k)}, \tag{S22}$$

where $(\Delta^2 \mathcal{Y}^{(k)})^{+}$ is the Moore-Penrose pseudoinverse [19]. We will refer to this as the partial Steffensen method.

The Moore-Penrose pseudoinverse is a generalisation of the matrix inverse for singular or non-square matrices. The matrix $A^{+}$ is the (unique) Moore-Penrose pseudoinverse of $A$ if it satisfies these conditions [19]:

1. $AA^{+}A = A$,
2. $A^{+}AA^{+} = A^{+}$,
3. $(AA^{+})^{*} = AA^{+}$,
4. $(A^{+}A)^{*} = A^{+}A$.

Here, $(\cdot)^{*}$ denotes the conjugate transpose. If $A$ is an invertible square matrix, then $A^{+} = A^{-1}$. It can be shown that $X = A^{+}B$ is the least squares solution to $AX = B$, in that it minimises the Frobenius norm; $||A(A^{+}B) - B||_F \leq ||AX - B||_F$, for any choice of $X$ where $AX$ is defined [20,21].

Suppose an $m \times n$ matrix $A$ is expressed in the singular value decomposition

(SVD) form, $A = V^*SU$, where $V$ and $U$ are unitary matrices of dimension $m \times m$ and $n \times n$, respectively. The $m \times n$ matrix $S = \mathrm{diag}(\sigma_1, ..., \sigma_p)$, where $\sigma_i$ are the singular values of $A$, $p = \min(m, n)$, and $\sigma_1 \geq \sigma_2 \geq ... \geq \sigma_p \geq 0$ [7]. The matrix $A^+$ can then be obtained via $A^+ = U^*S^+V$, where $S^+$ is formed by taking the reciprocal of the non-zero diagonal elements of $S$ then transposing the resulting matrix [20].

For numerical implementation of Equation (S22), many popular options for matrix computation such as MATLAB, Julia and Python's NumPy library have built in `pinv()` commands that compute the Moore-Penrose pseudoinverse using SVD [12,15,23]. Further, it may be more efficient to instead use in-built least-squares routines, such as `lsqminnorm()` in MATLAB [15].

#### 4 Acceleration algorithms

In this section we present algorithms for the iterative acceleration methods used in this work. We present algorithms for the Wegstein, Aitken-Steffensen and Anderson methods as applied to systems. In accordance with notation used in the main document, we consider accelerating a fixed point iterative process, $X^{(k+1)} = F(X^{(k)})$, where $X^{(k)} = [x_0^{(k)}, x_1^{(k)}, ...x_N^{(k)}]^{\mathrm{T}}$ is the $k$th iterate for the control, consisting of $N+1$ values, and $F = [f_0, f_1, ...f_N]^{\mathrm{T}}$ is the $N+1$ dimensional operator. We refer to evaluation of $F(X)$ as one *function evaluation*, as this is analogous to the number of iterations in a FBSM process and this is what we seek to reduce. We denote the total number of function evaluations required to achieve convergence as $\mathcal{N}$.

##### *4.1 Wegstein method*

The algorithm for the Wegstein method is adapted from [8,26]. This method requires element-wise computations; with elements denoted by subscript $i$. The element-wise computations are performed on every element, and there is no interaction between elements, such that $i = 1, ..., N+1$, always. This range is not explicitly mentioned in Algorithm 3 as there is no ambiguity.

**Algorithm 3: Wegstein method**

i. From $X^{(0)}$, generate $X^{(1)} = F(X^{(0)})$ and $X^{(2)} = F(X^{(1)})$.

ii. Compute $A_i^{(1)}$ and $q_i^{(1)}$ element-wise; $A_i^{(1)} = \frac{x_i^{(2)} - x_i^{(1)}}{x_i^{(1)} - x_i^{(0)}}$, $q_i^{(1)} = \frac{A_i^{(1)}}{A_i^{(1)} - 1}$, bound $q_i$ if desired.

iii. Update $X^{(2)}$ element-wise; $x_i^{(2)} = q_i^{(1)} x_i^{(1)} + (1 - q_i^{(1)}) f_i(x_i^{(1)})$.

iv. Iterate for $k = 2, 3, ...$, until converged or iteration limit met:

v. Compute $A_i^{(k)}$ and $q_i^{(k)}$ element-wise; $A_i^{(k)} = \frac{f_i(x_i^{(k)}) - f_i(x_i^{(k-1)})}{x_i^{(k)} - x_i^{(k-1)}}$, $q_i^{(k)} = \frac{A_i^{(k)}}{A_i^{(k)} - 1}$, bound $q_i$ if desired.

vi. Update $X^{(k+1)}$ element-wise; $x_i^{(k+1)} = q_i^{(k)} x_i^{(k)} + (1 - q_i^{(k)}) f_i(x_i^{(k)})$.

vii. Check for convergence. If not converged, return to Step iv.

The Wegstein method requires two evaluations of $F$ initially, then one more evaluation of $F$ to update $X^{(2)}$. From Step iv. onward, only one evaluation of $F$ is required per iteration, as the elements $f_i(x_i^{(k-1)})$ used to obtain $A_i^{(k)}$ can be stored from the previous iteration. If $A_i = 1$ in Step ii. or Step v., prescribe a value of $q_i$ to avoid division by zero, for example $q_i = 0$. Aside from the choice of bounds on the $q_i$, variations to the Wegstein method arise by altering how frequently $q$ is updated. In the simplest case, $q$ is determined only once, and not updated in subsequent iterations. Updating $q$ every $n$th iteration can be effective for appropriately chosen $n$, however the choice of $n$ that minimises $\mathcal{N}$ appears to be highly problem dependent and must be empirically determined [22]. As the Wegstein method is implemented element-wise, no expensive matrix operations are performed. However, these element-wise computations ignore interactions between variables and can result in instability for some problems [18].

##### *4.2 Aitken-Steffensen method*

The algorithm for the Steffensen method is adapted from [4,9]. Recall that $X^{(k)} = [x_0^{(k)}, x_1^{(k)}, ...x_N^{(k)}]^{\mathrm{T}}$ is the $k$th iterate, consisting of $N+1$ elements, and $F = [f_0, f_1, ...f_N]^{\mathrm{T}}$ is the $N+1$ dimensional operator of the iterative process. Further, $\Delta X^{(k)} = X^{(k+1)} - X^{(k)}$, $\mathcal{X}^{(k)}$ is a matrix constructed with columns $(X^{(k)}, X^{(k+1)}, ..., X^{(k+N)})$, such that $\mathcal{X}^{(k)}$ is a square matrix of dimension $N+1$, with $\Delta\mathcal{X}^{(k)} = \mathcal{X}^{(k+1)} - \mathcal{X}^{(k)}$, and $\Delta^2\mathcal{X}^{(k)} = \Delta\mathcal{X}^{(k+1)} - \Delta\mathcal{X}^{(k)}$.

**Algorithm 2a: Steffensen method**

i. Iterate for $k = 0, 1, ...,$ until converged or iteration limit met:

ii. From $X^{(k)}$, generate $X^{(k+1)} = F(X^{(k)})$, $X^{(k+2)} = F(X^{(k+1)}), ...,$ $X^{(k+N+2)} = F(X^{(k+N+1)})$.

iii. Define $\Delta X^{(k)} = X^{(k+1)} - X^{(k)}$, $\Delta X^{(k+1)} = X^{(k+2)} - X^{(k+1)}$, ..., $\Delta X^{(k+N+1)} = X^{(k+N+2)} - X^{(k+N+1)}$.

iv. Append results from Step iii. to construct $N+1 \times N+1$ matrices $\Delta\mathcal{X}^{(k)} = (\Delta X^{(k)}, \Delta X^{(k+1)}, ..., \Delta X^{(k+N)})$, and $\Delta\mathcal{X}^{(k+1)} = (\Delta X^{(k+1)}, \Delta X^{(k+2)}, ..., \Delta X^{(k+N+1)})$.

v. Define $\Delta^2\mathcal{X}^{(k)} = \Delta\mathcal{X}^{(k+1)} - \Delta\mathcal{X}^{(k)}$.

vi. Compute $\hat{X}^{(k+1)} = X^{(k)} - \Delta\mathcal{X}^{(k)}(\Delta^2\mathcal{X}^{(k)})^{-1}\Delta X^{(k)}$.

vii. Check for convergence. If not converged, return to Step ii. starting with $X^{(k)} = \hat{X}^{(k+1)}$.

For efficient numerical implementation, the inverse matrix is not formed in Step vi. Rather, we compute $\hat{X}^{(k+1)} = X^{(k)} - \Delta\mathcal{X}^{(k)}P$, where $P$ is obtained by solving the linear system $(\Delta^2\mathcal{X}^{(k)})P = \Delta X^{(k)}$. To recover the Aitken method, continue to compute $\hat{X}^{(k)}$, $k = 1, 2, ...,$ and check for convergence in the $\hat{X}$ series, but do not set $X^{(k)} = \hat{X}^{(k+1)}$ in Step vii; instead use $X^{(k+N+2)}$, the most recently computed $X$ from Step ii.

As noted in §3, we expect the number of iterations required for the FBSM to converge without acceleration to be fewer than the number of FBSM iterations required to construct $\mathcal{X}^{(k)}$. As such, we also present the algorithm for a partial Aitken-Steffensen method outlined in §3, requiring $m+1 \ll N$ function evaluations per iteration.

**Algorithm 2b: Partial Steffensen method**

i. Choose a value $m$ such that $1 \leq m < N+1$.

ii. Iterate for $k = 0, 1, ...,$ until converged or iteration limit met:

iii. From $X^{(k)}$, generate $X^{(k+1)} = F(X^{(k)})$, $X^{(k+2)} = F(X^{(k+1)}), ...,$ $X^{(k+m+1)} = F(X^{(k+m)})$.

iv. Define $\Delta X^{(k)} = X^{(k+1)} - X^{(k)}$, $\Delta X^{(k+1)} = X^{(k+2)} - X^{(k+1)}$, ..., $\Delta X^{(k+m)} = X^{(k+m+1)} - X^{(k+m)}$.

v. Append results from Step iv. to construct $N+1 \times m$ matrices $\Delta\mathcal{Y}^{(k)} = (\Delta X^{(k)}, \Delta X^{(k+1)}, ..., \Delta X^{(k+m-1)})$, and $\Delta\mathcal{Y}^{(k+1)} = (\Delta X^{(k+1)}, \Delta X^{(k+2)}, ..., \Delta X^{(k+m)})$.

vi. Define $\Delta^2\mathcal{Y}^{(k)} = \Delta\mathcal{Y}^{(k+1)} - \Delta\mathcal{Y}^{(k)}$.

vii. Compute $\hat{X}^{(k+1)} = X^{(k)} - \Delta\mathcal{Y}^{(k)}(\Delta^2\mathcal{Y}^{(k)})^{+}\Delta X^{(k)}$, where $(\Delta^2\mathcal{Y}^{(k)})^{+}$ is the Moore-Penrose pseudoinverse.

viii. Check for convergence. If not converged, return to Step ii. starting with $X^{(k)} = \hat{X}^{(k+1)}$.

As noted in §3, for numerical implementation of Step vii. it may be more efficient to use in-built least-squares routines, such as `lsqminnorm()` in MATLAB [15], rather than computing the Moore-Penrose pseudoinverse explicitly via SVD or other means. To recover the partial Aitken method, continue to compute $\hat{X}^{(k)}$, $k = 1, 2, ...,$ and check for convergence in the $\hat{X}$ series, but do not set $X^{(k)} = \hat{X}^{(k+1)}$ in Step vii.; instead use $X^{(k+m+1)}$, the most recently computed $X$ from Step iii.

##### *4.3 Anderson Acceleration*

**Algorithm 4: Anderson acceleration**

i. Choose a value $M \geq 1$, let $m = 1$.

ii. From $X^{(0)}$, generate $X^{(1)} = F(X^{(0)})$ and $X^{(2)} = F(X^{(1)})$.

iii. Define $\Delta X^{(0)} = X^{(1)} - X^{(0)}$, $\Delta X^{(1)} = X^{(2)} - X^{(1)}$, append to form the $N + 1 \times 2$ matrix $\mathcal{G} = (\Delta X^{(0)}, \Delta X^{(1)}) = (G^{(0)}, G^{(1)})$, and compute $\Delta G^{(0)} = \Delta X^{(1)} - \Delta X^{(0)}$. Note that $X^{(2)}$ and $\Delta X^{(1)}$ in this initialisation are overwritten in Step vi. when $k = 1$, by their Anderson algorithm generated counterparts.

iv. Iterate for $k = 1, 2, ...$ until converged or iteration limit met:

v. Solve the least squares problem $\gamma = \arg\min_{\gamma} ||G^{(k)} - \gamma dG^{(k-1)}||$.

vi. Compute $X^{(k+1)} = X^{(k)} + G^{(k)} - (dX^{(k-1)} + dG^{(k-1)})\gamma$, $\Delta X^{(k)} = X^{(k+1)} - X^{(k)}$, $G^{(k+1)} = F(X^{(k+1)}) - X^{(k+1)}$, and $\Delta G^{(k)} = G^{(k+1)} - G^{(k)}$.

vii. `if` $m < M$

Update $\mathcal{G}$,$dX$ and $dG$ by appending the results from Step vi. as the right-most column. Increment $m = m + 1$.

`else`

Update $\mathcal{G}$, $dX$ ad $dG$ by removing the left-most column then appending the results from Step vi. as the right-most column.

viii. Check the condition number of the matrix $dG$; if it exceeds some prescribed tolerance, update $\mathcal{G}$, $dX$ and $dG$ by removing the left-most column. Decrement $m = m - 1$. Repeat Step viii. until the condition number falls below the tolerance or $m = 1$.

ix. Check for convergence. If not converged, return to Step iv.

The algorithm for Anderson Acceleration is adapted from [6,25]. Recall that for a fixed point process where we seek $X = F(X)$, we can define a corresponding root finding problem; $G(X) := F(X) - X = \mathbf{0}$, where $\mathbf{0}$ is the zero column vector of length $N + 1$. For convenience of notation in the algorithm, we denote the difference between iterates; $\Delta X^{(k)} = X^{(k+1)} - X^{(k)}$, residuals; $G^{(k)} = F(X^{(k)}) - X^{(k)}$, and difference between residuals $\Delta G^{(k)} = G^{(k+1)} - G^{(k)}$. Further, $\mathcal{G}^{(k)}$ is a matrix constructed with columns $(G^{(k-m)}, G^{(k-m+1)}, ..., G^{(k)})$, $dX^{(k)}$ is a matrix with columns $(\Delta X^{(k-m)}, \Delta X^{(k-m+1)}, ..., \Delta X^{(k-1)})$, and $dG^{(k)}$ is a matrix with columns $(\Delta G^{(k-m)}, \Delta G^{(k-m+1)}, ..., \Delta G^{(k-1)})$.

The algorithms for Anderson acceleration presented in the literature can vary significantly between sources depending on how the least squares problem is handled, and which commodities are stored and accessed at what stage of the process. The algorithm we present here, along with the associated MATLAB implementation, is one of many approaches, and we recommend resources such as [2,6,25] for a more thorough explanation and further efficiency improvements. We attempt to strike a balance between understandable implementation, efficiency and effectiveness. As we are focussed on using these acceleration techniques to reduce $\mathcal{N}$, we include Step viii. to check the conditioning of $dG$, which is straightforward to implement and can reduce $\mathcal{N}$. However, we do not incorporate the approach of performing QR decomposition on $dG$, that enables more computationally efficient updating and solutions to the least squares problems [25]; as this adds significant complexity to the algorithm and does not further our particular goal in this work of reducing $\mathcal{N}$. For clarity, in Step iii. $\Delta X^{(0)} = G^{(0)}$ and $\Delta X^{(1)} = G^{(1)}$, however this is only true for the initialisation. When the next iterations of these values are calculated in Step vi., $X$ (and hence $\Delta X$) values are updated via the Anderson formula $X^{(k)} + G^{(k)} - (dX^{(k-1)} + dG^{(k-1)})\gamma$, whereas $G$ values are updated based on a function evaluation of the most recently computed $X$.

#### 5 Test nonlinear systems

We test each iterative acceleration technique on three nonlinear systems, before applying the techniques to FBSM problems. Equation (S23) is a $2 \times 2$ system used in [17] to test the Steffensen method. Equations (S24) and (S25) are arbitrarily constructed systems of dimension $3 \times 3$ and $4 \times 4$, respectively. These systems all have at least one real root that can be obtained via fixed point iteration, to use as a benchmark for the effectiveness of each acceleration algorithm. We also present approximate numerical solutions for the other real roots, obtained using `vpasolve()` in MATLAB [16].

The $2 \times 2$ system is:

$$\begin{pmatrix} x_1 \\ x_2 \end{pmatrix} = \begin{pmatrix} \frac{1}{60}(3x_1^3 - 3x_1^2\,x_2 + 6x_1\,x_2^2 + 61.488) \\ \frac{1}{50}(-x_1^3 + 6x_1^2x_2 + 3x_2^3 - 32.496) \end{pmatrix}. \tag{S23}$$

The approximate real solutions to Equation (S23) are:

$$\begin{pmatrix} x_1 \\ x_2 \end{pmatrix} = \begin{pmatrix} 1.4 \\ -1 \end{pmatrix}, \begin{pmatrix} -5.2024 \\ -0.9414 \end{pmatrix}, \begin{pmatrix} -2.2912 \\ -2.9166 \end{pmatrix}, \begin{pmatrix} -1.0562 \\ 4.1194 \end{pmatrix}, \begin{pmatrix} 1.5442 \\ -1.1372 \end{pmatrix}, \begin{pmatrix} 3.2302 \\ 2.3117 \end{pmatrix}, \begin{pmatrix} 3.9247 \\ 1.7874 \end{pmatrix}.$$

The $3 \times 3$ system is:

$$\begin{pmatrix} x_1 \\ x_2 \\ x_3 \end{pmatrix} = \begin{pmatrix} x_1 - \frac{1}{4}(x_1^2 + x_2^2 - 5) \\ x_2 - \frac{1}{2}(x_1x_2 - 2) \\ \frac{1}{3}(x_2 - x_1x_3) \end{pmatrix}. \tag{S24}$$

The approximate real solutions to Equation (S24) are:

$$\begin{pmatrix} x_1 \\ x_2 \\ x_3 \end{pmatrix} = \begin{pmatrix} 1 \\ 2 \\ 0.5 \end{pmatrix}, \begin{pmatrix} -1 \\ -2 \\ -1 \end{pmatrix}, \begin{pmatrix} -2 \\ -1 \\ -1 \end{pmatrix}, \begin{pmatrix} 2 \\ 1 \\ 0.2 \end{pmatrix}.$$

The $4 \times 4$ system is:

$$\begin{pmatrix} x_1 \\ x_2 \\ x_3 \\ x_4 \end{pmatrix} = \begin{pmatrix} \sqrt{4x_1x_4 - x_2^2 + 5} \\ \frac{2}{x_1}(1 - x_2x_4) + \frac{x_3}{10} \\ \frac{1}{3}(x_2 - x_1) + \frac{1}{x_3} \\ \frac{1}{50}(x_1x_3 - \frac{3}{2}x_2x_4) \end{pmatrix}. \tag{S25}$$

The approximate real solutions to Equation (S25) are:

$$\begin{pmatrix} x_1 \\ x_2 \\ x_3 \\ x_4 \end{pmatrix} = \begin{pmatrix} 1.9325 \\ 0.9613 \\ -1.1749 \\ -0.0441 \end{pmatrix}, \begin{pmatrix} 0.9965 \\ 1.9858 \\ -0.8486 \\ -0.0160 \end{pmatrix}, \begin{pmatrix} 1.0039 \\ 2.0205 \\ 1.1837 \\ 0.0224 \end{pmatrix}, \begin{pmatrix} 2.0566 \\ 1.0233 \\ 0.8425 \\ 0.0336 \end{pmatrix}.$$

##### *5.1 Results*

Tables in this section contain the number of function evaluations required for convergence for each system, using each of the acceleration algorithms and for various initial guesses. Recall that we use $\mathcal{N}$ to refer to the number of times the right hand side of the system is evaluated for the iterative procedure to converge. The convergence criteria for all schemes except for Aitken's method is $||F(X) - X|| < 1 \times 10^{-10}$. Where the method fails to converge within 1000 function evaluations, we note *DNC* in the table. For Aitken's method, convergence is determined based on the difference between subsequent results in the Aitken series; $||\hat{X}^{(k+1)} - \hat{X}^{(k)}|| < 1 \times 10^{-10}$.

##### 5.1.1 *Wegstein method*

For the Wegstein method, results are presented for a static value of $q$, and for updating $q$ every $n$th iteration, $n \in \{1, 2, ..., 6\}$. We consider the standard bounds of $-5 \leq q \leq 0$ [3,22], and the unbounded case. The effectiveness varies, with some combinations significantly outperforming fixed point iteration and other combinations requiring far more function evaluations for convergence, or not converging at all. In several cases the Wegstein method converges to a different root to the fixed point iteration, as the Wegstein updating step can cause the iterates to leave the basin of attraction of the solution found via fixed point iteration. Though we are seeking real solutions, the square root term in Equation (S25) can introduce complex values during the iterative process. This creates a challenge when implementing bounds, as $\mathbb{C}$ is not ordered. We handle this by bounding the magnitude of the complex number while maintaining its angle in the complex plane. Code implementing this step is provided on GitHub, though we do not discuss it in detail in this work as it does not impact the acceleration of the FBSM for optimal control problems.

| | | Fixed Point | | Update Wegstein $q$ every $n$th iteration, bounded $-5 \leq q \leq 0$ | | | | | | |
|---|---|---|---|---|---|---|---|---|---|---|
| Eq. | $X^{(0)}$ | Solution | $\mathcal{N}$ | Static | 1 | 2 | 3 | 4 | 5 | 6 |
| S23 | $[0; 0]$ | $[1.4; -1]$ | 167 | 141 | 87 | 74 | 60 | 46 | 42 | 44 |
| | $[1; 2]$ | $[1.4; -1]$ | 167 | 135 | 105 | 55 | 50 | 46 | 53 | DNC |
| | $[1; 1]$ | $[1.4; -1]$ | 167 | 131 | 81 | 66 | 64 | 58 | 67 | 56 |
| | [-1; -1] | $[1.4; -1]$ | 167 | 144 | 91 | 79 | DNC | 50 | 47 | 44 |

| | | Fixed Point | | Update Wegstein $q$ every $n$th iteration, unbounded | | | | | | |
|---|---|---|---|---|---|---|---|---|---|---|
| Eq. | $X^{(0)}$ | Solution | $\mathcal{N}$ | Static | 1 | 2 | 3 | 4 | 5 | 6 |
| S23 | $[0; 0]$ | $[1.4; -1]$ | 167 | 141 | 247 | 32 | 26 | 26 | 27 | 27 |
| | $[1; 2]$ | $[1.4; -1]$ | 167 | 200 | 242 | 26 | 20 | 58 | DNC | 56* |
| | $[1; 1]$ | $[1.4; -1]$ | 167 | 131 | 247 | 26 | DNC | 58* | DNC | DNC |
| | [-1; -1] | $[1.4; -1]$ | 167 | 144 | 251 | 28 | 20 | 22 | 22 | 26 |

* indicates that the procedure converged to another root; $[3.9247; 1.7874]$.

| | | Fixed Point | | Update Wegstein $q$ every $n$th iteration, bounded $-5 \leq q \leq 0$ | | | | | | |
|---|---|---|---|---|---|---|---|---|---|---|
| Eq. | $X^{(0)}$ | Solution | $\mathcal{N}$ | Static | 1 | 2 | 3 | 4 | 5 | 6 |
| S24 | $[0; 0; 0]$ | $[2; 1; 0.2]$ | 56 | DNC | 62 | 62 | 60 | 64 | 64 | 71 |
| | $[1; 2; 3]$ | $[1; 2; 0.5]$ | 23 | 25 | 25 | 25 | 25 | 25 | 25 | 25 |
| | $[1; 1; 1]$ | $[2; 1; 0.2]$ | 57 | 59 | 61 | 59 | 59 | 59 | 59 | 59 |
| | [-1; -1; -1] | $[2; 1; 0.2]$ | 57 | 59 | DNC | 60 | DNC | 61 | 59 | 59 |

| | | Fixed Point | | Update Wegstein $q$ every $n$th iteration, unbounded | | | | | | |
|---|---|---|---|---|---|---|---|---|---|---|
| Eq. | $X^{(0)}$ | Solution | $\mathcal{N}$ | Static | 1 | 2 | 3 | 4 | 5 | 6 |
| S24 | $[0; 0; 0]$ | $[2; 1; 0.2]$ | 56 | DNC | 90 | 21 | 20 | 22 | 24 | 32 |
| | $[1; 2; 3]$ | $[1; 2; 0.5]$ | 23 | 3 | 3 | 3 | 3 | 3 | 3 | 3 |
| | $[1; 1; 1]$ | $[2; 1; 0.2]$ | 57 | 53 | 84 | 22 | 71* | 50 | 28 | 26 |
| | [-1; -1; -1] | $[2; 1; 0.2]$ | 57 | DNC | 93* | 22* | 49 | 22* | 27* | 26* |

* indicates that the procedure converged to another root; $[-2; -1; -1]$.

| | | Fixed Point | | Update Wegstein $q$ every $n$th iteration, bounded $-5 \leq q \leq 0$ | | | | | | |
|---|---|---|---|---|---|---|---|---|---|---|
| Eq. | $X^{(0)}$ | Solution | $\mathcal{N}$ | Static | 1 | 2 | 3 | 4 | 5 | 6 |
| S25 | [1; 2; 3; 4] | [1.9325; | 99 | DNC | 80 | 74 | 98 | 126 | DNC | DNC |
| | [1; 1; 1; 1] | 0.9613; | 100 | 102 | 92 | 102 | 110 | 102 | DNC | 103 |
| | [-1; -1; -1; -1] | -1.1749; | 96 | DNC | 62 | 71 | 86 | 117 | 206 | DNC |
| | | -0.0441] | | | | | | | | |

Results for Equation (S25) exclude the initial condition [0; 0; 0; 0] as the system is not defined for $x_1 = 0$ or $x_3 = 0$. The solution is presented once only for each table concerning Equation (S25), corresponding to all initial values considered.

| | | Fixed Point | | Update Wegstein $q$ every $n$th iteration, unbounded | | | | | | |
|---|---|---|---|---|---|---|---|---|---|---|
| Eq. | $X^{(0)}$ | Solution | $\mathcal{N}$ | Static | 1 | 2 | 3 | 4 | 5 | 6 |
| S25 | [1; 2; 3; 4] | [1.9325; | 99 | DNC | 74 | 74 | 89 | DNC | DNC | DNC |
| | [1; 1; 1; 1] | 0.9613; | 100 | 99 | 99* | 70* | 62* | DNC | DNC | DNC |
| | [-1; -1; -1; -1] | -1.1749; | 96 | DNC | 59 | 70 | 91 | 360 | DNC | DNC |
| | | -0.0441] | | | | | | | | |

*indicates that the procedure converged to another root, [2.0566; 1.0233; 0.8425; 0.0336].

###### 5.1.2 *Aitken-Steffensen methods*

For the partial Aitken and partial Steffensen methods we produce results for $m = 1, 2, ...N$ where $N$ is the size of the system and $m < N$ corresponds to the partial implementation outlined in §4. The Steffensen method required fewer function evaluations than the Aitken method in most cases, though the Aitken method converged for all cases while the Steffensen method did not. In one instance the partial Steffensen method converged to a different root to the fixed point iteration. All implementations of the Aitken and Steffensen methods that converged, did so with fewer function evaluations than the fixed point iteration.

| | | Fixed Point | | Aitken | | Steffensen | |
|---|---|---|---|---|---|---|---|
| Eq. | $X^{(0)}$ | Solution | $\mathcal{N}$ | $m = 1$ | $m = 2$ | $m = 1$ | $m = 2$ |
| S23 | [0; 0] | $[1.4; -1]$ | 167 | 92 | 66 | 61 | 16 |
| | [1; 2] | $[1.4; -1]$ | 167 | 92 | 63 | 55 | 19 |
| | [1; 1] | $[1.4; -1]$ | 167 | 92 | 69 | 71 | 19 |
| | [-1; -1] | $[1.4; -1]$ | 167 | 92 | 66 | 59 | 19 |

| | | Fixed Point | | Aitken, $m =$ | | | Steffensen, $m =$ | | |
|---|---|---|---|---|---|---|---|---|---|
| Eq. | $X^{(0)}$ | Solution | $\mathcal{N}$ | 1 | 2 | 3 | 1 | 2 | 3 |
| S24 | [0; 0; 0] | [2; 1; 0.2] | 56 | 38 | 39 | 28 | 47 | 22 | 17 |
| | [1; 2; 3] | [1; 2; 0.5] | 23 | 4 | 6 | 8 | 3 | 4 | 5 |
| | [1; 1; 1] | [2; 1; 0.2] | 57 | 38 | 36 | 28 | 43 | 25 | 17 |
| | [-1; -1; -1] | [2; 1; 0.2] | 57 | 38 | 39 | 28 | DNC | DNC | DNC |

| | | Fixed Point | | Aitken, $m =$ | | | | Steffensen, $m =$ | | | |
|---|---|---|---|---|---|---|---|---|---|---|---|
| Eq. | $X^{(0)}$ | Solution | $\mathcal{N}$ | 1 | 2 | 3 | 4 | 1 | 2 | 3 | 4 |
| S25 | [1; 2; 3; 4] | [1.9325; 0.9613; -1.1749; -0.0441] | 99 | 50 | 48 | 40 | 35 | 73 | 76 | 53 | 21 |
| | [1; 1; 1; 1] | | 100 | 50 | 45 | 40 | 35 | DNC | 79* | 61 | 36 |
| | [-1; -1; -1; -1] | | 96 | 46 | 42 | 36 | 30 | 72 | 57 | 32 | 15 |

* indicates that the procedure converged to another root, [2.0566; 1.0233; 0.8425; 0.0336].

##### *5.1.3 Anderson Acceleration*

For Anderson acceleration we produce results for $M \in \{1, 2, ..., 5\}$, with a tolerance of $1 \times 10^{10}$ when checking the conditioning of the matrix. Convergence was achieved in all implementations of Anderson acceleration, though several cases converged to a different root to the fixed point iteration. Of all the methods considered, Anderson acceleration produced the most consistent reduction in function evaluations relative to the fixed point iteration when applied to these test nonlinear systems.

| | | Fixed Point | | Anderson | | | | |
|---|---|---|---|---|---|---|---|---|
| Eq. | $X^{(0)}$ | Solution | $\mathcal{N}$ | $M=1$ | $M=2$ | $M=3$ | $M=4$ | $M=5$ |
| S23 | $[0;0]$ | $[1.4;-1]$ | 167 | 11 | 15 | 13 | 15 | 16 |
| | $[1;2]$ | $[1.4;-1]$ | 167 | 13 | 12 | 13 | 16 | 17 |
| | $[1;1]$ | $[1.4;-1]$ | 167 | 12 | 12 | 14 | 16 | 17 |
| | [-1;-1] | $[1.4;-1]$ | 167 | 12 | 12 | 14 | 16 | 19 |

| | | Fixed Point | | Anderson | | | | |
|---|---|---|---|---|---|---|---|---|
| Eq. | $X^{(0)}$ | Solution | $\mathcal{N}$ | $M=1$ | $M=2$ | $M=3$ | $M=4$ | $M=5$ |
| S24 | $[0;0;0]$ | $[2;1;0.2]$ | 56 | 28 | 18 | 12 | 14 | 14 |
| | $[1;2;3]$ | $[1;2;0.5]$ | 23 | 3 | 3 | 3 | 3 | 3 |
| | $[1;1;1]$ | $[2;1;0.2]$ | 57 | 24 | 14 | 12 | 12 | 13 |
| | [-1;-1;-1] | $[2;1;0.2]$ | 57 | 53* | 30* | 22 | 35* | 27* |

* indicates that the procedure converged to another root, [-1;-2;-1].

| | | Fixed Point | | Anderson | | | | |
|---|---|---|---|---|---|---|---|---|
| Eq. | $X^{(0)}$ | Solution | $\mathcal{N}$ | $M=1$ | $M=2$ | $M=3$ | $M=4$ | $M=5$ |
| S25 | $[1;2;3;4]$ | [1.9325; | 99 | 32 | 29 | 21 | 22 | 27 |
| | $[1;1;1;1]$ | 0.9613; | 100 | 37* | 27* | 14* | 14* | 16* |
| | [-1; -1; -1; -1] | -1.1749; | 96 | 25 | 19 | 12 | 13 | 13 |
| | | -0.0441] | | | | | | |

* indicates that the procedure converged to another root, $[2.0566; 1.0233; 0.8425; 0.0336]$.

### 6 Control results

In this section we present results for the acceleration algorithms applied to control problems, for a wide range of tuning parameters. For Wegstein's method we vary $\omega$ and the frequency with which $q$ is updated. For each problem we select a bounding on $q$ that works reasonably, though we do not attempt to find the optimal bounds. For the Aitken and Steffensen methods we vary $\omega$ and $m$. For Anderson acceleration we vary $\omega$ and $M$, with a tolerance of $1 \times 10^{10}$ when checking the conditioning of the matrix. We do not vary or attempt to optimise this tolerance. Each value in the tables correspond to $\mathcal{N}$, the number of function evaluations required for convergence. Simulations are terminated when $\mathcal{N}$ reaches 100; any value in the tables that is 100 or greater corresponds to a combination of tuning parameters that did not yield convergence within this specified maximum. This does not necessarily imply that this combination would not have converged if additional function evaluations were performed. For the linear problems, we do not vary $\omega$ as the FBSM with no acceleration converges with minimum $\mathcal{N}$ when $\omega = 0$.

A heatmap is applied to each table, with colours scaled relative to the result from the FBSM with the best tuning but without acceleration. Recall that with the tuning that minimises $\mathcal{N}$, the FBSM with no acceleration requires $\mathcal{N} = 57$ for the linear continuous control problem, $\mathcal{N} = 8$ for the linear bang-bang control problem, $\mathcal{N} = 38$ for the AML continuous control problem, and $\mathcal{N} = 34$ for the AML bang-bang control problem. Acceleration results that reflect a reduction in $\mathcal{N}$ relative to the these FBSM results are shaded in the green spectrum, while worse performance is shaded in the red spectrum. The midpoint of the colour spectra, yellow, corresponds to the FBSM result with the best tuning, without acceleration. In the main document we use robustness to refer to the ability of a method to reduce $\mathcal{N}$ over a range of tuning parameters. Visually, tables with large groups of green shaded cells indicate robustness, while isolated green cells surrounded by orange-red cells suggest a lack of robustness.

For the AML bang-bang control problem with the Aitken method, for values of $\omega \leq 0.35$, we observe apparent convergence to controls that are not bang-bang. The iterative procedure terminates as the convergence criteria is met; however the resulting controls contain intermediate values between the lower and upper bounds. Similarly for the AML continuous control problem with the Aitken method, for values of $\omega \leq 0.35$, we observe apparent convergence. However, explicitly calculating the pay-off associated with these controls, and comparing it to the pay-off associated with the control obtained via the standard FBSM, indicates that the controls obtained via the Aitken method for $\omega \leq 0.35$ are not optimal. These sections of the partial Aitken method tables have been denoted as $\mathcal{N} = 100$, to indicate a failure to converge to the optimal control; regardless of the number of iterations taken to achieve the apparent convergence to controls that are not optimal.

We note that implementation of Anderson Acceleration involves computing condition numbers of matrices. Matrices that are ill-conditioned (indicated by a large condition number), are close to singular, such that significant numerical error can arise when computing the inverse, or obtaining the solution of a corresponding linear system of equations [10]. It is known that for ill-conditioned matrices, computation of the condition number can itself be highly sensitive [10]. Computing condition numbers is commonly impacted by underflow and overflow, or rounding errors [5,10]. For this reason, users attempting to reproduce results of the Anderson Acceleration method using different software or hardware may find that in some instances convergence is achieved with a different $\mathcal{N}$ to what is indicated in the tables, depending on whether the estimated condition number at each iteration suggests that the matrix is ill-conditioned. For a thorough discussion of condition numbers and issues arising from floating point arithmetic we direct readers to [11].

##### 6.1 Linear continuous control problem

Wegstein, updating $q$ every $n$th iteration, $-2 \le q \le 0$

| $\omega$ | $n$=1 | $n$=2 | $n$=3 | $n$=4 | $n$=5 | $n$=6 | $n$=7 | $n$=8 | $n$=9 | $n$=10 |
|---|---|---|---|---|---|---|---|---|---|---|
| 0 | 24 | 32 | 26 | 23 | 27 | 38 | 44 | 42 | 47 | 62 |

Partial Aitken

| $\omega$ | $m$=1 | $m$=2 | $m$=3 | $m$=4 | $m$=5 | $m$=6 | $m$=7 | $m$=8 | $m$=9 | $m$=10 |
|---|---|---|---|---|---|---|---|---|---|---|
| 0 | 12 | 12 | 12 | 15 | 18 | 14 | 16 | 18 | 20 | 22 |

Partial Steffensen

| $\omega$ | $m$=1 | $m$=2 | $m$=3 | $m$=4 | $m$=5 | $m$=6 | $m$=7 | $m$=8 | $m$=9 | $m$=10 |
|---|---|---|---|---|---|---|---|---|---|---|
| 0 | 31 | 16 | 13 | 11 | 13 | 8 | 9 | 10 | 11 | 12 |

Anderson

| $\omega$ | $M$=1 | $M$=2 | $M$=3 | $M$=4 | $M$=5 | $M$=6 | $M$=7 | $M$=8 | $M$=9 | $M$=10 |
|---|---|---|---|---|---|---|---|---|---|---|
| 0 | 12 | 9 | 8 | 7 | 7 | 7 | 7 | 7 | 7 | 7 |

##### 6.2 Linear bang-bang control problem

Wegstein, unbounded

| $\omega$ | $n$=1 | $n$=2 | $n$=3 | $n$=4 | $n$=5 | $n$=6 | $n$=7 | $n$=8 | $n$=9 | $n$=10 |
|---|---|---|---|---|---|---|---|---|---|---|
| 0 | 9 | 9 | 9 | 9 | 9 | 9 | 9 | 9 | 9 | 9 |

Partial Aitken

| $\omega$ | $m$=1 | $m$=2 | $m$=3 | $m$=4 | $m$=5 | $m$=6 | $m$=7 | $m$=8 | $m$=9 | $m$=10 |
|---|---|---|---|---|---|---|---|---|---|---|
| 0 | 10 | 12 | 12 | 15 | 18 | 21 | 16 | 18 | 20 | 22 |

Partial Steffensen

| $\omega$ | $m$=1 | $m$=2 | $m$=3 | $m$=4 | $m$=5 | $m$=6 | $m$=7 | $m$=8 | $m$=9 | $m$=10 |
|---|---|---|---|---|---|---|---|---|---|---|
| 0 | 29 | 19 | 17 | 16 | 13 | 15 | 9 | 10 | 11 | 12 |

Anderson

| $\omega$ | $M$=1 | $M$=2 | $M$=3 | $M$=4 | $M$=5 | $M$=6 | $M$=7 | $M$=8 | $M$=9 | $M$=10 |
|---|---|---|---|---|---|---|---|---|---|---|
| 0 | 11 | 14 | 17 | 19 | 21 | 22 | 21 | 24 | 26 | 28 |

#### 6.3 *AML continuous control problem with the Wegstein method*

Wegstein, updating $q$ every $n$th iteration, $-1 \leq q \leq 1$

| $\omega$ | $n$=1 | $n$=2 | $n$=3 | $n$=4 | $n$=5 | $n$=6 | $n$=7 | $n$=8 | $n$=9 | $n$=10 |
|---|---|---|---|---|---|---|---|---|---|---|
| 0.95 | 68 | 60 | 62 | 87 | 100 | 100 | 100 | 100 | 100 | 100 |
| 0.90 | 61 | 54 | 67 | 54 | 57 | 100 | 100 | 100 | 100 | 100 |
| 0.85 | 61 | 52 | 62 | 83 | 72 | 77 | 100 | 100 | 100 | 100 |
| 0.80 | 62 | 56 | 59 | 73 | 68 | 100 | 100 | 100 | 100 | 100 |
| 0.75 | 68 | 56 | 53 | 89 | 62 | 100 | 95 | 100 | 100 | 100 |
| 0.70 | 57 | 54 | 71 | 62 | 72 | 92 | 70 | 100 | 100 | 74 |
| 0.65 | 57 | 53 | 55 | 61 | 62 | 90 | 65 | 100 | 56 | 100 |
| 0.60 | 57 | 53 | 56 | 65 | 67 | 36 | 100 | 100 | 100 | 100 |
| 0.55 | 63 | 58 | 50 | 62 | 42 | 26 | 59 | 66 | 100 | 100 |
| 0.50 | 64 | 58 | 53 | 88 | 47 | 47 | 79 | 100 | 100 | 92 |
| 0.45 | 68 | 59 | 41 | 78 | 74 | 74 | 90 | 100 | 100 | 63 |
| 0.40 | 54 | 64 | 50 | 90 | 72 | 100 | 100 | 100 | 76 | 100 |
| 0.35 | 72 | 63 | 48 | 77 | 67 | 66 | 93 | 100 | 100 | 100 |
| 0.30 | 62 | 68 | 67 | 55 | 72 | 86 | 100 | 45 | 100 | 100 |
| 0.25 | 63 | 72 | 60 | 74 | 100 | 100 | 100 | 100 | 100 | 100 |
| 0.20 | 57 | 65 | 56 | 75 | 87 | 92 | 86 | 100 | 97 | 100 |
| 0.15 | 63 | 65 | 68 | 86 | 73 | 100 | 86 | 100 | 100 | 100 |
| 0.10 | 64 | 66 | 66 | 71 | 77 | 100 | 100 | 100 | 100 | 100 |
| 0.05 | 64 | 55 | 71 | 72 | 77 | 100 | 100 | 100 | 100 | 94 |
| 0.00 | 60 | 54 | 59 | 70 | 92 | 100 | 100 | 100 | 100 | 100 |

| $\omega$ | Aitken $m$=1 | $m$=2 | $m$=3 | $m$=4 | $m$=5 | $m$=6 | $m$=7 | $m$=8 | $m$=9 | $m$=10 |
|---|---|---|---|---|---|---|---|---|---|---|
| 0.95 | 100 | 102 | 100 | 100 | 102 | 105 | 104 | 108 | 100 | 110 |
| 0.90 | 100 | 102 | 100 | 100 | 102 | 105 | 104 | 108 | 100 | 110 |
| 0.85 | 100 | 102 | 96 | 100 | 96 | 91 | 88 | 90 | 90 | 88 |
| 0.80 | 86 | 81 | 72 | 75 | 60 | 63 | 64 | 72 | 60 | 55 |
| 0.75 | 68 | 63 | 56 | 55 | 54 | 49 | 48 | 54 | 50 | 44 |
| 0.70 | 56 | 54 | 48 | 45 | 42 | 42 | 40 | 45 | 40 | 44 |
| 0.65 | 48 | 45 | 40 | 35 | 36 | 35 | 32 | 36 | 40 | 44 |
| 0.60 | 42 | 39 | 36 | 35 | 36 | 35 | 32 | 36 | 40 | 44 |
| 0.55 | 36 | 39 | 36 | 35 | 30 | 35 | 32 | 36 | 40 | 44 |
| 0.50 | 36 | 33 | 32 | 30 | 30 | 35 | 32 | 36 | 40 | 33 |
| 0.45 | 36 | 33 | 32 | 35 | 36 | 35 | 32 | 36 | 40 | 44 |
| 0.40 | 78 | 60 | 40 | 40 | 42 | 42 | 48 | 54 | 50 | 55 |
| 0.35 | 100 | 100 | 100 | 100 | 100 | 100 | 100 | 100 | 100 | 100 |
| 0.30 | 100 | 100 | 100 | 100 | 100 | 100 | 100 | 100 | 100 | 100 |
| 0.25 | 100 | 100 | 100 | 100 | 100 | 100 | 100 | 100 | 100 | 100 |
| 0.20 | 100 | 100 | 100 | 100 | 100 | 100 | 100 | 100 | 100 | 100 |
| 0.15 | 100 | 100 | 100 | 100 | 100 | 100 | 100 | 100 | 100 | 100 |
| 0.10 | 100 | 100 | 100 | 100 | 100 | 100 | 100 | 100 | 100 | 100 |
| 0.05 | 100 | 100 | 100 | 100 | 100 | 100 | 100 | 100 | 100 | 100 |
| 0.00 | 100 | 100 | 100 | 100 | 100 | 100 | 100 | 100 | 100 | 100 |

| $\omega$ | Steffensen $m$=1 | $m$=2 | $m$=3 | $m$=4 | $m$=5 | $m$=6 | $m$=7 | $m$=8 | $m$=9 | $m$=10 |
|---|---|---|---|---|---|---|---|---|---|---|
| 0.95 | 61 | 28 | 25 | 26 | 25 | 29 | 33 | 28 | 31 | 45 |
| 0.90 | 53 | 31 | 21 | 21 | 25 | 22 | 25 | 28 | 31 | 34 |
| 0.85 | 47 | 28 | 21 | 21 | 19 | 22 | 25 | 28 | 31 | 34 |
| 0.80 | 61 | 28 | 25 | 21 | 25 | 22 | 25 | 28 | 31 | 34 |
| 0.75 | 53 | 34 | 25 | 21 | 25 | 22 | 25 | 28 | 31 | 34 |
| 0.70 | 49 | 37 | 25 | 21 | 25 | 22 | 25 | 28 | 31 | 34 |
| 0.65 | 53 | 34 | 25 | 21 | 25 | 22 | 25 | 28 | 31 | 34 |
| 0.60 | 61 | 34 | 25 | 21 | 19 | 22 | 25 | 28 | 31 | 34 |
| 0.55 | 67 | 34 | 25 | 21 | 19 | 22 | 25 | 28 | 31 | 34 |
| 0.50 | 69 | 31 | 25 | 21 | 19 | 22 | 25 | 28 | 31 | 23 |
| 0.45 | 69 | 31 | 25 | 21 | 25 | 22 | 25 | 28 | 31 | 34 |
| 0.40 | 69 | 31 | 25 | 21 | 25 | 22 | 25 | 28 | 31 | 34 |
| 0.35 | 53 | 31 | 25 | 26 | 31 | 29 | 33 | 37 | 41 | 45 |
| 0.30 | 67 | 34 | 29 | 31 | 37 | 36 | 49 | 46 | 51 | 56 |
| 0.25 | 65 | 31 | 33 | 46 | 55 | 71 | 65 | 82 | 101 | 100 |
| 0.20 | 65 | 34 | 37 | 56 | 37 | 106 | 81 | 100 | 101 | 100 |
| 0.15 | 73 | 31 | 45 | 71 | 37 | 106 | 49 | 100 | 101 | 100 |
| 0.10 | 51 | 43 | 57 | 76 | 79 | 106 | 105 | 100 | 101 | 100 |
| 0.05 | 67 | 55 | 73 | 86 | 103 | 106 | 105 | 100 | 101 | 100 |
| 0.00 | 71 | 64 | 101 | 61 | 103 | 106 | 105 | 100 | 101 | 100 |

#### 6.6 AML continuous control problem with Anderson acceleration

| $\omega$ | Anderson $M$=1 | $M$=2 | $M$=3 | $M$=4 | $M$=5 | $M$=6 | $M$=7 | $M$=8 | $M$=9 | $M$=10 |
|---|---|---|---|---|---|---|---|---|---|---|
| 0.95 | 100 | 100 | 100 | 33 | 34 | 32 | 31 | 28 | 28 | 27 |
| 0.90 | 100 | 40 | 27 | 25 | 24 | 26 | 23 | 20 | 20 | 20 |
| 0.85 | 62 | 45 | 22 | 22 | 20 | 17 | 17 | 18 | 19 | 19 |
| 0.80 | 53 | 34 | 26 | 23 | 24 | 21 | 22 | 19 | 21 | 22 |
| 0.75 | 44 | 30 | 28 | 29 | 24 | 26 | 33 | 27 | 34 | 31 |
| 0.70 | 40 | 31 | 36 | 30 | 35 | 33 | 54 | 44 | 47 | 55 |
| 0.65 | 33 | 26 | 29 | 58 | 30 | 35 | 50 | 41 | 57 | 54 |
| 0.60 | 30 | 26 | 27 | 28 | 32 | 45 | 71 | 45 | 46 | 71 |
| 0.55 | 33 | 25 | 26 | 26 | 32 | 32 | 41 | 40 | 42 | 54 |
| 0.50 | 32 | 29 | 27 | 28 | 30 | 28 | 30 | 39 | 38 | 37 |
| 0.45 | 33 | 25 | 27 | 25 | 28 | 27 | 26 | 29 | 30 | 28 |
| 0.40 | 31 | 23 | 28 | 27 | 38 | 29 | 36 | 31 | 41 | 41 |
| 0.35 | 28 | 24 | 23 | 26 | 30 | 30 | 31 | 39 | 36 | 38 |
| 0.30 | 26 | 24 | 25 | 29 | 27 | 29 | 33 | 33 | 45 | 51 |
| 0.25 | 23 | 21 | 25 | 28 | 38 | 33 | 65 | 49 | 58 | 56 |
| 0.20 | 22 | 21 | 25 | 28 | 27 | 57 | 32 | 46 | 87 | 75 |
| 0.15 | 20 | 21 | 24 | 32 | 39 | 52 | 34 | 44 | 45 | 53 |
| 0.10 | 22 | 19 | 24 | 32 | 35 | 36 | 31 | 42 | 69 | 50 |
| 0.05 | 24 | 22 | 30 | 34 | 37 | 45 | 47 | 74 | 75 | 48 |
| 0.00 | 26 | 25 | 27 | 80 | 40 | 56 | 64 | 43 | 100 | 94 |

Wegstein, updating $q$ every $n$th iteration, $-1 \leq q \leq 1$

| $\omega$ | $n$=1 | $n$=2 | $n$=3 | $n$=4 | $n$=5 | $n$=6 | $n$=7 | $n$=8 | $n$=9 | $n$=10 |
|---|---|---|---|---|---|---|---|---|---|---|
| 0.95 | 31 | 32 | 11 | 14 | 17 | 20 | 23 | 26 | 29 | 32 |
| 0.90 | 30 | 30 | 29 | 30 | 17 | 20 | 23 | 26 | 29 | 32 |
| 0.85 | 30 | 30 | 29 | 14 | 17 | 20 | 23 | 26 | 29 | 32 |
| 0.80 | 31 | 20 | 26 | 26 | 17 | 20 | 23 | 26 | 29 | 32 |
| 0.75 | 31 | 24 | 17 | 14 | 17 | 20 | 23 | 26 | 29 | 32 |
| 0.70 | 30 | 24 | 11 | 46 | 27 | 20 | 23 | 42 | 29 | 32 |
| 0.65 | 28 | 28 | 26 | 42 | 37 | 44 | 37 | 42 | 29 | 32 |
| 0.60 | 30 | 26 | 20 | 50 | 27 | 44 | 44 | 26 | 29 | 32 |
| 0.55 | 30 | 26 | 32 | 22 | 17 | 20 | 23 | 26 | 29 | 32 |
| 0.50 | 31 | 30 | 32 | 14 | 17 | 32 | 23 | 26 | 29 | 32 |
| 0.45 | 31 | 28 | 32 | 34 | 17 | 32 | 23 | 26 | 29 | 32 |
| 0.40 | 27 | 28 | 38 | 46 | 27 | 32 | 23 | 26 | 29 | 32 |
| 0.35 | 25 | 36 | 38 | 46 | 52 | 56 | 37 | 26 | 29 | 32 |
| 0.30 | 28 | 30 | 41 | 46 | 52 | 50 | 44 | 50 | 56 | 62 |
| 0.25 | 29 | 32 | 35 | 38 | 42 | 50 | 44 | 50 | 20 | 22 |
| 0.20 | 32 | 34 | 35 | 34 | 32 | 14 | 16 | 18 | 20 | 22 |
| 0.15 | 27 | 30 | 35 | 34 | 27 | 14 | 16 | 26 | 11 | 12 |
| 0.10 | 27 | 28 | 35 | 30 | 27 | 14 | 16 | 18 | 29 | 12 |
| 0.05 | 28 | 30 | 29 | 26 | 37 | 14 | 23 | 10 | 11 | 12 |
| 0.00 | 33 | 32 | 35 | 30 | 37 | 26 | 9 | 10 | 11 | 12 |

| $\omega$ | Aitken $m$=1 | $m$=2 | $m$=3 | $m$=4 | $m$=5 | $m$=6 | $m$=7 | $m$=8 | $m$=9 | $m$=10 |
|---|---|---|---|---|---|---|---|---|---|---|
| 0.95 | 86 | 87 | 88 | 90 | 102 | 105 | 104 | 108 | 100 | 110 |
| 0.90 | 42 | 48 | 52 | 55 | 54 | 56 | 64 | 63 | 70 | 66 |
| 0.85 | 32 | 33 | 36 | 40 | 42 | 42 | 48 | 45 | 50 | 55 |
| 0.80 | 24 | 24 | 28 | 30 | 30 | 35 | 40 | 36 | 40 | 44 |
| 0.75 | 18 | 21 | 20 | 25 | 24 | 28 | 24 | 27 | 30 | 33 |
| 0.70 | 16 | 18 | 20 | 20 | 24 | 21 | 24 | 27 | 30 | 33 |
| 0.65 | 12 | 15 | 16 | 15 | 18 | 21 | 24 | 27 | 20 | 22 |
| 0.60 | 10 | 12 | 12 | 15 | 18 | 21 | 16 | 18 | 20 | 22 |
| 0.55 | 10 | 12 | 16 | 15 | 18 | 21 | 24 | 27 | 30 | 33 |
| 0.50 | 8 | 9 | 12 | 15 | 12 | 14 | 16 | 18 | 20 | 22 |
| 0.45 | 10 | 12 | 12 | 15 | 18 | 21 | 16 | 18 | 20 | 22 |
| 0.40 | 14 | 15 | 16 | 20 | 18 | 21 | 24 | 27 | 30 | 22 |
| 0.35 | 100 | 100 | 100 | 100 | 100 | 100 | 100 | 100 | 100 | 100 |
| 0.30 | 100 | 100 | 100 | 100 | 100 | 100 | 100 | 100 | 100 | 100 |
| 0.25 | 100 | 100 | 100 | 100 | 100 | 100 | 100 | 100 | 100 | 100 |
| 0.20 | 100 | 100 | 100 | 100 | 100 | 100 | 100 | 100 | 100 | 100 |
| 0.15 | 100 | 100 | 100 | 100 | 100 | 100 | 100 | 100 | 100 | 100 |
| 0.10 | 100 | 100 | 100 | 100 | 100 | 100 | 100 | 100 | 100 | 100 |
| 0.05 | 100 | 100 | 100 | 100 | 100 | 100 | 100 | 100 | 100 | 100 |
| 0.00 | 100 | 100 | 100 | 100 | 100 | 100 | 100 | 100 | 100 | 100 |

| $\omega$ | Steffensen $m$=1 | $m$=2 | $m$=3 | $m$=4 | $m$=5 | $m$=6 | $m$=7 | $m$=8 | $m$=9 | $m$=10 |
|---|---|---|---|---|---|---|---|---|---|---|
| 0.95 | 101 | 100 | 101 | 66 | 103 | 106 | 105 | 100 | 101 | 100 |
| 0.90 | 51 | 100 | 41 | 41 | 73 | 64 | 33 | 64 | 71 | 78 |
| 0.85 | 33 | 100 | 101 | 41 | 43 | 43 | 49 | 46 | 51 | 45 |
| 0.80 | 29 | 34 | 29 | 26 | 37 | 36 | 33 | 37 | 41 | 34 |
| 0.75 | 29 | 28 | 17 | 26 | 25 | 22 | 33 | 28 | 21 | 34 |
| 0.70 | 23 | 22 | 21 | 21 | 19 | 22 | 17 | 19 | 21 | 23 |
| 0.65 | 21 | 19 | 17 | 16 | 19 | 15 | 17 | 19 | 11 | 12 |
| 0.60 | 21 | 13 | 13 | 11 | 13 | 15 | 9 | 10 | 11 | 12 |
| 0.55 | 19 | 16 | 13 | 11 | 13 | 15 | 17 | 19 | 21 | 23 |
| 0.50 | 19 | 16 | 17 | 11 | 7 | 8 | 9 | 10 | 11 | 12 |
| 0.45 | 21 | 16 | 17 | 16 | 13 | 15 | 9 | 10 | 11 | 12 |
| 0.40 | 17 | 19 | 17 | 21 | 25 | 22 | 17 | 19 | 21 | 12 |
| 0.35 | 17 | 16 | 17 | 21 | 19 | 22 | 25 | 28 | 31 | 23 |
| 0.30 | 19 | 16 | 17 | 16 | 19 | 22 | 25 | 28 | 31 | 34 |
| 0.25 | 19 | 16 | 17 | 16 | 19 | 22 | 25 | 28 | 31 | 34 |
| 0.20 | 19 | 13 | 17 | 21 | 19 | 22 | 25 | 28 | 31 | 34 |
| 0.15 | 19 | 13 | 17 | 21 | 25 | 22 | 25 | 28 | 31 | 34 |
| 0.10 | 19 | 16 | 17 | 21 | 25 | 29 | 33 | 37 | 31 | 34 |
| 0.05 | 19 | 19 | 17 | 26 | 25 | 29 | 33 | 37 | 41 | 45 |
| 0.00 | 17 | 22 | 21 | 26 | 25 | 36 | 33 | 46 | 41 | 56 |

| $\omega$ | Anderson $M$=1 | $M$=2 | $M$=3 | $M$=4 | $M$=5 | $M$=6 | $M$=7 | $M$=8 | $M$=9 | $M$=10 |
|---|---|---|---|---|---|---|---|---|---|---|
| 0.95 | 97 | 100 | 100 | 100 | 100 | 100 | 100 | 100 | 100 | 100 |
| 0.90 | 93 | 62 | 100 | 100 | 100 | 100 | 98 | 100 | 100 | 100 |
| 0.85 | 72 | 71 | 56 | 49 | 62 | 62 | 100 | 91 | 83 | 100 |
| 0.80 | 55 | 56 | 68 | 50 | 68 | 100 | 46 | 60 | 56 | 100 |
| 0.75 | 41 | 40 | 50 | 58 | 48 | 76 | 50 | 70 | 57 | 61 |
| 0.70 | 52 | 23 | 40 | 43 | 36 | 76 | 52 | 45 | 46 | 41 |
| 0.65 | 18 | 20 | 37 | 28 | 38 | 41 | 45 | 35 | 54 | 100 |
| 0.60 | 26 | 24 | 25 | 42 | 30 | 42 | 33 | 32 | 31 | 34 |
| 0.55 | 24 | 19 | 29 | 26 | 28 | 23 | 27 | 26 | 30 | 31 |
| 0.50 | 18 | 24 | 17 | 34 | 28 | 63 | 32 | 69 | 39 | 100 |
| 0.45 | 23 | 24 | 26 | 31 | 28 | 29 | 42 | 56 | 47 | 17 |
| 0.40 | 17 | 25 | 32 | 38 | 39 | 29 | 38 | 43 | 52 | 80 |
| 0.35 | 24 | 21 | 22 | 32 | 38 | 20 | 17 | 17 | 17 | 17 |
| 0.30 | 23 | 23 | 32 | 23 | 42 | 45 | 20 | 19 | 19 | 19 |
| 0.25 | 27 | 28 | 47 | 30 | 38 | 44 | 41 | 33 | 23 | 19 |
| 0.20 | 22 | 27 | 26 | 51 | 47 | 78 | 35 | 41 | 39 | 25 |
| 0.15 | 25 | 24 | 26 | 28 | 29 | 31 | 92 | 39 | 26 | 19 |
| 0.10 | 25 | 20 | 28 | 27 | 28 | 40 | 36 | 57 | 46 | 22 |
| 0.05 | 26 | 28 | 33 | 30 | 30 | 33 | 32 | 37 | 72 | 49 |
| 0.00 | 24 | 19 | 26 | 26 | 32 | 30 | 42 | 33 | 43 | 49 |

##### 6.11 *Linear fixed endpoint control problem*

For the linear fixed endpoint control problem we present $\Sigma = \mathcal{N}_1 + \mathcal{N}_2 + \ldots$ in the tables; the cumulative number of function evaluations required for convergence of the adapted FBSM. With no acceleration, the adapted FBSM applied to the linear fixed endpoint control problem requires solving three two-point boundary value problems (TPBVPs), incurring a total of $\Sigma = 177$ function evaluations. This value is used as the midpoint (yellow) of the heatmaps. Importantly, the acceleration techniques do not reduce the number of TPBVPs that need to be solved, but rather facilitate solving each TPBVP with reduced $\mathcal{N}$, leading to reduced $\Sigma$. For the Wegstein method we apply the same bounds on $q$ as we did for the linear continuous control problem, noting that further tuning of the bounds may improve results.

Wegstein, updating $q$ every $n$th iteration, $-2 \leq q \leq 0$

| $\omega$ | $n$=1 | $n$=2 | $n$=3 | $n$=4 | $n$=5 | $n$=6 | $n$=7 | $n$=8 | $n$=9 | $n$=10 |
|---|---|---|---|---|---|---|---|---|---|---|
| 0 | 91 | 105 | 98 | 95 | 113 | 120 | 111 | 102 | 105 | 126 |

Partial Aitken

| $\omega$ | $m$=1 | $m$=2 | $m$=3 | $m$=4 | $m$=5 | $m$=6 | $m$=7 | $m$=8 | $m$=9 | $m$=10 |
|---|---|---|---|---|---|---|---|---|---|---|
| 0 | 42 | 36 | 36 | 45 | 54 | 84 | 48 | 54 | 60 | 66 |

Partial Steffensen

| $\omega$ | $m$=1 | $m$=2 | $m$=3 | $m$=4 | $m$=5 | $m$=6 | $m$=7 | $m$=8 | $m$=9 | $m$=10 |
|---|---|---|---|---|---|---|---|---|---|---|
| 0 | 173 | 57 | 51 | 33 | 39 | 38 | 27 | 30 | 33 | 36 |

Anderson

| $\omega$ | $M$=1 | $M$=2 | $M$=3 | $M$=4 | $M$=5 | $M$=6 | $M$=7 | $M$=8 | $M$=9 | $M$=10 |
|---|---|---|---|---|---|---|---|---|---|---|
| 0 | 39 | 30 | 30 | 24 | 24 | 24 | 24 | 24 | 24 | 24 |

##### 6.12 *AML fixed endpoint control problem*

For the AML fixed endpoint control problem with no acceleration techniques, the adapted FBSM requires solving ten TPBVPs; incurring $\Sigma = 434$ function evaluations. This is achieved using the best tuning ($\omega = 0.55$) from the AML continuous control problem. In this particular instance, $\omega = 0.55$ also happens to be the best tuning for the AML fixed endpoint control problem if holding $\omega$ constant, when considering $\omega \in [0, 1)$ at increments of 0.05. These $\omega$ values will not necessarily coincide in general, as the adapted FBSM requires solving several related but distinct TPBVPs, each potentially calling for different ideal tuning. For each of the acceleration methods we employ the tuning parameters that minimised $\mathcal{N}$ for the AML continuous control problem. This does not imply that we are using the best tuning parameters for the acceleration methods in the context of the AML fixed endpoint control problem. This is important, as it demonstrates the effectiveness of the techniques in accelerating fixed endpoint control problems without requiring prohibitive tuning. In the following table, we present the cumulative $\mathcal{N}$ after each FBSM within the secant steps of the adapted FBSM. In the right-most column, corresponding to the tenth and final TPBVP, we present $\Sigma$. While Wegstein's method performs significantly worse than the case with no acceleration, we find that the Anderson, Aitken and Steffensen methods are all able to reach convergence in fewer function evaluations.

No acceleration, $\omega = 0.55$

| Secant step | 1 | 2 | 3 | 4 | 5 | 6 | 7 | 8 | 9 | 10 |
|---|---|---|---|---|---|---|---|---|---|---|
| Cumulative $\mathcal{N}$ | 38 | 98 | 147 | 188 | 229 | 270 | 311 | 352 | 393 | 434 |

Wegstein, $\omega = 0.55$, updating $q$ every 6th iteration, $-1 \leq q \leq 1$

| Secant step | 1 | 2 | 3 | 4 | 5 | 6 | 7 | 8 | 9 | 10 |
|---|---|---|---|---|---|---|---|---|---|---|
| Cumulative $\mathcal{N}$ | 26 | 172 | 356 | 423 | 505 | 675 | 809 | 929 | 1045 | 1161 |

Partial Aitken, $\omega = 0.5$, $m = 5$

| Secant step | 1 | 2 | 3 | 4 | 5 | 6 | 7 | 8 | 9 | 10 |
|---|---|---|---|---|---|---|---|---|---|---|
| Cumulative $\mathcal{N}$ | 30 | 72 | 114 | 144 | 180 | 216 | 252 | 288 | 324 | 360 |

Partial Steffensen, $\omega = 0.5$, $m = 5$

| Secant step | 1 | 2 | 3 | 4 | 5 | 6 | 7 | 8 | 9 | 10 |
|---|---|---|---|---|---|---|---|---|---|---|
| Cumulative $\mathcal{N}$ | 19 | 44 | 69 | 88 | 113 | 138 | 163 | 188 | 213 | 238 |

Anderson, $\omega = 0.85$, $M = 6$

| Secant step | 1 | 2 | 3 | 4 | 5 | 6 | 7 | 8 | 9 | 10 |
|---|---|---|---|---|---|---|---|---|---|---|
| Cumulative $\mathcal{N}$ | 17 | 45 | 67 | 86 | 109 | 128 | 147 | 166 | 185 | 204 |
